# Erasure of Biologically Meaningful Signal by Unsupervised scRNAseq Batch-correction Methods

**DOI:** 10.1101/2021.11.15.468733

**Authors:** Scott R Tyler, Ernesto Guccione, Eric E Schadt

**Affiliations:** Department of Oncological Sciences, Tisch Cancer Institute, Icahn School of Medicine at Mount Sinai, New York, NY, USA; Center for OncoGenomics and Innovative Therapeutics (COGIT); Department of Pharmacological Sciences and Mount Sinai Center for Therapeutics Discovery, Icahn School of Medicine at Mount Sinai, New York, NY, USA; Black Family Stem Cell Institute, Icahn School of Medicine at Mount Sinai, New York, NY 10029, USA; Bioinformatics for Next Generation Sequencing (BiNGS) Shared Resource Facility, Icahn School of Medicine at Mount Sinai, New York, NY, USA; Department of Genetics and Genomic Sciences, Icahn School of Medicine at Mount Sinai, New York, NY, USA

## Abstract

Single cell RNAseq (scRNAseq) batches range from technical-replicates to multi-tissue atlases, thus requiring robust batch-correction methods that operate effectively across this spectrum of between-batch similarity. Commonly employed benchmarks quantify *removal* of batch effects and preservation of *within-batch* variation, the preservation of biologically meaningful differences *between* batches has been under-researched. Here, we address these gaps, quantifying batch effects at the level of cluster composition and along overlapping topologies through the introduction of two new measures. We discovered that standard approaches of scRNAseq batch-correction erase cell-type and cell-state variation in real-world biological datasets, single cell gene expression atlases, and *in silico* experiments. We highlight through examples showing that these issues may create the artefactual appearance of external validation/replication of findings. Our results demonstrate that either biological effects, if known, must be balanced between batches (like bulk-techniques), or technical effects that vary between batches must be explicitly modeled to prevent erasure of biological variation by unsupervised batch correction approaches.

---

An ideal batch-correction method removes batch confounded differences in technical effects, while retaining biological signals of interest. With single cell RNA sequencing (scRNAseq) advancing at a rapid pace, a wave of normalization, batch-correction, and benchmarks to assess the accuracy of such methods have stormed onto the scene. Such methods attempt to tackle difficult issues in scRNAseq experiments, such as individual samples often being processed as their own batch, rather than in multiplex. However, our knowledge of unwanted technical variation that can occur between batches is presently limited to count depth per cell^1^, nuclear-long non-coding RNAs (lncRNAs)^2, 3^, and cellular damage^4, 5^. While some unsupervised approaches attempt to use our (limited) understanding of the technical effects (referred to here as Class-1 algorithms)^6–8^, many methods in common use do not explicitly model technical or biological effects (referred to here as Class-2 algorithms), aligning topologies agnostically to the variation source. Class-2 algorithms include CombatSeq-Negative-Binomial, which rotates datasets within count-space, and others that perform a latent dimension mapping (LDM) with a loss function decreasing batch differences: Multiple Canonical Correlation Analysis (MCCA),^9^ Harmony,^10^ Liger,^11^ Scanorama, and scVI^12, 13^. There are also semi-supervised Class-3 algorithms, such as scANVI and scGEN^14, 15^; these attempt to model the biology of interest assigned to a given sample by the experimenter in the form of prior-expectation informed by domain-knowledge. These use a joint learning approach with respect to the human-assigned class labels and the corresponding functional-omic measurements. Here we focus on the efficacy of unsupervised methods (Class-1/2), which cannot be fairly compared to semi-supervised given that semi-supervised methods include prior expectation of “important biology” into the results.

The variations among samples we observe in scRNAseq experiments are a mixture of technical and biological effects. We reasoned that batch correction algorithms that do not explicitly model or discriminate between either type of effect (biological or technical), are at risk of removing both sources of variation (**Supplemental Note-1, Supp. Fig. 1; proof-1**). Because Class-2 algorithms do not explicitly discriminate technical or biological effects, we hypothesized that they may erase biologically meaningful sources of variation in practice. However, even testing this is a challenge, because the ground-truth biology in complex, biologically derived datasets by definition are not known *a priori* suggesting that to discriminate biological and technical effects, a deeper understanding of technical effects impacting batch-to-batch variation is critical.

While others have previously noted the risks of over-mergers^16^ by Class-2 algorithms for the reasons described above, the potential for this property has not been quantified. Furthermore, in practice, a literature mining meta-analysis of Class-2 algorithms indicates their broad use for batch-correction (**Extended Data** (**ED**) **Table 1**); Chi-square test of independence “batch-correction” and <algorithm>; all comparisons: *P*≈0), suggesting that they are none-the-less used with the intention of removing batch confounded technical effects (i.e. batch-correction).

An increasing number of benchmarks attempt to assess batch-correction algorithm efficacy by judging the ability to thoroughly mix batches from different biological samples and technologies (despite these components being intrinsically confounded),^9, 17^; these benchmarks will often assess the ability to re-discover the cluster/cell-type labels defined by those who originally performed the analysis, thus setting their approach to the ground truth^17, 18^. While these approaches are important for cross-technology/species meta-analyses of highly similar datasets, a typical bench biologist performs a treatment/control or disease/non-diseased experiment on a single platform, without *a priori* knowledge of the similarity *between* batches. Furthermore, in practice, the typical bench-biologist will generally follow guidance provided by algorithm authors through their publications/websites when applying the algorithms. In this common use-case, we sought to understand and benchmark the most widely used methods of batch-correction, characterizing how well they can remove technical effects while preserving biological effects of interest.

Here we show that many existing benchmark-metrics either assume biological-equality between separate biological-individuals or assume that published labels accurately identified all meaningful sources of variation. We introduce a new functional framework to aid in parsing technical and biological sources of variation. We then introduce three new methods/tools to extend existing benchmarks for scRNAseq experiments by characterizing local topological differences within-clusters/across batches (WCAB) and within topologically overlapped regions within clusters/across batches (TOWCAB), and by characterizing cluster composition between batches in a way that is invariant to the number of clusters [percent-maximum-difference (PMD)]. We used these metrics to assess the ability of Class 1 and 2 unsupervised algorithms^9–11, 19, 20^ to remove technical differences, and preserve batch-confounded, and non-confounded cell-types/-states both from a cluster-centric perspective (PMD) and along topologically overlapping regions (TOWCAB). We reprocessed several published datasets using different batch correction approaches and found that these issues can dramatically alter interpretation of scRNAseq experiments. First, we examine single- and multi-tissue atlases and demonstrate biological heterogeneity represented in contiguous topologies using the published coordinates, identifying cell-type erasure by two class-2 algorithms. Second, we demonstrate with *in silico* and real-world datasets that transcriptome mixtures (representing cell-states or impure droplets) can result in aberrant co-clustering across batches. Finally, we reanalyzed published human brain scRNAseq datasets, showing that a Class-2 algorithm created the impression of between-dataset concordance, in a manner that has already impacted cancer treatment research, thus raising the stakes of batch-correction, as these approaches move towards clinical care.

## Results

Despite the broad acceptance of cell-identity/-state being fluid and dynamic, many scRNAseq batch-correction benchmarks use manually labeled cluster annotations, assuming that cells from different donors, labeled by different bioinformaticians, are biologically equivalent. As a result of this assumption, such benchmarks penalize a batch correction method for not thoroughly mixing cells of the same label. However, these labels are generated by experimenters doing their best to identify the biological sources of variation that are expected to be important in their experiment. In practice, manually generated labels may not capture *all* aspects of cell-identity, including differences in cell-state (i.e.: circadian state) that may differ between biological samples assayed as separate batches. Common metrics such as ASW, iLISI, cLISI, and kBET-cell-type penalize for maintaining this unlabeled source of variation (**Supplemental Note-1; Supplemental Fig. 2**, **ED Table 2**). The rationale behind these penalties is the presumption that user-defined labels annotate *all* biologically meaningful variation, and within-label topologies should thus be thoroughly mixed, discounting the possibility that the experimenters providing the reference labels may not have provided labels reflecting all biologically derived attributes of cell identity.

We therefore sought to characterize unsupervised batch-correction algorithms in a manner that did not have these limitations or assumptions. Ideally, a batch-correction algorithm should remove batch confounded differences in technical effects while preserving biological effects. Given that we seek to study biological effects and cannot presume to know them *a priori*, we began from first principles, asking what types of functions are involved in defining the signals detected from various high-dimensional molecular measures (e.g., functional genomic data) and how might they be confounded or balanced between batches.

We began from the premise that all observations in a scRNAseq dataset are manifest from either biological (B) or technical (T) effects (**Fig. 1**; **Supplemental Note-2**). Prior to experimenter intervention to perform the measurements, only biological effects exist (**Fig. 1a**). Once sample collection has begun technical effects begin to accumulate, the technical effects will be generated through both the biological response to sample processing (T_r_), which is not of interest and can be reduced by inhibition of transcription/translation^21, 22^, technical effects of sample processing/measurement (T_m_) (different lysis buffer composition, sequencing depth, etc) (**Fig. 1b**). In the context of technical-replicates, we know that the biology of interest (B) is the same between the two batches, leaving only the differences in technical effects of measurement and processing (T_m_), and the cell’s response to processing (T_r_) (which has been shown to be cell-type specific^22^). When treated as a ‘pattern matching’ problem however, we are at risk of removing biology of interest, when these sets of functions are not explicitly distinguished, if there is any batch confounded biology, including in biological replicates (**Fig. 1c**, **Supplemental Note-3**).

**Figure 1:**
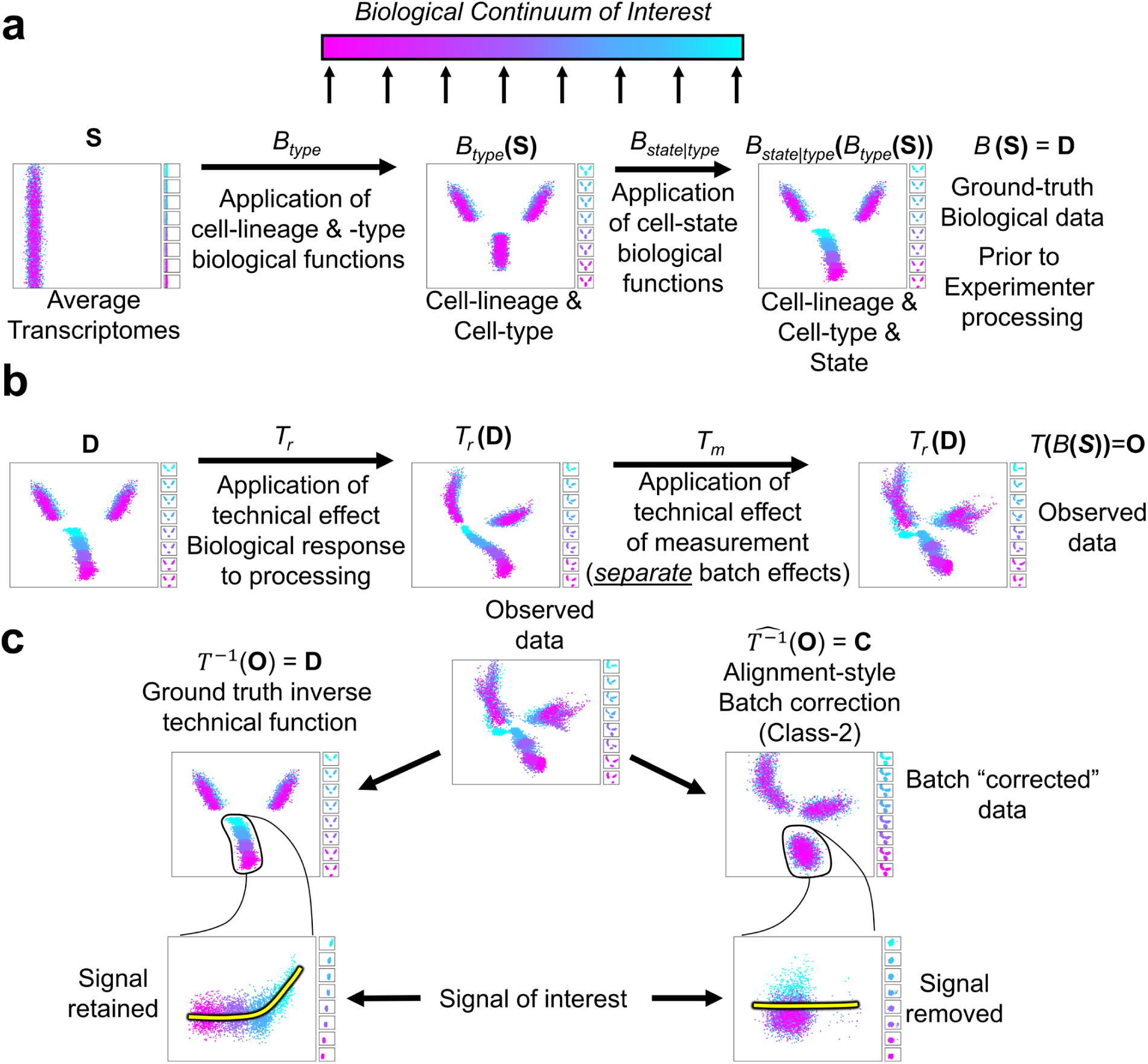
Functional framework for aiding experimental design in single-cell assays. A 2-dimensional representation of cells to illustrate the functional framework we devised that describe the transformations that cells (and their -omic measures) take. **a**, Biological (B) transformations are encoded in sets that are applied to the cells, beginning with the biological functions applied to an average transcriptome (**S**) that encode transformations related to cell-lineage/-type (B_type_). Next, in an experimental design or disease process, various changes in cell-state can exist encoded by functions contained in (B_state_). **b**, Once an experimenter begins to intervein to perform the experiment, technical effects occur, as cells react to processing through biologically derived processes (T_r_), and bias/errors batch effects are introduced when samples are processed separately (T_m_) (for example with different master-mixes, causing differing amplification efficiency, etc). This generates the observed data matrix (**O**). **c**, Next, batch correction algorithms are applied, in an attempt to find the ground truth inverse function of technical effect functions (*T^−1^*). However, only a hypothesized approximation of this function can be made 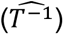; when 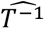 does not model whether the applied inverse functions are from the biological set (B), or technical set (T), we cannot be certain the 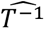 does not contain inverse functions from the biological set.

This type of framework captures the fact that with true technical replicates, local differential expression analysis could measure the impact of technical effects directly (**Fig 2a**, **Supplemental Note-2**; **Supplemental Fig. 3**). We thus developed an open-source tool to assess differential gene detection (rather than expression, given that true technical replicates have no true differences in transcription prior to processing) within-clusters/across-batches (WCAB)^23^, and then used protein-protein interaction and pathway analysis to identify the biological coherence of the differential transcriptomic signatures, using local backgrounds of only what was detected within each cluster to prevent generic tissue enrichment, high expression, or annotation bias^24, 25^. Surprisingly, in technical-replicates derived from separate tissues (mouse-brain^26^ and mouse-heart^27^), we observed consistent differential detection signatures regardless of correction strategy (**Fig. 2b**), that were enriched for large structural components of the cell, such as mitochondria, the endoplasmic reticulum (ER), nuclear targeted genes, and separately hemoglobin (**Fig. 2c**). These gene-complexes (T_m_; **ED Table 3**) were differentially detected within-clusters/across-batches in a large proportion of clusters, and conserved across tissue types (**Fig. 2d-g**). However, the cell-stress and iron response (T_r_) was highly divergent across tissue-types in comparison to T_m_ genes/pathways (T=10.69, P=1.24e-4, 1-sample T-test) (**Fig. 2d**), fitting with prior reports of cell-type specific responses to processing^21, 22^. We observed these same complexes were differentially detected in a different species and tissue-type (human PBMCs) (**Fig. 2e-g**). The more conservative intersection of mouse technical replicates (or their human orthologues) for this classification system accounted for 77.3%, 69.2%, and 57.3% of differentially detected genes between technical replicates for mouse-brain/-heart, and human-PMBCs, respectively.

**Figure 2:**
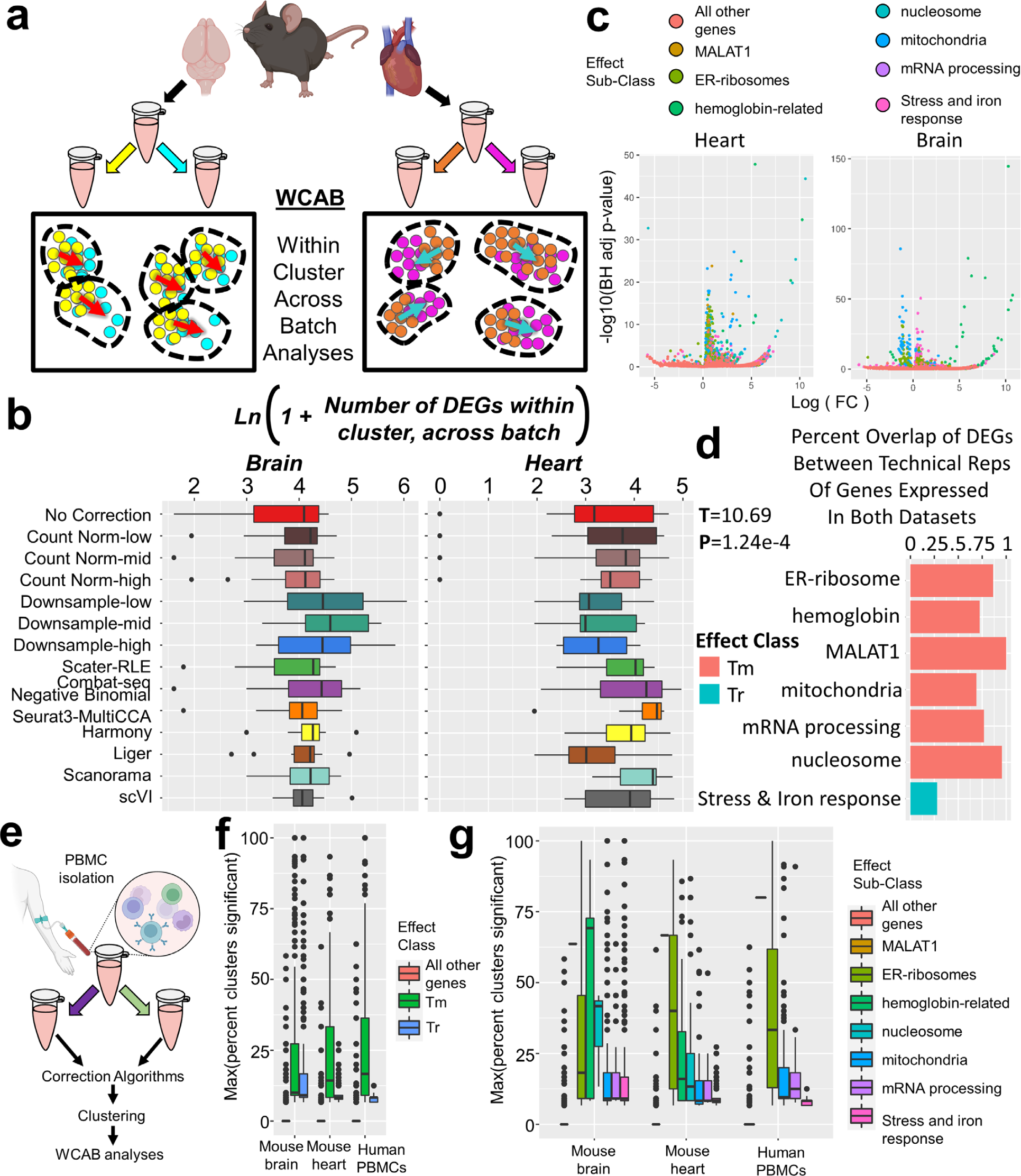
Locally differentially expressed genes across technical-replicates. **a**, Using brain and heart technical-replicates, batch-corrections were performed followed by differential expression within-clusters/across-batches (WCAB). **b**, Broad differential expression within clusters was seen regardless of correction method is shown by boxplots quantifying the log number of differentially expressed genes within clusters across batches. **c**, Differentially expressed genes were largely those localized to large structural components of the cell including mitochondria, ER, and nucleus. Another class of genes fell into the cell stress response. **d**, The conserved differentially expressed complexes are identified as the technical effects of measurement (T_m_), while the technical response (T_r_) to measurement is less conserved across datasets. **e**, Human PBMC technical-replicates were assayed by WCAB analyses. **f**, Boxplots showing that T_m_ effects uniformly impacted a higher percentage of clusters than T_r_, which was more cluster specific. **g**, Boxplots showing the percentage of clusters in which a gene was differentially expressed in the noted dataset; displayed is the maximum of correction methods for each gene (See **ED Table 3** for all results).

### Efficacy of batch-correction algorithms with identical cells and technical-replicates

While technical replicates capture the combined impact of batch effects, such effects may still contain several sources of technical variation. We therefore began our benchmark specifically quantifying the ability of algorithms to remove the effect of count depth, quantifying the number of sub-sampled cells that are co-clustered with their original cells, an approach recently dubbed “molecular-cross-validation”^28^. Class-2 methods performed marginally better than no correction or Class-1 algorithms when only analyzing mouse brain, however, this advantage was no longer present when analyzed in the presence of a third dataset for which we expected little overlap in cluster composition (homeostatic intestine) (**Box 1b**; **ED Fig. 1**).

Many scRNAseq analyses build on clustering results; it is therefore important to quantify the effect of batch-correction on clustering results. However, batch-correction algorithms may leave differing levels of variation in the final dataset, resulting in differing numbers of clusters; yet there is generally no ground-truth number of clusters in biological datasets. We therefore created the “percent-maximum-difference” (PMD) (**Fig. 3a**) metric/test to quantify the overall similarity in cluster composition across batches. Unlike X^2^, X^2^ -log10(p-value), Cramer’s V, inverse Simpson’s Index, or count normalized Shannon Entropy across clusters, PMD is provably invariant to the number of clusters found when relative overlap in cluster composition is preserved, operates linearly across the spectrum of batch similarity, is unaffected by batch size differences or overall number of cells, and does not require that batches be similar (**ED Fig. 2**; see **Methods** and **Supplemental Note-4** for proof, benchmark, and detailed characterization of PMD; **Supplemental Figs. 7, 8**).^29^ PMD yields a single quantitative representation on a scale of 0-1 ranging from highly similar batches to highly dissimilar batches. A PMD=1 indicates batches with no overlapping clusters (highly dissimilar), while PMD≈0 indicates highly similar batches, say, from the same single cell suspension (see **ED Fig. 2b,c** for a visual representation and calculation details).

**Figure 3:**
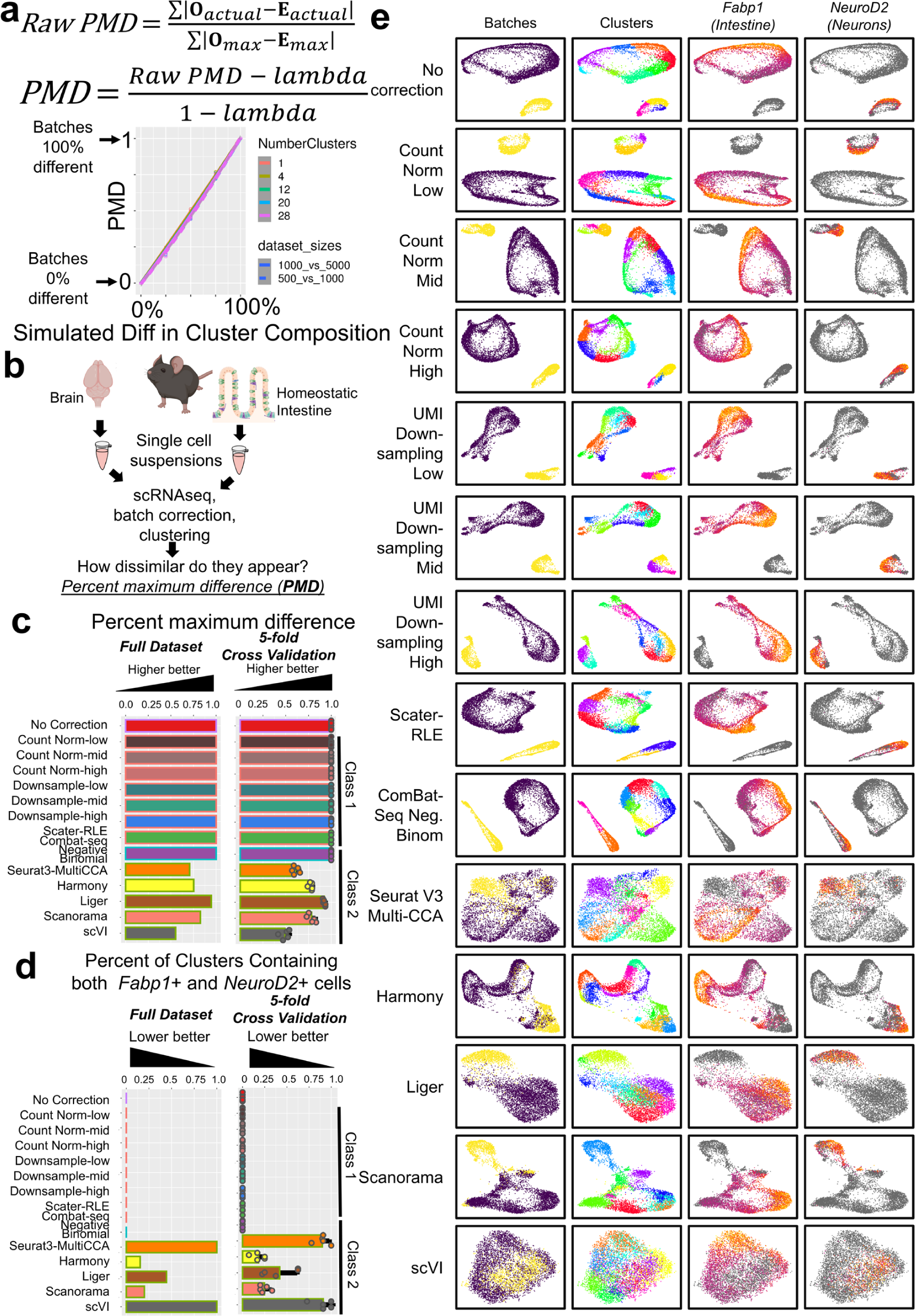
Batch-correction algorithm performance when samples are expected to have no overlap. **a**, The PMD equation, to compute relative difference between batches based on their cluster composition, and a scatterplot from simulations (Supplemental Note-4; Supplemental Figs. 7,8), showing linearity and robustness to differences in number of clusters or dataset sizes. **b**, A schematic representation of the dataset integration task in which there is little to no overlap in cell-types expected (brain and intestine samples) quantified by the percent maximum difference (PMD). **c**, A bar chart displaying the PMD between the input brain and intestine datasets. Five-fold cross-validation of this integration task show highly concordant results. **d**, A bar chart showing the percentage of clusters containing both *FABP1*+ and *NEUROD2*+ cells (markers of intestinal epithelia and neurons respectively) at ≥5 UMI in original datasets; all bar charts show means+/−SD. **e**, Dataset mixing visualized by a spring-embedding plots of the kNNs used for clustering, colorized by batch, cluster, and marker-gene positivity.

Using PMD, we quantified batch similarity after correction, when cells came from the same single cell suspension (technical-replicates), where the PMD is expected to be low or near-zero (**Box 1c:** using technical replicates from neurons, heart, and PBMCs as model systems)^30, 31^. Most algorithms (and no-correction) performed quite well at the cluster level by PMD (**ED Fig. 3a-c**). The results were similar across a range of tissues, including mouse brain, mouse heart, and human peripheral blood mononuclear cells (PBMCs) (**ED Fig. 3d,e**), with none of the batch correction methods giving rise to a PMD greater than 0.04.

### Efficacy with dissimilar biologic datasets

We next performed negative controls (no expected cluster overlap) by comparing brain^31^ and intestinal epithelium^32^, given that the only immune cells recovered after epithelial and crypt enrichment were T-cells, and the brain dataset contained no detected CD3E or CD3G; however, this is still an expectation of dissimilarity based on biological knowledge (**Box 1d**; **Fig. 3b**; expected PMD=1). While Class-1 algorithms indeed indicated no overlap in cluster composition (PMD=1), Class-2 algorithms with latent-dimension-mapping (LDM) had PMD scores up to 0.4, which corresponds to 60% relative similarity in cell type composition between the brain and intestine (**Fig. 3c**). Class-2+LDM methods indeed harbored mixed neuron/intestinal epithelial clusters (>30% of clusters mixed in full datasets), while Class-1 approaches did not (**Fig. 3d,e**).

### Results with biologic datasets of uncertain overlap

In contrast to the above positive- and negative-controls, batches will typically give rise to clusters of cells that have unknown overlap, and even cells of the same type may differ by cell-state across batches. We therefore analyzed homeostatic intestine and irradiated intestine (**Box 1e**; **Fig. 4a**)^32^, expecting some overlap given both tissue types began with the same base cell-types, but then given cellular damage caused by radiation exposure, we would also expect dramatically different cell-states, replication, and repair (**Fig. 4b**). Results ranged from PMD values of 0.99 with no-correction, to PMD values of 0.20 (Harmony) (**Fig. 4c**). These results demonstrate that the choice of correction method can drastically alter the biologic conclusions of an experiment with batches of divergent cell-states.

**Figure 4:**
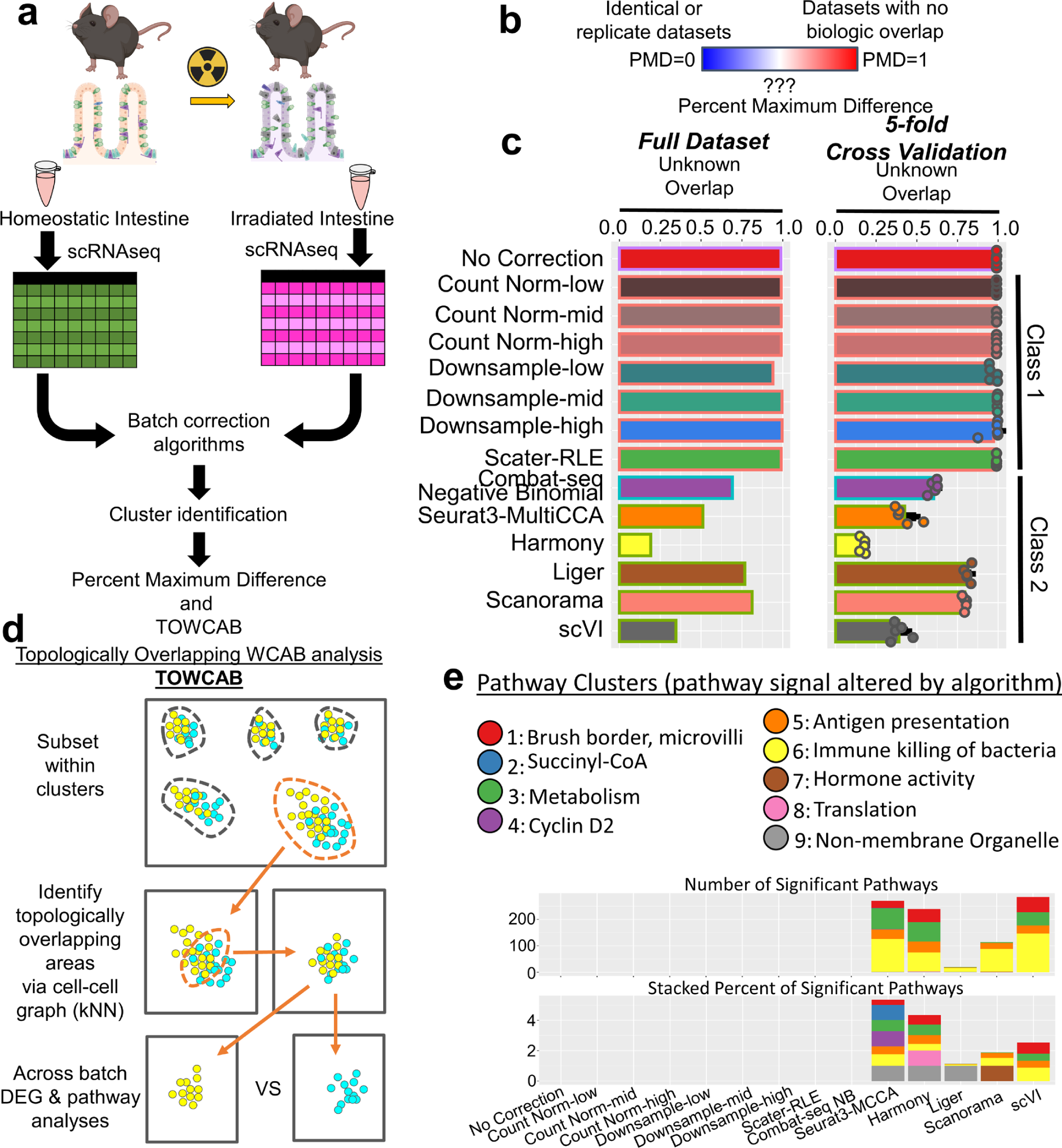
Class-2 algorithms with LDM can create biologically heterogenous topologies. **a**, Schematic of analysis approach: Homeostatic and irradiated mouse intestines were analyzed, and assessed both by PMD, and a refinement of WCAB, that selects only topologically overlapping areas within clusters (TOWCAB)^32^. **b**, There was no clear *a priori* expectation how similar these datasets would be by PMD. **c**, A wide range of PMD values was observed ranging from nearly completely different without batch-correction and with depth normalization approaches to mostly the same (20.2% maximum difference) with Harmony. Five-fold cross-validation of this integration task show highly concordant results (means +/− SD). **d**, Quantification of the number of, or stacked percentage of, significant pathways whose signal are altered along topologically overlapping regions.

To quantify the biological pathways that were subject to erasure by some batch-correction approaches, we refined our WCAB approach to filter each cluster to include *only* topologically overlapping regions of the pairwise batch-batch comparisons (**Fig. 4d**). We performed this procedure because topologically overlapping regions are often interpreted as containing biologically similar cells. This is in contrast to another possibility in which two topological locations are “zippered” together or placed adjacent by a batch-correction algorithm in a manner that creates an apparently contiguous object in similarity space, but where the objects are better described as two conjoined, but non-overlapping objects. To identify and distinguish these types of situations, we developed the open-source Topologically Overlapping Within-Cluster/Across-Batch (TOWCAB) method (**ED Fig. 4**).

We applied the TOWCAB method to the homeostatic/irradiated intestine data to directly quantify which attributes of cell identity were removed by the different batch correction algorithms. We identified 9 clusters of pathways that were differentially expressed within topologically overlapping areas produced by Class-2+LDM algorithms (**Fig. 4e**; **ED Fig. 5a**). These included immunologic pathways related to antigen presentation, response to pathogens, reactive oxygen species, and other processes that would be expected to be biologically important during irradiation mediated disruption of the intestinal barrier (**Fig. 4e**). Importantly, these pathways were pre-filtered to exclude any pathway that was observed as significantly different in any of the technical replicates (**Fig. 2**), thus quantifying only pathways that had no prior evidence of being impacted by technical sources of variation.

**Figure 5:**
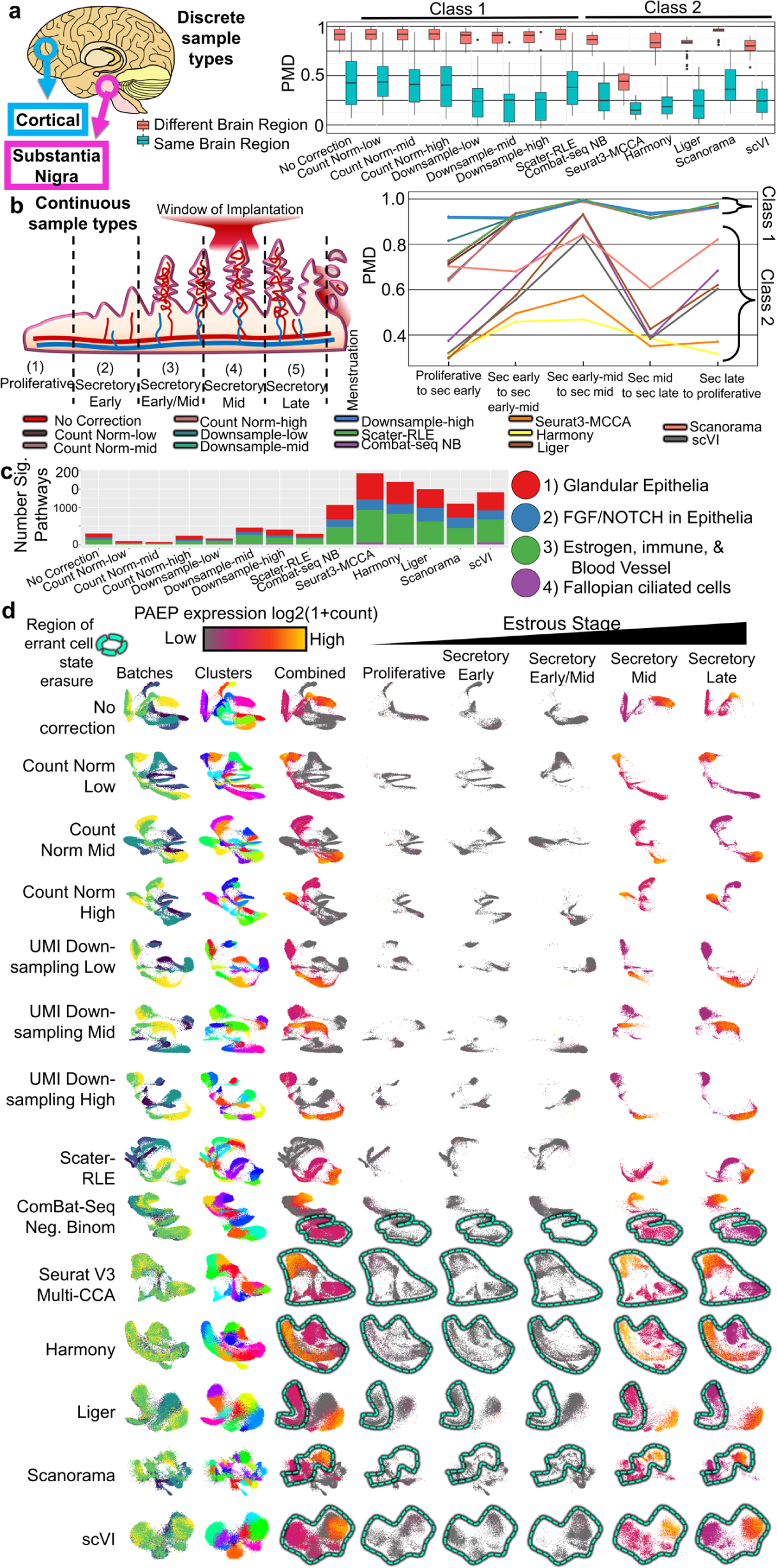
Batch-correction of discrete or continuous sample types. **a**, Boxplots showing measured PMD values for within and between tissue-types on batch-correction approaches using on two human brain sample-types (cerebral cortex (n=5) and substantia nigra (n=7)) that share all cell-types, but appear in differing abundance. **b**, Continuous variation in cell-abudnace and state within the endometrium over estrous was assayed for batch correction methods, displaying line-graphs of apparent batch-batch similarity over the estrous cycle (**g**:x-axis=time; y-axis =PMD, color=correction method). **c**, TOWCAB analyses show that Class-2 batch-correction methods erase a greater amount of biologically meaningful pathways. **d**, Plotting expression of *PAEP* (a known cell-state gene, up in mid-secretory) illustrates this cell-state erasure by Class-2 methods standard methods.

### Discrete sample types vs continuously divergent sample types

A remaining question is the impact of integrating samples with many replicates that originate from discrete sample-types or those that exist across a continuum of changing cell-states and cell-type abundances. To address the former, we integrated cortical and substantia nigra samples, each with biological replicates^33^. Pairwise sample-sample PMDs were calculated and displayed for within vs between region similarity (**Fig. 5a**); this show a clear separation in which samples from within a region (cortical from substantia nigra) were more similar relative to across region comparisons for all methods (including no-correction), with the exception of Seurat’s MCCA (Class-2).

For clusters to ‘split’, however, a large effect must be present, meaning that the topologies may be slightly askew across batches, but not sufficiently distinct to induce cluster splitting. We assessed topological heterogeneity, via TOWCAB, revealing that Class-2 algorithms topologies were more heterogeneous than Class-1 (**ED. Fig. 5b**). These altered pathways included those related to neurons, synapses, dendrites, and ion transport, indicating that it was variation in these pathways that were erased in order to produce a “corrected” topology.

In contrast to the above (differential cell type abundance between two regions of the same tissue), another realistic scenario is the presence of dynamic continuous processes across batches. We therefore analyzed the endometrium across the menstruation cycle which has previously described changes in cell-states and cell-type abundances, particularly at the window of implantation^34^. Class-1 approaches were consistent with each other identifying these samples as largely dissimilar, with changing cell states and cell-type abundances. On the other hand, Class-2 algorithms increased sample-sample similarity (lower PMD) (**Fig. 5b**). TOWCAB analyses found that Class-2 algorithms erased pathways that appeared biologically pertinent, including angiogenesis, microvillus related structures, immune cells, and endometrial cells (all well described aspects of the estrous cycle^35, 36^; **Fig. 5c**). Notably, variation of the Progestagen Associated Endometrial Protein gene (*PAEP*; or Glycodelin-A), (which is dynamically regulated throughout estrous^37^) is topologically erased by Class-2 algorithms along the axis of this cell-state (**Fig. 5d**).

It is again important to note that these pathway analyses each used custom backgrounds of locally expressed genes within clusters ensuring that this was not generic enrichment for brain or endometrium related pathways relative to the whole genome, but were rather locally enriched among the genes expressed in topologically overlapping regions of the datasets, taking into account what *could have been* detected. Further, any pathways that were previously shown to be differentially detected in our technical replicate studies were excluded for this analysis, as removal of variance along these dimensions should not be penalized, as this is the goal of batch-correction.

### Erasure of cell-type signal within an atlas

To differentiate the effects of our specific implementations of these algorithms from what is seen in practice, we performed TOWCAB analyses on a recent release of human urinary bladder cells (30,942 cells; 10 donors, aligned using Seurat’s MCCA), based on their released UMAP coordinates^38^. Indeed, we found that different cell-types were merged across batches including B- and urothelial-cells (**ED Fig. 6**). This further highlights both the efficacy of TOWCAB analysis in the discovery of biologically heterogeneous regions of the topology, and also highlights that we cannot always rely upon cell-type labels from atlases as “ground-truth” labels for benchmarking. We similarly observed what appeared to be B-cell-related droplets, that were labeled as satellite stem cells, from a muscle sample, that were topologically placed within an otherwise fibroblast related cluster within the *tabula sapiens* resource (based on stromal UMAP coordinates following scVI correction; see **Supplemental Note-1, Supplemental Fig. 1**)^39^.

**Figure 6:**
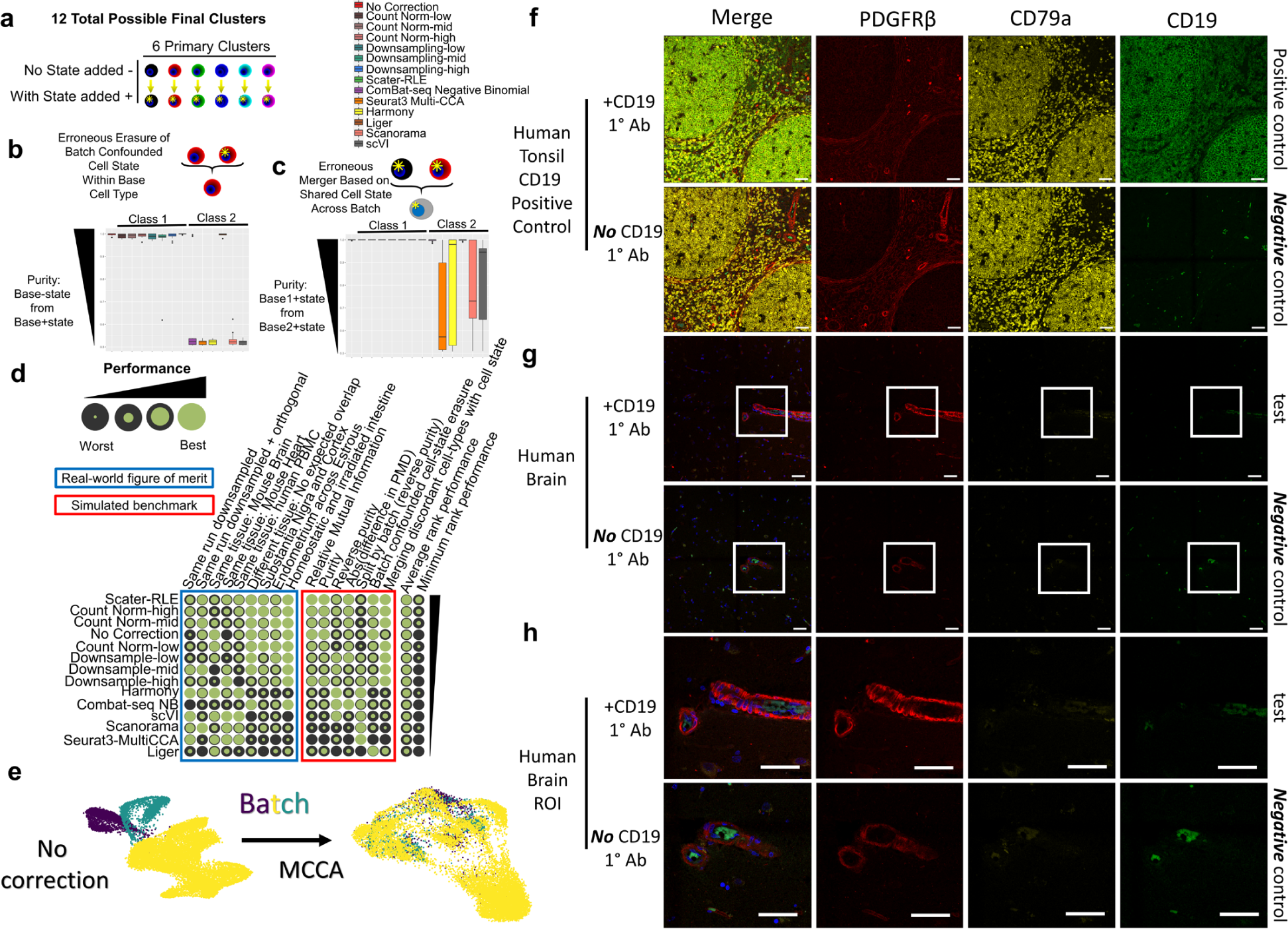
Class-2 algorithms over-merge when transcriptome mixtures are present. **a**, Twelve possible clusters were used in simulations; they were generated from six primary clusters, with a cell-state optionally added (**Supplementary Note-5**; **Supplementary Fig. 9**). **b,c,** Boxplots and schematics illustrating the possible failure-mode for erasure of “cell-state” or “cell-type” sources of variation when batch confounded. Boxplots are the typical display of median, with the box highlighting the second-to-third quartiles and whiskers extending out to the first and fourth quartiles; outliers are defined as 1.5x the inner quartile range. **d**, Each tested method (row) is sorted by their average rank in real-world (blue box) and synthetic (red box) assessments. **e**, After batch-correction of the three human brain datasets, MCCA (as was performed in the original manuscript) shows thorough mixing of batches. **f**, We performed CD19 positive control staining on human tonsil B-cells along with PDGFRβ and CD79a including a no CD19 primary antibody negative control. **g-h**, With the same exact staining and imaging conditions as our positive control, we stained 4 donor brains including the no CD19 primary negative control. The small signal in the CD19 channel was luminal rather than perivascular, was also faintly positive in the CD79a channel and present in the no-CD19 primary antibody condition, consistent with background autofluorescence. We did not observe CD19/PDGFRβ co-positive perivascular cells. Scale bars indicate 50um.

### Synthetic dataset benchmarking with simulated cell-types/-states

Given these examples in which Class-2 algorithms appear to erase biological signal of interest, we turned to simulations to further demonstrate this issue, where through the simulations complete control of the ground-truth cell-lineage -type and/or -state can be achieved. To this end, we simulated cell-states *de novo*, *in silico*, with a linear-mixed model of negative-binomial distributions, coarsely representing a possible cell-type/-state mixture under various batch confounded and non-confounded circumstances, comparing known ground-truth results to results downstream of the different batch correction algorithms. Using this approach, with a known ground truth (rather than reliance upon consistency with previously published labels), we found that Class-2 algorithms largely erased batch confounded mixed expression programs (**Supplemental Note-5**; **Fig. 6a-c**). These results mirror our biological findings that Class-2 approaches may over-merge discrete cellular populations, which should be either split by cell-type (**Fig. 3,5**, **ED Fig. 5**) or cell-state (**Fig. 4**, **ED Fig. 5**).

By combining the results of our molecular cross-validation (higher percent-same), technical replicates (lower PMD), negative control integrations (higher PMD), more complex integrations (lower TOWCAB significant pathways), and simulations (closer to ground-truth), we found that Scater’s relative-log-expression (RLE) ranked the highest of methods assessed in this work (**Fig. 6d**).

### Real world implications of mixed-model based over-mergers by Class-2 algorithms

Given that we had found *in silico* linear mixture models create shared sources of variation that enable over-mergers by Class-2 algorithms, we hypothesized that *biophysical* linear-mixtures (doublets or cell-fragment mixtures in a droplet^40, 41^) may act as “seeds” for over-integration, with contaminated droplets masquerading as a “novel cell-type” with unexpected marker gene positivity. Given our above findings, application of Class-2 integrations may result in apparent topologic-replication of these results (biologically heterogeneous cells intermingled topologies containing mixed droplets).

For example, CD19 (a canonical B-cell marker) has recently been described as expressed in a subset of human brain mural cells taking this as evidence that an anti-CD19 CAR-T cell therapy would also kill these brain cells^42^. This finding appeared to be confirmed following a Class-2 batch-correction (Seurat’s MCCA) with other datasets, after which they observed co-clustering which was interpreted as a repeated discovery of this population^42^.

We performed in-depth analysis of these datasets including examination of 1) total counts, 2) number of observed genes, 3) creation of linear mixed models of “pure” B-cell and “pure” mural cells comparing these models to the global transcriptome of the candidate CD19+/mural-droplets, 4) transcriptome-wide correlations with CD19, and 5) expression of marker genes (See **Supplemental Note-6**; **Supplemental Fig. 11**). All of these analyses were consistent with a model that the originally observed CD19+/mural-marker co-positive droplets were simply a mixture of transcripts from B- and mural-cell lineages. Indeed, in the original dataset nearly all droplets were contaminated with transcripts from multiple lineages (**Supplemental Note-6**). Lastly, we analyzed a recently released atlas of fresh human brain vasculature and associated cells; we observed only 1 out of 16,681 droplets that were positive for both CD19 and mural cell markers, while also being negative for other B-cell markers (**Supplemental Fig. 11**)^43^. We hypothesized that the appearance of independent replication through co-clustering following MCCA (**Fig. 6e**) may have been an over-merger that may not replicate at the bench.

While peroxidase based anti-CD19 immuno-histochemistry in the original report showed peroxidase signal in some cells near or in vasculature, there was no co-stain with B- or mural-cell markers. To test whether CD19 is in fact expressed in mural cells, we co-stained for CD19, CD79a (pan-B-cells), and PDGFRβ (mural cells) in four human brains using no-primary negative controls to set thresholds for CD19 positivity. We did not observe cells that appeared positive for CD19 and PDGFRβ that were also negative for CD79a, despite CD19 staining positive controls working well (human tonsil) (**Fig. 6f-h**). Notably however, a negative finding can never be definitive; given the exceptional clinical importance of CD19 expression in off-targets in the brain for anti-CD19 CAR-T-cell therapies, these findings should be further interrogated.

Overall, our findings confirm the necessity of checking the assumption that topologically mixed cells are biologically similar after Class-2 alignments, through methods such as TOWCAB, or Class-1 algoirthms my be applied, with the best performing method by our benchmark being Scater’s implementation of relative-log-expression (RLE)^44, 45^.

## Discussion

The field of scRNAseq has been evolving rapidly, both at the bench and computationally. Here we have shown that the most commonly used metrics for benchmarking methods of batch-correction often rely on either 1) the assumption of biological equality across different samples, or 2) the assumption that humans who performed the analysis to generate the reference labels, were successful in optimally finding and labeling all meaningful attributes of biological variation. The methods that rely on the second assumption reward for *similarity* to the methods used by the original experimenters rather than accuracy. While all methods are subject to biases and losses of accuracy given deviations from underlying assumptions realized in practice, understanding the sensitivity of a method to such deviations is critical in achieving accurate interpretations of method outputs. Here we have attempted to highlight a range of issues that can potentially arise in seeking accurate interpretations from single cell RNAseq data when assumptions for different batch correction algorithms are unknowingly violated.

Given the premise that scientific measurements of biologic systems are manifest from the biological- and technical-functions, where, importantly, the biological-functions are what we aim to *measure*, a more complete understanding of the technical functions at play in complex experiments such as scRNAseq, is needed to disentangle them (either computationally or experimentally) when batch and biological variation are intrinsically confounded. To this end, we described a functional framework to help formalize the issues, which ultimately led to our identification of differentially detected transcripts encoding proteins that appear to localize to large structural components of the cell, (and hemoglobin when RBCs have not been removed), as strong contributors to the technical effects.

We also fill a gap in our ability to accurately quantify batch overlap based on cluster composition, which required our development of the PMD measure: a measure that is provably invariant to the number of clusters (of course under its own set of assumptions), is linear with respect to similarity, and robust to dataset size or differences in batch sizes (**Supplementary Fig. 7,8**). These properties, for the first time, enabled a quantitative assessment of normalization/batch-correction algorithms across a full spectrum of similarities based on cluster composition without requiring the assumption of biological equality between biological replicates, or that near perfect annotation of the underlying biology was achieved by the experimenters.

From the perspective of topologies, we introduced (TO)WCAB analyses that quantify differentially expressed/detected genes and pathways, specifically within topologically intermingled areas across batches. Our results suggest that when using unsupervised batch-correction algorithms, TOWCAB-like analyses should be performed and reported so that experimenters, reviewers, and readers can understand which attributes of gene expression were removed by the algorithm. Such understanding can protect against misinterpretation that adjacent cells within a topologic embedding are biologically similar, when they are in fact different. Our results also motivate exercising further caution against interpreting topological concordance as an indication of similarity after Class-2 batch-correction, especially when comparing treatments, genotypes, or other condition groups. Topological concordance as an indication of similarity has has become increasingly common practice ^42, 46^. Similarly, WCAB can be employed when interpreting co-clustering after Class-2 algorithm application.

Importantly however, Class-2 approaches remain the only viable option when integrating across sequencing technologies or across species. Despite this, our results demonstrate that these algorithms cannot be blindly employed without checking for biological heterogeneity across apparently contiguous topologies. We also provided new tools and approaches, enabling the means to check these assumptions (PMD, WCAB, and TOWCAB). Lastly, we show that these are not theoretical concerns, but may have already had direct impact on atlases and research into clinical treatments for cancer. We further hope that the community will build technical replicate atlases to enable explicit modeling and removal of technical effects in single cell -omics.

## Acknowledgements

Support for this work was provided by T32CA078207 (SRT), K99HG011270 (SRT), U01 AG046170 (EES), and RC2 DK122532 (EES).

## Author Contributions

SRT conceived of and performed all analysis, bench-work, and wrote manuscript. EG provided feedback and edited figures. EES wrote Proofs 1 and 2, and guided research and edited manuscript.

## Methods

### Biologic datasets: 10x Genomics datasets

All datasets, baring those of intestinal origin were obtained from Chromium’s publicly available datasets.

PBMCs:

(https://cf.10xgenomics.com/samples/cell-vdj/3.0.0/vdj_v1_mm_c57bl6_pbmc_5gex/vdj_v1_mm_c57bl6_pbmc_5gex_filtered_feature_bc_matrix.h5)^47^

1k neurons:

(https://cf.10xgenomics.com/samples/cell-exp/3.0.0/neuron_1k_v3/neuron_1k_v3_filtered_feature_bc_matrix.h5)^31^

10k neurons:

(https://cf.10xgenomics.com/samples/cell-exp/3.0.0/neuron_10k_v3/neuron_10k_v3_filtered_feature_bc_matrix.h5)^30^

### Other biologic datasets

The intestinal datasets, which were aligned to the Ensembl reference, as with those from Chromium, were previously published.^32^ Several datasets from the human cortical and substantia nigra^33^ and the endometrium were also previously released^34^. However, all raw input datasets used here are re-distributed in the benchmarking repository for ease of replication. Human brain datasets are distributed as an Rdata file within the data/cd19_mural_cell_data directory of the repository. Any other public datasets are downloaded in-line within the code in the repository.

### Class-2 algorithm literature mining for co-mentions with the term “batch-correction”

Searches on google scholar were performed to quantify the relative frequency at which Class-2 algorithms were co-mentioned with the term batch-correction. Searches were performed on July 24^th^, 2022. The number of results containing “single cell RNAseq” were used as the baseline “total possible” results, then search terms were added to this base including with and without the term “batch-correction” and with each Class-2 method included in this manuscript ultimately to tabulate a contingency table for each method within the subset of single cell results that either mention or don’t mention the method at hand and along the other contingency table axis either mention or don’t mention the term “batch-correction.” Results are contained in **ED Table 1**.

### Global dataset downsampling

For the benchmarks that included sub-sampled “child cells,” this 50% transcript sub-sampling of the dataset was created using the pyminer_norm package (available by pip: python3 -m pip install bio-pyminer-norm) using the command line call:

> python3 -m pyminer_norm.random_downsample_whole_dataset -i <dataset.tsv> -o

> <dataset_ds50.tsv> -percent 0.5

### Implementation of normalization approaches

#### No normalization

For no normalization, the datasets are simply concatenated by genes with no other modifications.

#### UMI downsampling

We created a new python package that is pip installable called bio_pyminer_norm to provide a parallelized and scalable implementation of UMI downsampling. This approach requires the user to determine lower (and optionally upper) bounds of the sum of the UMI counts and total number of observed genes for which cells will be included or discarded. As implemented here, we used 3 cutoffs to show the impact of changing sensitivity with this parameter. Cutoffs were as follows for all biological datasets: low:10^3.55^, mid: 10^3.6^, high: 10^3.65^ all with a minimum number of observed features of 10^3.1^; for simulated datasets (which were deeper), cutoffs were: low:10^3.65^, mid: 10^3.7^, high: 10^3.75^ all with a minimum number of observed features of 10^3.15^. Next, each cell is bootstrap-sampled, having its observed transcriptome simulated, shuffled, and an equal number of transcripts randomly selected. This is downsampled to the lower cutoff value for total sum UMI.

#### Count Normalization

Cells/droplets were normalized as counts per x in which x is an arbitrary scalar. In our implementation, we use three different cutoffs, which were the same as implemented for downsampling.

#### Scater-RLE

Following guidance from the Scater repository, we implemented this normalization approach using the following call: scater::normalize(exprs, exprs_values = “counts”, return_log = TRUE, log_exprs_offset = NULL, centre_size_factors = TRUE, preserve_zeroes = FALSE).^19^

#### Combat-seq Negative Binomial

Combat-seq with Negative binomial distribution is distributed in the sva-devel package. Following author suggested tutorial, we used the following implementation: ComBat_seq(exprs, as.character(batch), group=NULL, full_mod=FALSE)^20^.

#### Seurat-v3 Multi-CCA

Following the SeuratV3 website tutorial on integration, we first normalized and processed each dataset using: NormalizeData then FindVariableFeatures. Next, we used the functions: FindIntegrationAnchors and IntegrateData to complete the MCCA data integration in SeuratV3; as recommended, clustering was performed on the first 30 PCs^9^.

#### Harmony

To use Harmony, we followed the author guidelines available on the repository website, first normalizing counts to counts per thousand, then log2 transforming the matrix, next genes were min-max scaled to range between 0 and 1. We used the following call: HarmonyMatrix(exprs, batch_table, “batch”, do_pca = TRUE)^10^.

#### Liger

Following the Liger repository tutorial, we used the createLiger function on the input list of datasets to be normalized; next we utilized the liger::normalize function, followed by the selectGenes, and scaleNotCenter functions.

As per author guidance on their repository tutorial, we used the Liger suggestK function to generate a K vs median KL divergence curve. They authors suggest using the ‘elbow-rule’ of this curve, however, given that there is no actual relationship between the units of number of clusters and KL-divergence, we implement a K selection method that sums the area under that curve (AUC), and select the K that first reaches 95% of this AUC. This K was used for the optimizeALS function. Lastly, we used the quantileAlignSNF function to complete Liger based batch-correction.^11^

Given that Liger only yields a low-dimensional representation of the data, no additional feature selection was performed and we used this representation for clustering directly.

### WCAB differential expression

To determine differential expression within clusters, we used the recently released Libra package with pseudo-bulking and pseudo-replicates^23^. We used three pseudo-replicates for each batch, randomly partitioning them to contain at least 3 cells per pseudo-replicate, giving a minimum requirement of 9 cells per batch to be able to perform DE analysis for each pairwise batch comparison. The default EdgeR flavor of DE analysis was used. For comparisons with more than two batches, a gene was considered differentially expressed if it was globally significant by Benjamini-Hochberg FDR *q*<0.05 for any pairwise comparison.

### Topological filtering for TOWCAB analysis

The topologies used for filtering were the same graphs used for Louvain clustering in all cases, except for when topologies were built from published coordinates; this is described in its own section.

For each batch-batch comparison, each cluster was subset. Within each cluster, each possible batch-batch pair is compared; within this subset, cells were quantified to assess the relative proportion of edges shared with droplets of their “host” batch, or the other batch it is being compared to. For each cell, 100 bootstrap shuffled edges were used to generate a cell-wise background to empirically measure the host-batch proportion with fully entropic mixing would yield for that given cell. Cells were kept that were within 1.5 standard deviations from the mean of the empirical background (indicating entropic mixing). Cells were also excluded that had >90% or <10% host-batch edges within the sub-network as these distantly connected cells could be spurious and not represent a large scale topological pattern.

### Illustration of assumptions made by standard benchmark metrics

The human 10k PBMC dataset from 10x genomics (and included within the data zip from the repository) was first analyzed to identify “primary” clusters which served as the “cell-state not annotated” references. Next, this dataset was split into random thirds, with each third getting its own cell state signal added.

#### Primary cluster identification

Using Seurat V3, we calculated the normalized dispersion values, using the top 2000 over-dispersed features which were used for clustering via PyMINEr, in which the cell-cell Spearman correlation matrix was calculated, which was then fed into the cell-cell Euclidean distance calculation. These distances were then used to create the locally weighted kNN followed by Louvain modularity as described in detail later in this methods section.

#### Cell-state spiking using underdispersed template genes

Some have questioned the utility of use of simulation based on our assumed parameterized distributions. To circumvent this concern, we used actual gene expression observed from within this dataset as templates. Given that we used a split of three, we desired three brackets of low, medium, and high expression in which we randomly assigned a bracket (low/mid/high) to each gene from the gene-set specific to each data split such that for gene-1 all of the cells from split-1 received low-template, split-2 received high, split-3 received mid; and gene-2 cells of split-1 received high, split-2 received mid, split-3 received low, etc. Each bracket to split assignment was performed randomly so that total counts were not wildly different between splits as would happen by assigning all of split-1’s spiked-in signal to the low bracket, split-2 to mid, split-3 to high, etc.

The number of genes that were needed as templates (n) from each bracket was equal to the number of target genes (defined in the next section). Summary statistics at the gene level for expression and variability were obtained through Seurat using the HVFInfo function. Non-expressed genes were removed, then genes were ranked by their mean expression and converted into percentiles. The low bracket was defined as beginning at the 15^th^ percentile, mid: 65^th^, high: 92.5^th^. Next, 2-times the number of required genes (n) were selected moving up from these lower bounds. Each bracket’s genes were then annotated as the n genes within the 2n selection that had the lowest dispersion values. Each genes’ bracket and dispersion status are visually displayed in **Supplemental Note-1**; **Supplemental Fig. 1**.

#### Identification of genes to target for spike-ins

We aimed for these spike-in genes to be of biologically cohesive and meaningful signal. To this end, we begin with a list of all genes from several go pathways relevant to MEK/ERK/ JAK-STAT signaling (GO:0000165, GO:0070371, GO:0007259), T-cell Activation (GO:0042110), T-cell differentiation (GO:0045580), and immune system process (GO:0002376). Genes of these pathways were collated via gProfiler, then filtered for only those genes that were not previously annotated as over-dispersed and used for identification of the primary clusters. This prevents any data leakage because the original labels were generated agnostically of the expression of these non-over-dispersed genes from these biologically meaningful pathways. Similarly, the above described template genes did not include any of these target genes to prevent self-sampling, again towards the goal of preventing any data-leakage.

#### Spiking signal into only a single population

In the experiments where we spiked signal into only a single cluster, we identified this “primary cluster” as described above that showed the highest expression of *CD3E*, a pan-T-cell marker. All other attributes of this analysis were identical to those described above, except for the fact that only this cluster was annotated as being split into thirds, and correspondingly, only this cluster received the spiked in signals to each of the three splits.

#### Assessment of status quo benchmarking metrics for the apparent performance of methods with cell-state annotated vs non-annotated labels

The three splits of the dataset were integrated using the same pipeline as described in the above section. We assessed: Adjusted Rand Index (ARI) by batch, ARI by celltype, normalized ARI by batch, normalized ARI by cell type, fscore ARI, average within distance, “average distance to nnkth nearest neighbor within cluster”^48^ (mnnd), “coefficient of variation of dissimilarities to the nearest within-cluster neighbor measuring, uniformity of within-cluster densities, weighted over all clusters”^48^ (cvnnd), maximum cluster diameter, widest gap between clusters, separation index, minimum separation between cluster, average silhouette window (ASW), dindex (“this index measures to what extent the density decreases from the cluster mode to the outskirts”^48^), denscut (“this index measures whether cluster boundaries run through density valleys”^48^), highdgap (“this measures whether there is a large within-cluster gap with high density on both sides”^48^), “correlation between distances and a 0-1-vector where 0 means same cluster, 1 means different clusters”^48^ (Pearson gamma), within cluster sum of the squared distances, entropy, average distance to the centroid of a cluster (pamc), average between cluster distance, minimum of average distance to other cluster, minimum separation between clusters, minimum of the average cluster silhouette width, lowest iLISI, average iLISI, highest cLISI using cluster labels with either the spiked cell state observed or not, average cLISI with spiked cell state observed or not, highest cLISI using “observed state” labels, average cLISI, the kBET average rejection rate, kBET-cell-type rejection rate, kBET-cell type with observed cell-states, mean and max kBET within primary clusters, mean and max kBET within cell-state annotated labels. R packages used for these metrics included: kBET^29^, lisi^10^, and fpc^48^. All quoted descriptions come from the fcp package documentation.

These metrics were compared for when cell state was “observed” vs “not-observed” based on primary clusters, or those split into the thirds as described above. The Spearman and Pearson correlations between metrics were quantified within these different cluster labels being used as noted in **Supplemental Note-1; Supplemental Fig. 2**. Given the large number of metrics quantified, we display only several of interest that showed discordant results when the cell-state was annotated by the experimenter or unknown to the experimenter.

#### Assessment of the impact of PCA reduction in generation of the reference primary clusters

The entire above procedure was repeated a second time, but first using reference labels that were generated by taking the first 30 principal components, then using them for input to the symmetric Euclidean distance matrix calculation prior to locally weighted kNN creation and Louvain modularity. This is in contrast to using the cell-cell Spearman correlation matrix as input for the calculation of the Euclidean distance matrix, thus preventing the dataset from ever being compressed into a low dimensional space.

Similar to comparing cell-state observed vs not-observed, we also quantified the correlations between PCA reduced vs non-reduced when generating reference labels as shown in **Supplemental Note-1**; **Supplemental Fig. 2**.

### Identification of T_m_ complexes through StringDB

Once the complete list of DEGs was collated for a given technical replicate comparison via WCAB analysis, within cluster Spearman correlations were performed within those DEGs, using a bootstrap shuffled null background to account for any non-normal distributions inherent to scRNAseq data, using a false positive rate cutoff of 0.05. These correlations were represented as an undirected graph network using iGraph^49^. This graph was then partitioned into connected components. The genes from each component were then pulled from the StringDB network of protein-protein interactions (PPI) graph networks^25^. These PPI networks were then used to create the final co-regulated (based on correlation) physically interacting graphs (based on the StringDB annotations). These graphs were then subjected to Louvain modularity to identify highly connected sub-networks which were then annotated by performing gProfiler analyses^24^, using the globally expressed genes in these datasets, that were *also* present within the StringDB database as the custom background, therefore preventing any generic enrichment for database annotation or high expression. When merging Tm lists between mouse brain and heart datasets, in some cases a gene appeared in different T_m_ complexes (due to slight differences in the boundaries found between clusters as called by Louvain modularity); to prevent double-counting these genes, it was assigned to the smaller complex. Consensus T_m_ annotations were arrived upon by taking the intersection of the T_m_ annotated genes from the mouse brain and heart technical-replicates.

### Quantification of non-technical DEG pathways

We aimed to be very conservative in identifying putative “biologically important” pathways that were differentially expressed across non-technical replicate comparisons. To this end, we performed within-cluster/across-batch DEG analysis as described above followed by gProfiler pathway analysis using cluster-level custom backgrounds based on what was expressed within this cluster. These significant pathways were then filtered to remove any pathway that appeared at all in the union of both brain and heart technical-replicates. Given the premise that all observations are the union of biological and technical, removing the technically differentially expressed pathways leaves the biologically differentially expressed pathways.

### scRNAseq simulations

#### Simulating cell-types

Using Splatter,^50^ we simulated 1500 cells with 10,000 genes using the parameters: “de.downProb”=.25, “de.prob”=.75, “de.facScale”=.75, “bcv.common”=.75. All other parameters were left at their default levels. Each cell was uniformly randomly assigned to one of 6 clusters, and to either batch 1 or batch 2, depending on the details of the simulation as noted in **Supplementary Fig. 9**.

#### Simulating cell-states

To simulate a cell-state module, a single ‘cluster’ was simulated with the same dimensions as the original dataset. All cells were then paired with a cell-state transcriptome. 25% of genes were then considered to be members of the cell-state. A gaussian weighting vector was generated using rnorm * 1/50 to shrink variance, and shifted to center around 0.25. Each gene belonging to the cell-state was assigned to a weight from this distribution. A linear mixture of the cell transcriptome and states is then created to form the final cell, using weights for the mixture similar to above, mixing cell-type and state with random weights either centered at 33.3% state or a half-Gaussian whose lower bound was zero for cells that were being injected with the cell-state or not respectively.

### Statistical analyses

Statistics comparing percent-same, PMD, relative mutual information, purity, and reverse purity across methods were performed using an ANOVA via the aov function in R on rank transformed versions of the metrics, given that variance was different across groups:

> aov(rank(<metric>) ∼ method)

For analysis of simulations, the number of groups shared across batches were also modeled as a covariate:

> aov(rank(<metric>) ∼ method + num_groups_shared)

TukeyHSD was used for post-hoc comparisons. All statistics were analyzed as two-sided comparisons.

### Clustering

#### Feature selection prior to clustering

Because the goal of this work was to examine the real-world realistic implementations of a standard bench-biologist following tutorials as advertised, feature selection algorithms were implemented as described in each methods’ tutorials. When not available, and the corrected version of the transcriptome as a ‘corrected expression matrix,’ automated highly variable gene selection, and clustering was applied equivalently to all methods that yield corrected transcriptomes, as previously reported by PyMINEr^51^. The mean variance relationship was fit by a lowess locally weighted regression curve. Residuals were calculated, and those genes whose residual was ≥2 standard deviations above the loess fit curve were selected. For methods yielding latent dimensions, these dimensions were used for clustering rather than applying additional feature selection as this is typically performed *before* latent dimension mapping.

#### Clustering

Clustering is performed similar to standard practice with Louvain modularity, with one modification; instead of approximating similarity space through compression of data into a latent space, we measure actual similarity/distance space, staying in high dimensions without latent dimension compression. The locally weighted kNN used for Louvain modularity was constructed as follows:

- Calculate the symmetric pairwise Spearman correlation matrix of each cell against all other cells
- Calculate the squared Euclidean distance of all cells to all other cells using the Spearman correlation matrix as input.
- For each cell, mask the 95% most dissimilar cells (or all but the 200 closest cells, whichever gives the lower number of connections), and the diagonal, self-distance=0, thus creating a local distance matrix.
- For all remaining cells that were not masked, the inverse Squared Euclidean distances were linearly normalized between 0 and 1 within each cell. This creates a local weighted distance matrix, which is then converted to a weighted graph, treating the local distance matrix as a weighted adjacency matrix, which is then subjected to Louvain modularity based clustering.

This process is fully automated in PyMINEr,^51^ hyperparameter free, and implemented equally to all batch-correction methods, using the -louvain_clust argument. Louvain modularity was used via the python-louvain package.

### TOWCAB assessment of published atlas based on published coordinates

These analyses were performed with the script called bin/from_coords.R within the main repository. Coordinates were converted to a kNN with k=ceil(sqrt(n)), where n is the number of cells in the dataset using the FastKNN package’s k.nearest.neighbors function based on the cell-cell Euclidean distance matrix calculated by the pdist function from the rdist package. This kNN and the corresponding expression matrix were fed into the run_towcab_analysis function within the towcab package. Figures were made using the UCSC Cell Browser (https://cells.ucsc.edu) browser that hosts the data^52^.

### Clustering performance metrics

The “percent-same” metric that was implemented in some biologic benchmarks is calculated by taking the pairs of cells from the original dataset, and its downsampled counterpart, and quantifying the number of pairs that were placed into the same cluster (numerator), relative to the total number of pairs (denominator). This metric only quantifies cells that are present in both datasets; if a parent or child cell was removed by a pipeline for quality control purposes, they were not counted given that it would not have a matching cell-pair to assess if the cluster was the same.

Mutual information was calculated using the mi.empirical function from the entropy R package. Relative mutual information is the ratio of the observed mutual information to the theoretic maximum mutual information, if the results were 100% accurate relative to the ground-truth clusters.

Purity was calculated using the purity function from the NMF package in R.^53^ The purity function takes in two arguments, first: the cluster results, and second: the ground-truth clusters. Purity is a metric that takes the method’s clusters, identifies its plurality ground-truth cluster, then returns the percent of all cells that fall into a method’s cluster whose plurality is the same ground-truth cluster. This particularly penalizes for merging ground-truth clusters together.

Reversing the arguments of the purity function does the opposite however, we therefore use reverse purity to quantify splitting ground-truth clusters (reverse purity).

The “true percent max difference” is reported in **ED Table 1**; this metric calculates the PMD based on the ground-truth known clusters in the simulated batches. This metric is used in combination with the percent maximum difference calculated using the observed clustering results for each method. The log ratio percent maximum difference is ln(PMD_observed_/PMD_ground-truth_). Difference in percent maximum difference is PMD_observed_-PMD_ground-truth_. Absolute difference in percent maximum difference is |PMD_observed_-PMD_ground-truth_|.

The “state merged with base purity across batch” is the typical purity calculation, but first filtered to include only cells from clusters for which the base cluster appears both with and without the state added, thus penalizing specifically for merging clusters that belong to the same base cluster with and without a state. This quantifies an algorithm’s propensity for erasing batch confounded ‘states’ or expression programs.

The “state merged with different base purity across batch” quantification is the purity function applied specifically to the filtered subset of ground-truth clusters with the cell-state added that did not belong to the same base “cell-type.”

The “split by batch reverse purity” metric is the reverse purity metric but selectively applied to the subset of cells that belonged to ground-truth clusters that appeared in both batches, whereas cell clusters that appeared only in a single batch were removed.

### Reanalysis of brain datasets for CD19+/mural-marker+ populations

Because no code or datasets were released with the original manuscript^42^, we could not re-perform the exact procedures used there. Where did however perform our own independent analysis of the same datasets which were release elsewhere in their original papers. All analysis and code is contained in the “bin/cd19_mural_cell_analysis.R” script in the primary benchmark repository (https://bitbucket.org/scottyler892/sc_norm_bench_v2). Reprocessed data is included in the data distribution Rds or is downloaded on the fly within the script.

#### Supervised annotation of B-cell-related and mural-cell related droplets and candidate CD19+/mural-marker+ copositive candidate droplets

To provide a strict conservative definition of droplets that were the most “pure” in identity, we defined conservative “pure” B- and mural-droplets as those containing ≥5 UMI for any of the indicated markers, with 0 sum expression of across-lineage markers: B-cell markers were: (*Vpreb1, Vpreb3, Ighm, Ighg3, Ighe, Ighg1, Ighg3, Ighd, Igll1, Sdc1, Tnfrsf8, Ms4a1, Cd79a, Cd79b*); mural-cell markers were: (*Cd248, Rgs5, Foxf2, Pdgfrβ, Cspg4).* The ‘candidate’ populations were defined as droplets with any observed expression of CD19 and simultaneous detection of a mural cell marker. The “clearly mixed” populations were defined as cells with ≥5 UMI for at least one mural marker, but also showed ≥10 UMI total expression of B-cell related genes (listed above), lymphocyte/leukocyte related genes: (*Ptprc, Cd38, Cd34, Cd3e, Cd3d, Cd3g, Cd3a, Il2rA, Cd28, Ctla4, Cd4, Cd8a, Cd8b; Cd14, Cd80, Cd84, Cd86, Tnfrsf4*), or MHC-II genes [mouse:(*H2-K1, H2-Ke6, H2-Oa, H2-DMa, H2-DMb2, H2-DMb1, H2-Ob, H2-Ab1, H2-Aa, H2-Eb1, H2-Eb2, H2-D1, H2-Q1, H2-Q2, H2-Q4, H2-Q6, H2-Q7, H2-Q10, H2-T24, H2-T23, H2-T22, H2-T3, H2-M10.2, H2-M10.1, H2-M10.3, H2-M10.4, H2-M11, H2-M9, H2-M1, H2-M10.5, H2-M10.6, H2-M5, H2-M3, H2-M2*), human:(*HLA-DMA, HLA-DMB, HLA-DOA, HLA-DOB, HLA-DPA1, HLA-DPB1, HLA-DPB2, HLA-DQA1, HLA-DQA2, HLA-DQB1, HLA-DQB2, HLA-DRA, HLA-DRB1, HLA-DRB5, HLA-DRB6*)].

Marker genes of other lineages were used only for heatmap displays rather than analysis or identification of populations. These included the following markers: [astrocyte:(*Gfap*); erythroid:(*Hbb, Hbg1, Hba1*); oligodendrocyte precursor cells(OPCs):(*Sox10*); microglia:(*Csf1r*); endothelial:(*Cdh5, Pecam1, Vcam1*); excitatory neurons:(*Neurod2*, *Neurod1*); inhibitory neurons:(*Lhx6, Gad1, Dlx5, Dlx2*).

#### Number of observed genes and counts

The number of observed genes and total counts per droplet have been previously described as a characteristic of doublets/multiplets^54^, and was therefore analyzed and found to be higher in candidate CD19+/mural-marker+ copositive droplets as noted in the manuscript.

#### Transcriptome correlation and MSE with 50:50 linear model of “pure” B-cell and “pure” mural-cell transcriptomes

After “pure” B- and mural-related droplets were identified, the average B- and mural-related transcriptomes were calculated. Then the average log2(counts+1) of these two transcriptomes was taken as the putative 50:50 mixture. Note that to account for differing abundance of these populations, we took random samples of droplets to equal depth (minimum count of either population of cells) over 20 iterations to generate a distribution of fits from random subsets of each reference controlling for an equal number of cells in each reference. This accounts for any excess “smoothing” and lower error that would result simply from sampling a greater number of cells. This transcriptome vector was used as a reference for both Spearman correlations and mean squared error (MSE) calculations with a ‘test’ population. This ‘test’ population was either the candidate CD19+/mural-marker copositive droplets, or a random subset of droplets from the “pure” mural population. Each ‘test’ population was quantified for its fit with sampling from a single cell through to all candidate cell populations, and a matching population of control droplets from the “pure” mural population (held out from references) (x-axis on **Supplemental Note-6; Supplemental Fig. 11c**); assaying this range of sample depth also controlling for the any “smoothing” differences seen by simply sampling more cells. This comparison allowed us to directly compare whether the CD19+ candidates are globally more B-like than the pure mural population.

Note that in the scenarios where we fit the “pure” mural subsets to the 50:50 reference, the test subset was excluded from the “pure” mural population used to generate the reference average, thus preventing data leakage. All marker genes used for identification of these populations were excluded from these analyses to both 1) ensure no data leakage, and 2) that we were quantifying the broad transcriptome patterns independently of the expression of CD19, further indicating that these cells were not simply mural cells that expressed CD19.

#### Fitting a linear mixed model of “pure” B- and mural-references

A linear model of the average log2(counts+1) candidate and reference pure-mural and pure-b transcriptomes was calculated as described above, but instead of assuming a 50:50 mixture, we quantified the relative proportion of these references that explain the candidate transcriptome by fitting a linear mixed model: candidates = b_1_*pure_mural_reference + b_2_*pure_b_reference, with an intercept of 0. The b_1_ and b_2_ coefficients were calculated by the FLXglm function from the flexmix package^55^ with the offset argument set to zero, fixing the intercept, thus preventing the attribute of overfitting that result from different sequence depth. Every gene not included in the marker-sets used to identify these populations was quantified independently, yielding a measure at the per-gene level quantifying the contribution from each lineage. Relative contributions of each “pure” population was calculated as b_1_/sum(b_1_+b_2_) and b_2_/sum(b_1_+b_2_), removing the net impact of differential ‘count-depth’ for specific cell subsets. For example, a random Poisson sub-sampling of a single transcriptome vector by 50% may symmetrically decrease the beta-coefficients by half, yet their relative contribution would remain the same ± random Poisson sampling error.

### Protein staining on human brain and tonsil

#### Samples

Samples were obtained from BioMax USA (Human tonsil: HuFPT161, Brain: HuFPT017, HuFPT018, HuFPT019, HuFPT020).

#### Sample preparation

Slides were baked for 1hr at 50C, then deparaffinized as follows: Xylenes 10 min, 2x 100% EtOH, 10 min, 2x; 95% EtOH, 5 min; 70% EtOH, 5 min; 50% EtOH, 5 min; rinse in diH2O; PBS, 10 min.

#### Antigen retrieval

Antigen retrieval was performed using citrate buffer in an instapot pressure-cooker. Pressure cooker was filled with fresh diH2O, with a vegetable steamer insert placed inside to hold plastic staining jar containing slides submerged in 220mL of 1x citrate buffer made from 10x concentrate (Invitrogen #00-4955-58). Slides were subjected to full temperature/pressure cooking conditions for 3 minutes excluding ramp time. Pressure was released and samples were allowed to cool to on a bench-top for 30 minutes prior to transfer into PBS.

#### Pre-bleaching

To decrease autofluorescence, slides were pre-bleached as previously described^56^. Place slides tissue side up in glass dish filled with 1L pre-bleach buffer (PBS+0.05% sodium azide. Glass dish was placed on reflective aluminum surface inside of a 4C walk-in fridge. A cardboard box lined with polyvinyl shrink-wrap (to protect cardboard from evaporating water), and further lined with aluminum foil to reflect light back onto sample was created and placed overtop of dish containing samples. Holes were cut out of cardboard container to allow for insertion of autofluorescence photo-bleaching lamps. Samples were then exposed to high intensity full spectrum LED lights (3x Relassy: Sunlike Full Spectrum Grow Light 45 Watt light incandescent equivalents and 1x SANSI full spectrum 400 Watt incandescent equivalents) for 48 hours. Note that some residual autofluorescence existed, highlighting the importance of the no primary negative controls that we employed. Sodium azide containing PBS was then disposed of in hazardous waste. Samples were then washed 3x in PBS.

#### Protein staining

Samples were blocked for 2hrs at room temperature in blocking buffer (PBS with 1mM CaCl_2_, 1mM MgCl_2_, 20% Donkey serum (Jackson Immuno Research: 017-000-121), 0.3% Triton X-100). Samples were stained in staining buffer (PBS with 1mM CaCl_2_, 1mM MgCl_2_, 1% Donkey serum, 0.3% Triton X-100). Antibody dilutions were as follows: goat anti-PDGFRβ 1:100 (Novus Biologicals: AF385), rabbit anti-CD79a 1:100 (Fisher Scientific PIMA516344; clone SP18), mouse anti-CD19 1:50 (GeneTex: GTX42325, clone LE-CD19).

Note that CD19 primary (1:50) and secondary (1:50) were used at higher concentrations than other channels to address any concerns over lack of sensitivity. Our positive control in human tonsil showed abundant signal as expected indicating the efficacy of our staining procedure.

Slides were washed 3x with PBS (5 min each), and stained in staining buffer with secondary antibodies, each from Jackson Immuno Research: 1:50 Alexa Fluor 488 AffiniPure Donkey Anti-Mouse IgG (H+L) (715-545-151); 1:100 Rhodamine Red-X (RRX) AffiniPure Donkey Anti-Rabbit IgG (H+L) (711-295-152); 1:100 Alexa Fluor 647 AffiniPure Donkey Anti-Goat IgG (H+L) (705-605-147). Slides were then washed 3x with PBS (5 min each) and mounted in Prolong Diamond Antifade with DAPI (Fisher Scientific: P36962) and sealed with nail polish. Images were all collected using the same laser settings on a Leica SP8 confocal microscope.

#### Image processing

Images were processed in ImageJ, using the median fluorescence value for each channel in the secondaries only negative control, as the minimum inclusion value, to provide an objective negative control for thresholding. This was performed for all donors and all channels. Donor HuFPT018, showed unquenchably high global background in all cells however, (equally between the test and no CD19 primary antibody conditions), and was therefore excluded.

### Software versions

All R analyses were performed in R version 3.6.0 (2019-04-26) -- “Planting of a Tree”. R packages require for analysis were: splatter: 1.8.0, scater: 1.12.2, Seurat: 3.1.2, DESeq2: 1.24.0, sva: 3.33.2, irlba: 2.3.3, liger: 0.4.2, harmony: 1.0, BiocParallel: 1.18.1, ggplot2: 3.3.0, reshape2: 1.4.3, pals: 1.6, tidyverse: 1.3.0, stringr: 1.4.0, scales: 1.1.0, entropy: 1.2.1, NMF: 0.22.0. Python was version 3.7.1.

### Data Availability

All dataset inputs used in this manuscript are distributed in the data folder/zip of the benchmark repository, or are public data that are downloaded by the scripts on the fly during analysis: https://bitbucket.org/scottyler892/sc_norm_bench_v2

### Code Availability

All code for this benchmark is freely available at the following repository: https://bitbucket.org/scottyler892/sc_norm_bench_v2

The UMI downsampling package we created is available by pip installation: python3 -m pip install bio-pyminer-norm

The R version of the downsampling package is located here: https://github.com/scottyler89/downsample and can be installed via the devtools package: devtools::install_github(’scottyler89/downsample’)

The UMI downsampling package repository and tutorials are located here: https://bitbucket.org/scottyler892/pyminer_norm/

The R implementation of PMD and associated functions/statistical tests are available here: https://github.com/scottyler89/PercentMaxDiff and can be installed via the devtools package: devtools::install_github(’scottyler89/PercentMaxDiff’)

The R implementation of towcab and pathway analyses is available here: https://github.com/scottyler89/towcab and can be installed via the devtools package: devtools::install_github(’scottyler89/towcab’)

---

**Extended Data Figure 1:**
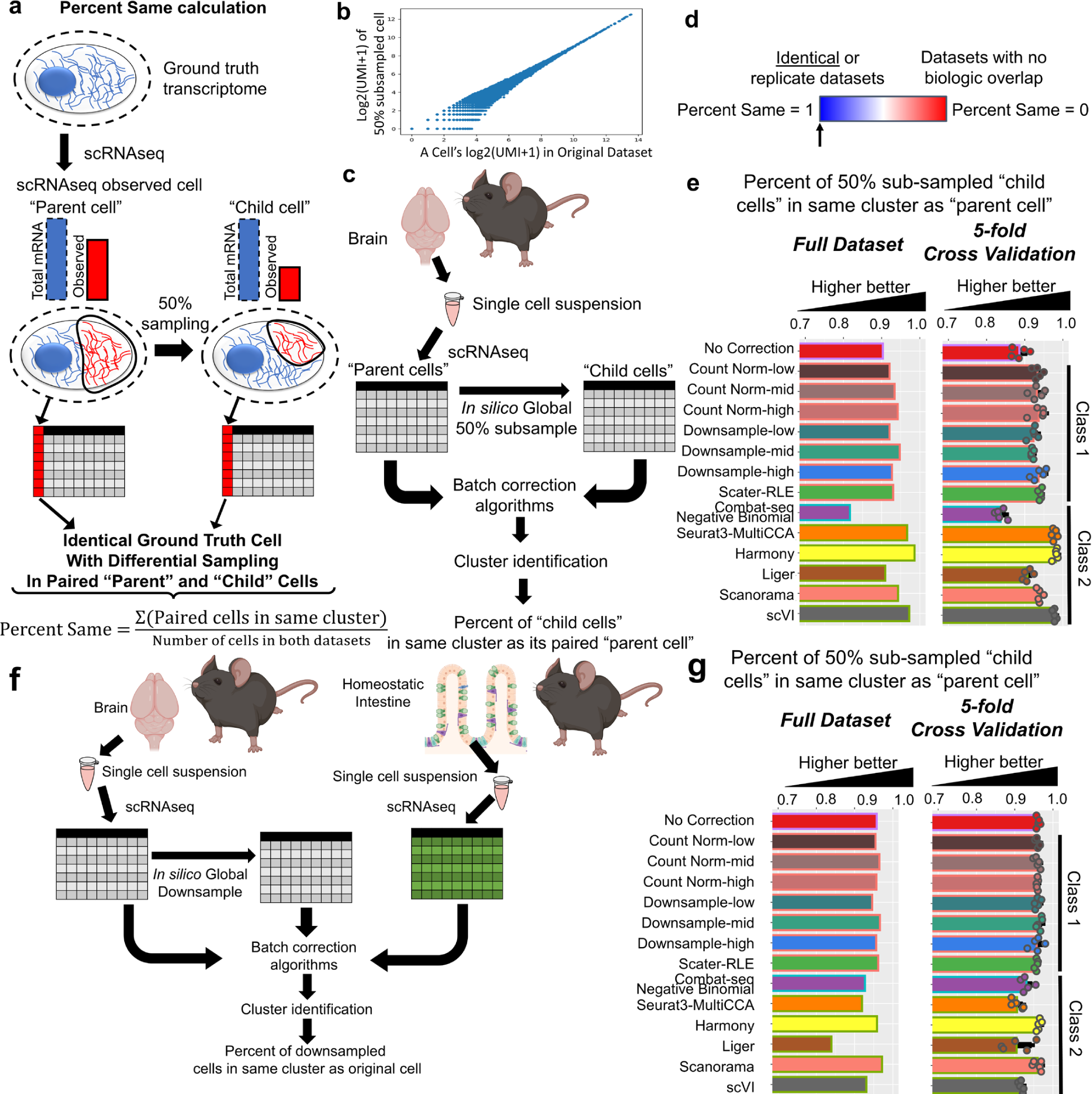
Performance in integrating identical datasets with differences only in UMI depth. **a**, Example schematic of the same cell with different transcript capture efficiency leading to different observed total UMI. **b**, Scatter plot showing a brain scRNAseq dataset that has been globally downsampled at random. Each point is a transcript’s log2(UMI+1) measure in the parental cell (X-axis) and downsampled cell (Y-axis). **c**, Pipeline overview for benchmark schematic. **d**, The percent-same should equal 1, because the datasets are from the same source droplets. **e**, A bar chart displaying the percentage of downsampled cells that were classified as belonging to the same cluster as its parental cell. A higher score indicates better performance. **f**, The overall experimental and pipeline schematic of this figure of merit for results in **g**. **g**, Bar charts of the percent-same metric in the full datasets or cross-validation splits (means +/− SD).

**Extended Data Figure 2:**
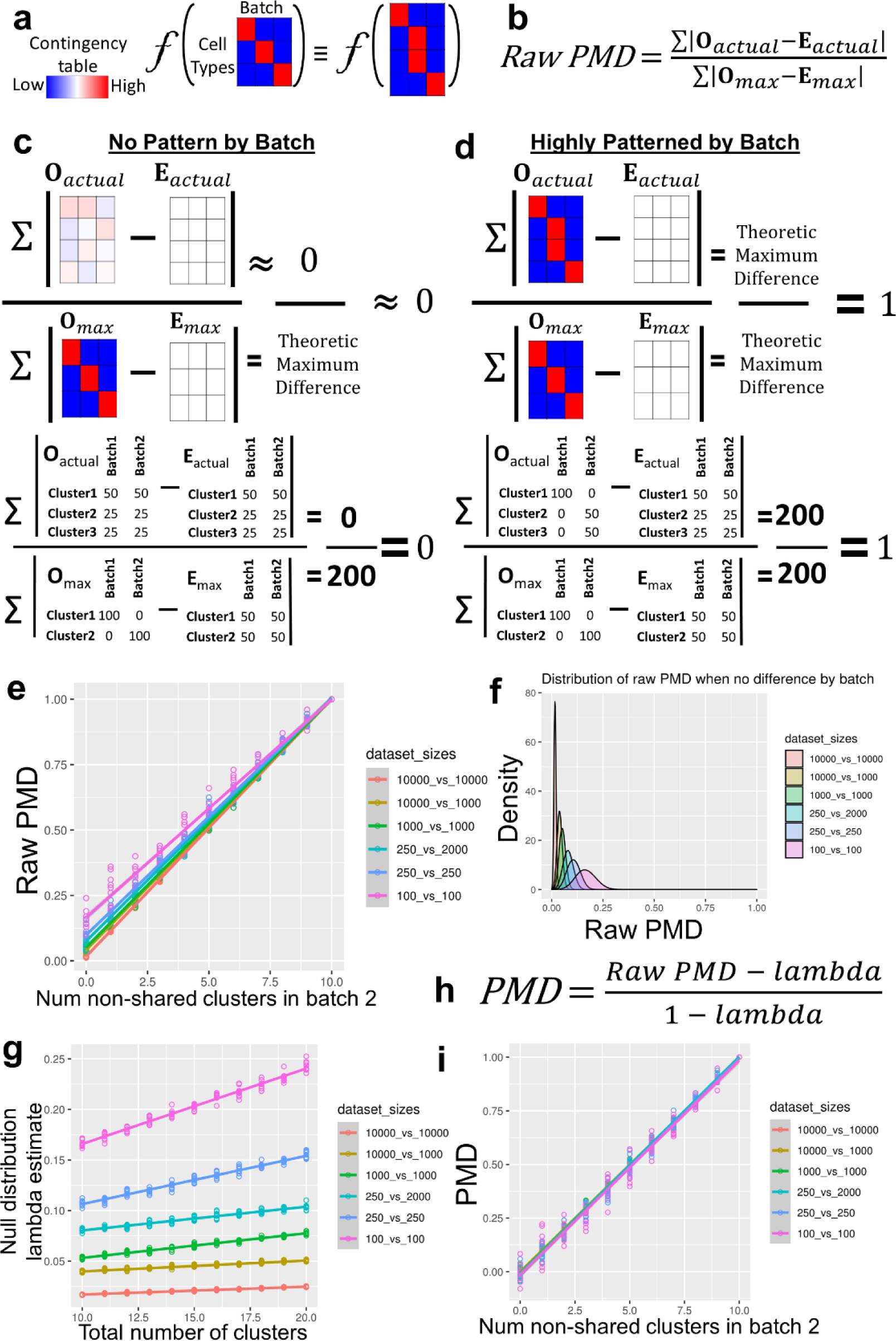
Percent Maximum Difference calculation. **a**, A schematic showing that an ideal metric quantifying differences in batches should be invariant to the number of clusters found within each batch if relative overlap across batches is unchanged. **b**, The equation for the Raw Percent Maximum Difference (Raw PMD) (before empiric power correction). **c**, A visual example of the Raw PMD calculation where there is very little asymmetry across batches. **d**, In contrast to the scenario in **c**, when the actual clustering results are highly asymmetric by batch, the numerator and denominator are equivalent and this ratio equals one. **e**, Raw PMD (y-axis) plotted against the number of clusters that are specific for a given batch (x-axis: higher is more batch asymmetry); simulations with different batch sizes are noted by color. **f**, Histograms of the intercepts of the lines in **e** revealed Poisson distributions, whose lambda parameter (peak of the distribution) differed based on the number of cells simulated (colors). **g**, Lambda parameter estimates (y-axis) of bootstrap shuffled null versions of the simulations shown in **e**, varied with the number of cells (color) and degree of overlap (x-axis), indicating that Raw PMD can be corrected by measuring the lambda parameter. **h**, The final PMD calculation, adjusted for power with the lambda parameter of the null distribution corrects for the intercept (numerator) and slope (denominator). **i**, After adjustment, PMD (y-axis), is linear with the degree of overlap between batches (as noted by number of batch specific clusters, x-axis), and equivalent regardless of the size of these batches (color).

**Extended Data Figure 3:**
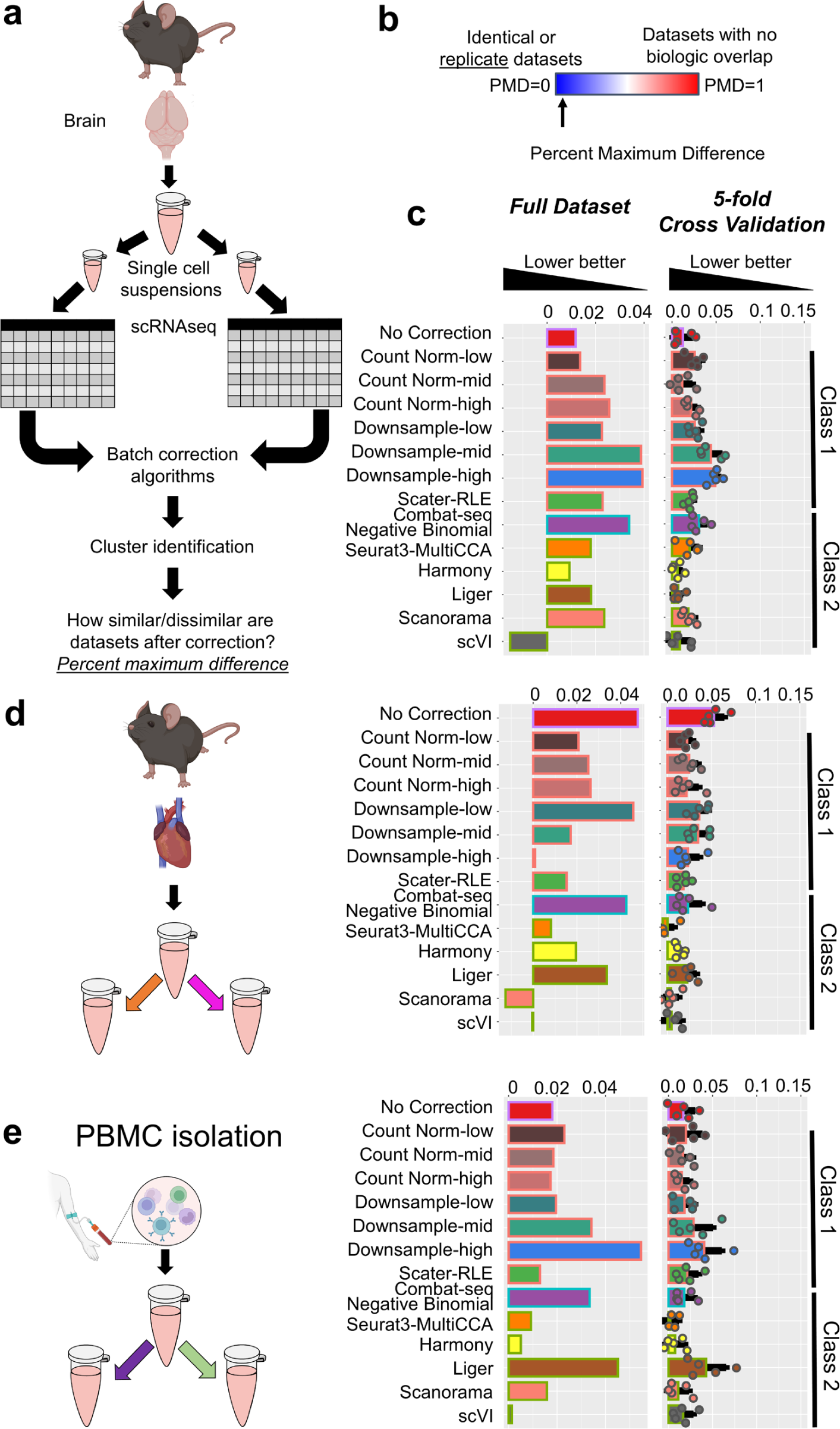
Batch-correction algorithm performance on merging technical replicates. **a**, A schematic representation of the experimental and analytic pipeline for this data integration example. **b**, Technical replicates share identical cluster composition, meaning that PMD should equal 0. **c**, A bar chart showing the percent maximum difference between full datasets and five-fold cross-validation subsets. **d-e**, Technical-replicates of (**d**) mouse heart and (**e**) human PBMCs were also assessed. Means+/−SD are shown for cross-validation splits.

**Extended Data Fig. 4:**
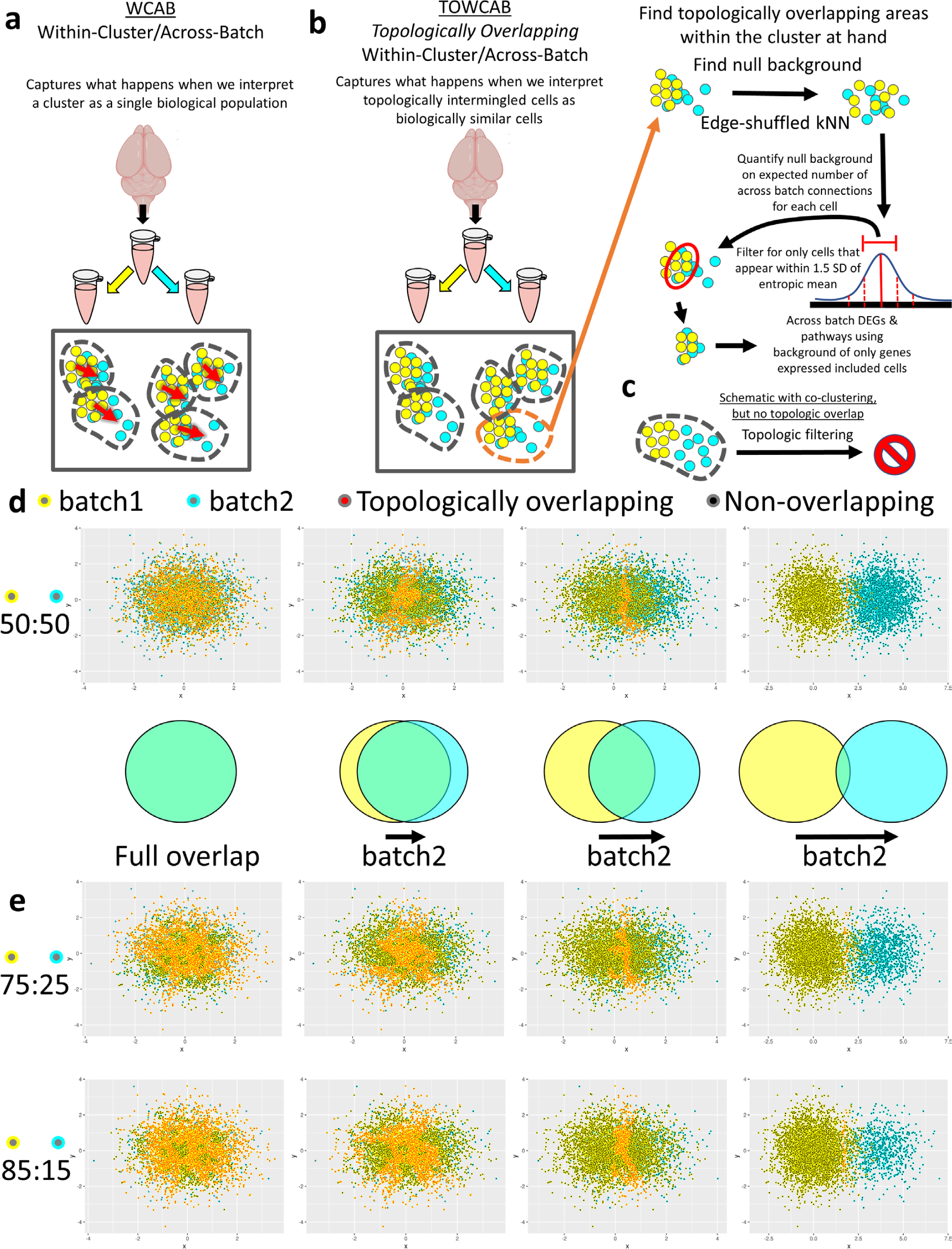
WCAB and TOWCAB analysis schematic. **a**, Within-Cluster/Across-Batch (WCAB) analyses take in the original count matrix, batch information, and clustering results. Within clusters, differential expression analyses are performed on a downsampled version of the datasets, providing the most conservative assessment of relative differential expression/detection with total counts per cell held constant. **b**, Topologically overlapping WCAB (TOWCAB) refines this approach, filtering for only cells that strongly match a null distribution of what entropic mixing of this topology would look like, therefore keeping only cells that are thoroughly mixed across batches within this subset of the topology. **c**, While WCAB analyses will include results in which two batches are identified as a single cluster, when in fact they were simply “zippered” along an edge, TOWCAB analyses will not find any topologically overlapped areas within the cluster because they are only adjacent rather than overlapping. **d**, A simulated example dataset shows this filtering process as implemented in our R towcab package. Displayed points are considered as a single cluster, with varying degrees of overlap by batch (colorized by border), while the center color indicates whether the point was considered thoroughly mixed enough to be considered “topologically overlapping” by batch. **e**, The use of bootstrap shuffled null distributions enables efficacy in discovering topologically overlapped regions even with class imbalance.

**Extended Data Fig. 5:**
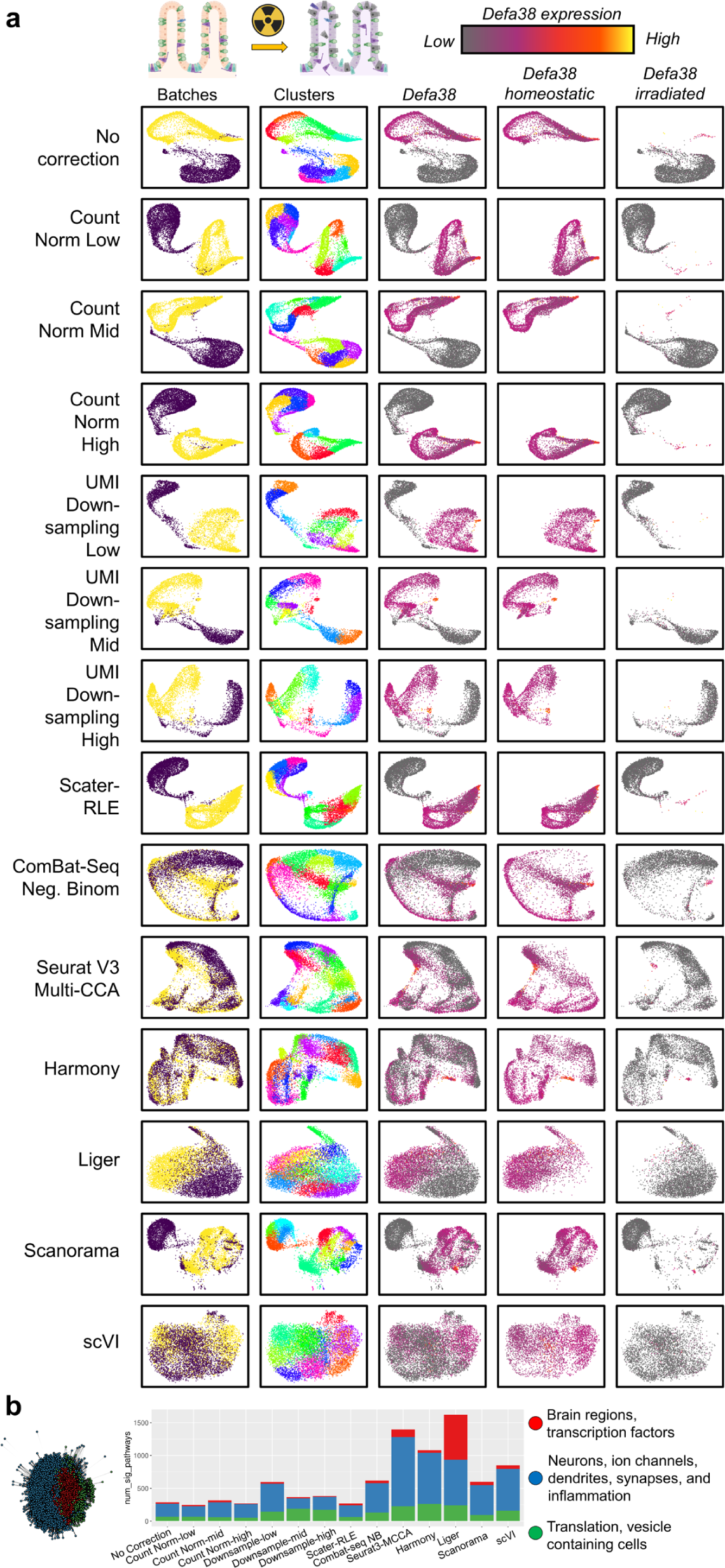
Erasure of cell-state or tissue relevant transcriptional signal by Class-2 methods in mouse intestine and human brain. **a,** Spring embedding plots of the kNNs used for clustering show that defensin alpha 38 expression is lower in irradiated intestine. Despite this, many Class-2 algorithms merge these topologies, fitting with the TOWCAB analyses shown in Fig. 4. **b**, TOWCAB analysis of brain regions; the graph network shows nodes representing pathways, connected by edges based on relative overlap in gene-lists, colorized by Louvain modularity to identify the pathway clusters. A bar chart showing the number of significant pathways whose signal was significantly differentially expressed within topologically overlapping regions of the cell-cell graph. All Class-2 algorithms showed a greater number of differentially detected/expressed pathways compared to Class-1 algorithms. Cluster-1 and −2 show clear biological importance to the brain.

**Extended Data Figure 6:**
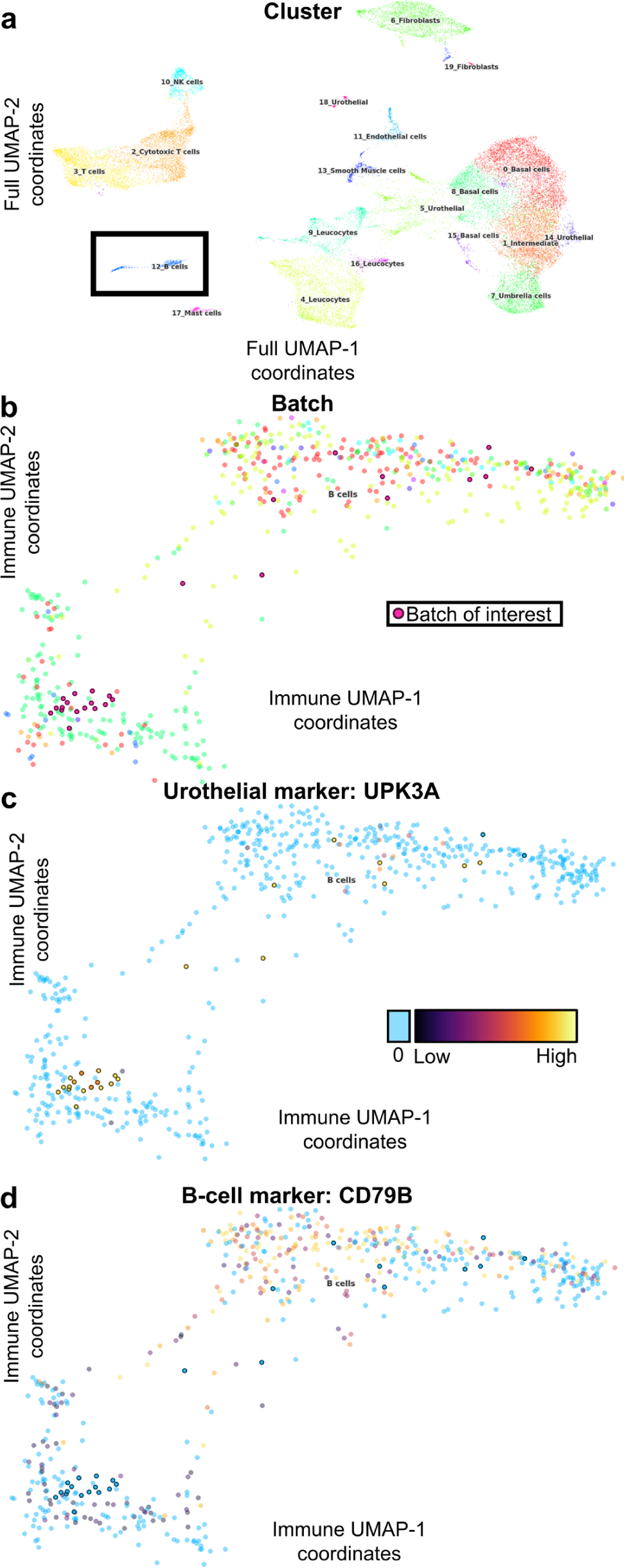
Application of TOWCAB reveals erasure of cell-type signal in an atlas embedding. **a**, Published UMAP coordinates of adult human ureter colorized by cluster; the cluster labeled as B-cells is boxed. **b**, Cells annotated as B-cells along the published Immune compartment-specific UMAP coordinates colorized by batch. Note the red batch with darker circles indicates the batch of interest. **c,d**, TOWCAB analysis identified differential expression of *UPK3A* and *CD79B* across batches. **c**, Cells from the batch of interest are expressing the urothelial marker *UPK3A*. **d**, Cells from this batch are not expressing the B-cell marker *CD79B*.

**Box 1:** A brief overview of the figures of merit based on real biological datasets, and synthetic datasets, indicating whether they are testing how approaches perform on similar or dissimilar datasets and whether or not cell state information is included in the figure of merit or synthetic benchmark.

|  | **Comparison Type** | **Comparison Description** | **Experimental Setup** | **Figure of Merit** |
| --- | --- | --- | --- | --- |
| **a** | Experimentally identical | Cells of a single tissue compared to subsamples of the transcriptomes of the same cells | Subsample from source “parent” cells to create “child” cells with all cells of the same tissue type | Cluster across all parent and child cells and compute the percentage of times child cells cluster with parent cells (**ED. Fig. 2a-e**) |
| **b** | Experimentally identical in context of highly dissimilar | As above (**Box 1a**), but in the context of unrelated cell types | Subsample from source “parent” cells to create child cells where these cells are of the same type, but mixed with unrelated cell types | Same as above **Box 1a** (**ED Fig. 1f-g**) |
| **c** | Experimentally differing only by Poisson sampling and technical | Cells from the same single cell suspension | scRNAseq data generated on multiple isolates of the same sample | Percent maximum difference where methods that minimize the PMD are preferred (**ED Fig. 3**) |
| **d** | Experimentally highly dissimilar | Cells of one type compared to cells of an unrelated type | Independent runs of scRNAseq data generated on disparate tissues expected to have little or no overlap in cellular composition | Percent maximum difference where methods that maximize the PMD are preferred (**Fig. 3**) |
| **e** | Experimentally batch confounded cell-state and cell-type abundance. | Cells of a given batch are of similar cell-type, but in different cell-states, and cell-type abundance | scRNAseq data generated on cells derived from normal intestine and cell derived from irradiated intestine, differences in endometrium throughout menstruation. | Percent maximum difference; TOWCAB analyses (**Fig. 4, 4d-e; ED. Fig. 5a**) |
| **f** | Batches assumed to be similar, but may have had biologically meaningful differences | Cells came from the same tissue, and may be assumed to be highly similar, but in reality may harbor biologically meaningful differences. | Within-dataset templated cell-states added, within brain region comparisons (SN vs cortex), and adult ureter atlas, and human brain mural cells. | TOWCAB analyses and protein staining (**Supplemental Fig. 2; Fig. 5,6e-h; ED Fig. 5b,6**). |
| **g** | Mixtures of similar and dissimilar cells | Cells of the same and different types compared for similar and dissimilar groupings. | Simulated batch-confounded and non-confounded cell-type/-states, the multi-tissue *tabula sapiens* atlas. | Various including batch similarity compared to ground-truth, overall clustering accuracy, extent of misclustering by batch (**Fig. 6a-c; Supplemental Fig.** |
|  |  |  |  | **9,10).** Visual inspection<br>of published<br>coordinates<br>**(Supplemental Fig. 1)** |

**Extended Data Table 1: Relative co-mentions of Class-2 algorithms with the term “batch-correction” within the population of manuscripts that contain the phrase “single cell RNAseq”. a**, Tabulated results of method/batch correction co-mentions and chi-square statistics. **b**, Raw data and links that were used for tabulation of co-mentions.

**Extended Data Table 2: Standard benchmarking metrics often rely on assumptions of perfect knowledge by the experimenter. a,** Raw values for each metric assessed when the added state was “observed” or “unannotated” by the experimenter. **b**, Raw values for each metric when reference labels were generated using Euclidean distance based on Spearman correlations vs Euclidean distance based on the first 30 PCs. **c**, The correlations of each metric when the state was either observed and annotated vs not observed or annotated by the experimenter. **d**, The correlations between metric values when reference labels were generated from Euclidean distance of Spearman correlations vs Euclidean distance of the first 30 PCs. **e**, Summary statistics demonstrating that many metrics yield significantly different results dependent on whether the experimenter had perfect knowledge and annotation of all cell-states compared to a situation where cell states remained unannotated in the provided reference labels. **f**, Summary statistics showing that some popular metrics yield significantly different results, rewarding for similarity to the processing methods used to generate reference labels. **g**, Summary of all results comparing correlation consistency or significant differences in means when states are observed vs non-observed.

**Extended Data Table 3: Study of Technical Effects on Chromium V3 with technical-replicates in different tissues and species**. **a**, A collated table displaying the technical class each gene belongs to, for each individual dataset. For each gene, the percentage of clusters for which it was differentially detected within a batch-correction run was quantified. **b-d**, Detailed lists of all genes and batch-correction methods and the percentage of clusters for which that gene was significantly different in the WCAB analysis is reported for (**b**) mouse brain, (**c**) mouse heart, and (**d**) human PBMCs. **e**, A collated table of the intersect of technical effects between moues brain and mouse heart can provide a draft of genes known to be impacted by technical effects. **f**, The union of pathways that were significantly different by WCAB analyses in mouse brain and heart can act as a conservative “background” of pathways that we observed as differing between technical-replicates in either of these tissues.

**Extended Data Table 4:** All results for simulation scenarios, and metrics used for quantifying batch-correction algorithm performance are provided.

**Extended Data Table 5: Pathway results of genes correlated with markers of interest in Zong 2018 dataset. a,** Pathways of genes significantly correlated with CD19 above a bootstrap shuffled null background. **b**, Pathways of genes significantly correlated with CD79a/b.

## Supplemental Notes

### Supplemental Note-1: Status quo methods of benchmarking batch correction

#### Assumptions made by status quo metrics

Most, if not all standard metrics used in benchmarking scRNAseq batch correction methods rely on one of two assumptions: either 1) they rely on the assumption that batches differ only in technical effects, and that there are no batch confounded differences in biological variation (ex: kBET and iLISI); or 2) manually provided cluster labels accurately capture all meaningful sources of biological variation, and clusters with the same label reflect biologically homogenous populations [within cluster average silhouette width (ASW), adjusted rand index (ARI), etc]. Broadly speaking, the later class tests for consistency within the manually provided labels. However, in scRNAseq benchmarks, these manually provided labels come from published annotations, which are arrived at after the application of the original manuscripts preferred analytic pipelines, doing their best to label what they think are pertinent and expected clusters.

The first class of assumptions will hold as correct when there are no biological differences between batches. However, technical replicates are the only scenario under which such an assumption can be taken for granted. Even, for example, within isogenic mouse lines, subtle differences in environment, somatic variants, and epigenetic changes can all dramatically alter molecular biology pathways. One illustration of this is a recent discovery that the sex of the experimenter alters the metabolic pathway of ketamine in isogenic mice. This was ultimately traced back to the scent of human males compared the scent of human females^57^. Additionally, there are many other sources of within-subject environmental variation impacting cell states, such as time since feeding, circadian rhythm patterns, and so on – none of which can be completely controlled. True technical replicates constructed by taking two separate pipette draws from a single tube provides the only near guarantee that samples only differ in downstream technical effects and random Poisson sampling of the original biological sample. Environmental factors can confound our interpretation when benchmarking batch correction methods even when using isogenic mice; with any biological replicate, biological variation within group as well as technical variation are included.

There are also several dangers in relying on previously published cluster labels. The first sub-assumption within these metrics is that the labels capture all meaningful sources of variation. For example, it has been reported that “27% of the β cell transcriptome exhibited circadian oscillation^58^.” This means that if a reference label is simply “β cell,” this biologically meaningful source of variation will not be captured within this label. Furthermore, there may be instances when the application of an experimenter’s prior expectations of the dataset can negatively impact their ability to produce accurate labels. For example, we have found in the *tabula sapiens* multi-tissue atlas, there exists a population of droplets that came from a muscle sample, that was labeled as satellite stem cells^39^. However, these droplets were negative for the markers of this cell type, but were instead positive for B-cell marker genes, yet were inserted into a topologic region annotated as fibroblast/stromal cells following Class-2+LDM integration (scVI) (**Supplemental Fig. 1**)^39^. This highlights the dangers of 1) relying on the use of prior expectations for processing and droplet annotation (since the example just given, the B-cells were labeled as muscle satellite cells and the sample origin was muscle), and 2) this Class-2+LDM integrated dataset resulted in biologically heterogeneous populations of cells along what appears to be connected topologies.

**Supplemental Fig. 1:**
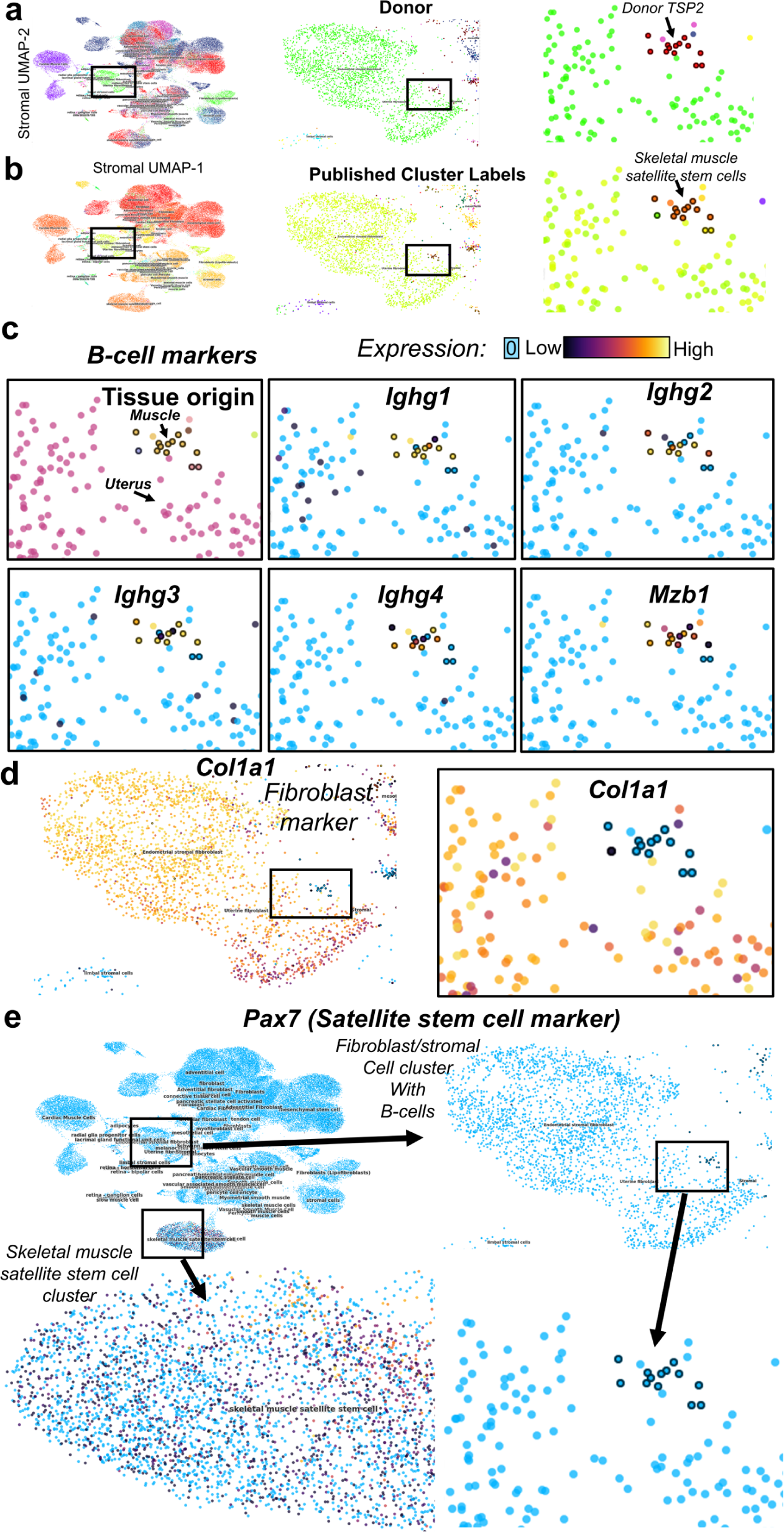
Example errant integration in the *tabula sapiens* atlas with mislabeling due to application of Class-2+LDM integration and tissue specific prior expectations. **a**, UMAP coordinate showing the *tabula sapiens* stromal compartment, colorized by donor. We focus on donor TSP2. **b**, UMAP colorized by the custom annotations provided by tissue experts; the highlighted subset is the region of interest with the donor of interest highlighted; these droplets were labeled as muscle satellite stem cells, while other cells within this topology were largely labeled as endometrial or uterine fibroblasts. **c**, The tissue of origin shows the droplets of interest were from a muscle sample, likely explaining their label of muscle satellite cells. Colorized expression for B-cell marker genes shows that these droplets are consistent with B-cells, yet are topologically integrated with endometrial/uterine fibroblasts. **d**, The droplets of interest were also negative for the fibroblast marker *Col1a1*. **e**, Droplets of interest were also negative for the satellite stem cell marker *Pax7*, despite many of the cells within the larger satellite cell cluster being positive for this marker.

In addition to the risks of errant labels, when relying on labels from prior publications, we are also at risk of label incompleteness. We hypothesized that if an *unknown* difference in cell-state existed across three different batches, commonly employed benchmarking metrics may reward for erasure of these non-annotated differences. To test this hypothesis, we partitioned a dataset (10k human PBMCs) randomly into thirds, and introduced high, medium, and low brackets of transcription randomly in a batch confounded manner, to a subset of genes from pertinent signal transduction and immune related pathways (**Supplemental Fig. 2a-c**). Note that these “cell-state spike-ins” were not generated by simulation, but were instead templated by three brackets of underdispersed genes, thus accounting for any concerns over simulation not matching biologically derived distributions (**Supplemental Fig. 2b**). Furthermore, because these genes were underdispersed, and therefore not included by the feature selection algorithm for the first round clustering results (state not-annotated), there was no data leakage between partitioning and cell-state spike-ins.

We sought to understand the impact of labeling or not labeling batch confounded biologically meaningful signal. If an experimenter were unaware of these differences in cell-state across batches, and used only the parent population’s label (as identified before *in silico* splits and signal spike-in), standard benchmarking metrics (kBET-cell-type, average silhouette width (ASW), cLISI) rewarded for erasure of biologically meaningful signal (**Supplemental Fig. 2d**), showing that perfect knowledge of cell-identity, including all biological differences between datasets, was required for accurate benchmarking with metrics that utilize user-defined cluster labels, resulting in a benchmark that rewards erasure of the unannotated batch confounded signal.

Under this paradigm, we examined standard benchmarking metrics (i.e.: ARI, kBET-cell-type, cLISI, etc, 27 metrics in total) comparing apparent method performance when this batch confounded signal was or was not annotated by the experimenter with respect to the reference labels (**ED Table 2**). Indeed, 14 of 27 metrics were statistically un-correlated when comparing results based on reference labels that did annotate state-differences vs results tabulated where state-differences were not included in reference labels (**ED Table 2a,c**). A further 13 of 27 metrics showed significant and consistent changes in means dependent on perfect annotation of cell-state (**ED Table 2a,e**). This left only 13/27 metrics that were not directly impacted in either of these manners, but these metrics were each batch normalized or agnostic, yet also hinged on the assumption that reference labels should carry no additional within group variation (i.e. ASW) (**ED Table 2g**).

We also assessed the impact of batch confounded cell states when applied only to a single cluster that we split it into three, through underdispersed templated cell-state spike-ins as described above. Given that most of these metrics provide global averages, across a dataset, these effects were blunted, except in the case of metrics looking for the region of “worst performance” such as minimum cluster ASW and maximum kBET-cell type (**Supplemental Fig. 2e,f**).

Additionally, the labels used by typical benchmarks were arrived at by humans using their preferred pipelines and analytic preferences. Therefore, using the labels generated by these individuals sets their given approach as equivalent to the ground-truth. This also suggests the possibility that processing steps taken during generation of reference labels can impact the apparent performance of a batch-correction method independent of accuracy, in a manner that rewards similarity to the approach used to generate refence labels. To test this, we compared results based on cluster label assignment approaches: 1) Louvain modularity based on a kNN built from the cell-cell Euclidean distances of the cell-cell Spearman correlation matrix; 2) Louvain modularity on a kNN built from Euclidean distances based on a dimensionally reduced version of the dataset (the first 30 principal components).

Indeed, some commonly used metrics including cLISI, entropy, minimum ASW were significantly different based on the processing method used for generating the reference labels, systematically favoring similarity over the processing method used for reference label generation (**Supplemental Fig. 2g**; **ED Table 2**). Conversely other metrics did not show a global mean shift, but in some cases, changing the clustering pipeline used for reference labels created un-correlated metrics (particularly for the kBET-cell-type, and several others that measure within-cluster spread, which yielded non-correlated results, simply by changing this minor step in processing when reference labels were generated) (**Supplemental Fig. 2g**; **ED Table 2**). While not the focus of this work, the change in benchmarking metrics dependent on latent dimension mapping fits with recent findings where operating on low dimensional projections can yield representations that are not reflective of true cell-cell similarity/distance^59^.

**Supplemental Figure 2:**
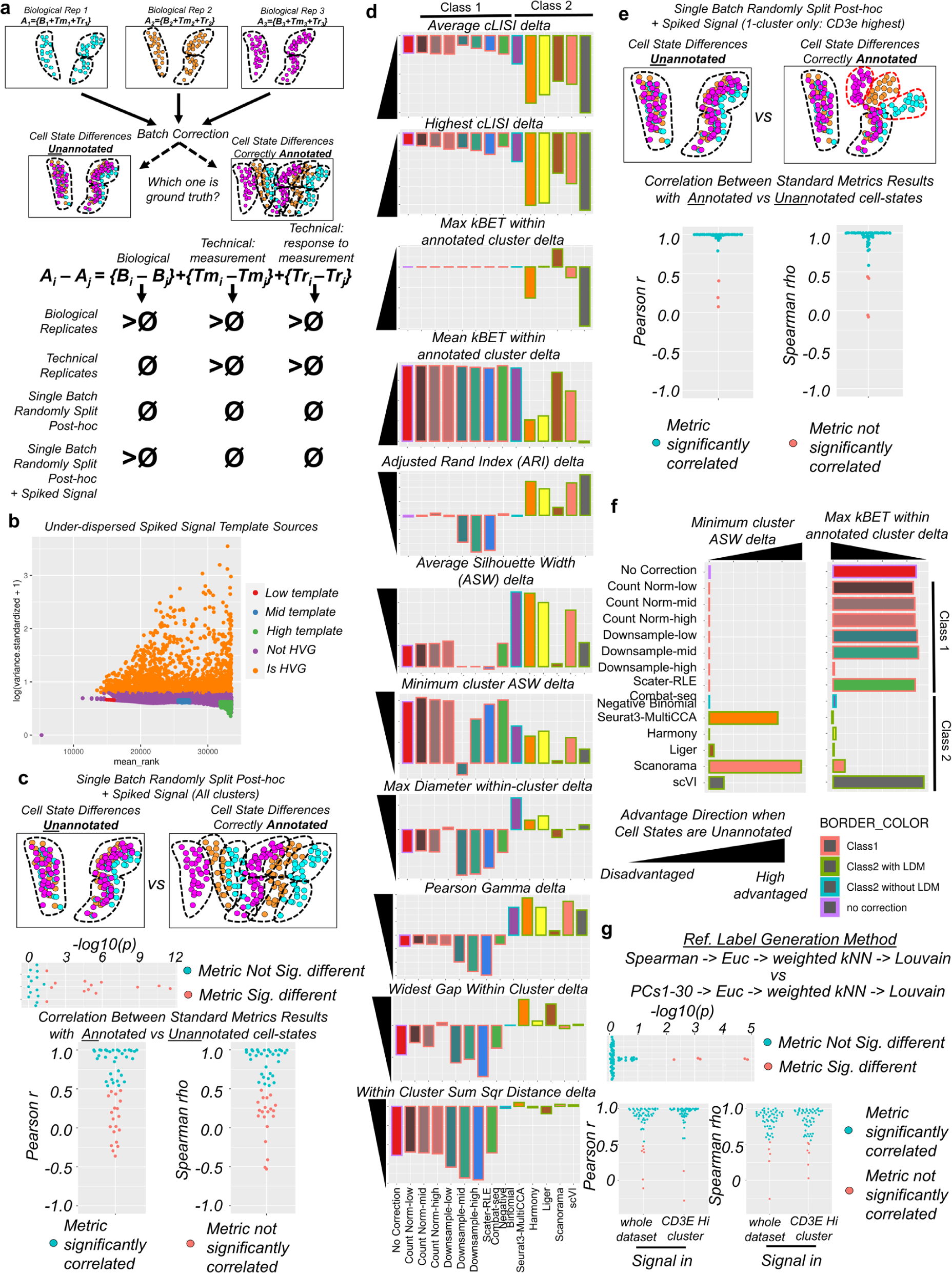
Application of standard ML metrics for benchmarking batch-correction that use externally provided labels assume perfect knowledge of biology by the experimenter. **a**, A schematic example of three batches that differ broadly from each other based on a shared source of variation, but the source of difference may fall into the set of biological functions, technical functions associated with measurement (T_m_), or the biological response to processing, which can also be considered a technical artifact, and not of interest (T_r_). We examined the effect of using different labels as ground-truth that either keep or discard this source of variation between batches. **b**, A scatter plot of rank-mean expression and log-standardized variance, colorized by dispersion status. Using human PBMCs, we used under-dispersed genes as a “reference bank” in lieu of simulation to ensure that distributions match biological distributions; these low-/mid-/high-reference banks were used as templates to spike-in cell-state signal in all annotated clusters. **c**, We compared the effects of using the cluster labels from pre-cell-state spiked signal and labels that were “split” based on the spiked-in immune state differences. Significant differences and correlations of standard ML and batch-correction metrics comparing the without-cell-state labels or with-cell-state-annotated labels as reference. In many cases, results were significantly different or uncorrelated (salmon color), indicating that these metrics frequently require perfect knowledge of the underlying biology, otherwise the unannotated sources of variation will dramatically impact results. **d**, A selection of commonly used metrics examining the difference between using cell-state-annotated and cell-state-unannotated reference labels shows that many of these metrics systematically advantage status quo Class-2 methods. **e**, Similar results are found when cell states differ only in a single cluster (MAPK/ERK/immune signaling spiked into T-cells only), but results are notably blunted because many of these methods average over the entire dataset. **f**, However, metrics that examine the worst performing cluster show a similar pattern as with **d**. **g**, Significant differences and correlations between when reference labels were generated using a LDM pre-processing step vs using high-dimensional Spearman correlation pre-processing.

While we agree with the broad consensus that these metrics in fact capture *some* attributes of preservation of human-annotated biological signal, the above examples clearly demonstrate that these metrics can also incentivize erasure of non-annotated attributes of cell-identity, and can incentivize consistency with the pipeline choice during generation of reference labels. Lastly, others have also used less supervised methods to annotate conservation of biological signal, such as conservation of cell cycle or highly variable genes^60^. These metrics benchmark preservation of *within-batch* biological variation rather than *across-batch* differences: an important, albeit orthogonal question to the one investigated here.

### Supplemental Note-2: How can we understand what is biological and what is technical when these are typically intrinsically confounded?

We began attempting to develop a functional framework to describe the often intrinsically confounded attributes of biologically meaningful variation and purely technical variation. In our framework (**Fig. 1**), we accept the premise that our observations must belong to one of these two classes: biological, B and technical, T. Notably however, within the technical set, others have previously described an interaction term in which the biological system responds to processing^21, 22^. We therefore split the technical set into two non-overlapping subsets that combine to encompass all technical effects T: 1) Those that are strictly technical effects of measurement and processing (T_m_), which may include things such as enzyme activity, buffer composition differences, GC content, internal priming, etc; and 2) those that capture the biological response to processing (T_r_) (**Supplementary Fig. 3a,b**). Others have previously combatted T_r_ using actinomycin-D to prevent active transcription as early as possible in processing^21, 22^.

**Supplementary Fig. 3:**
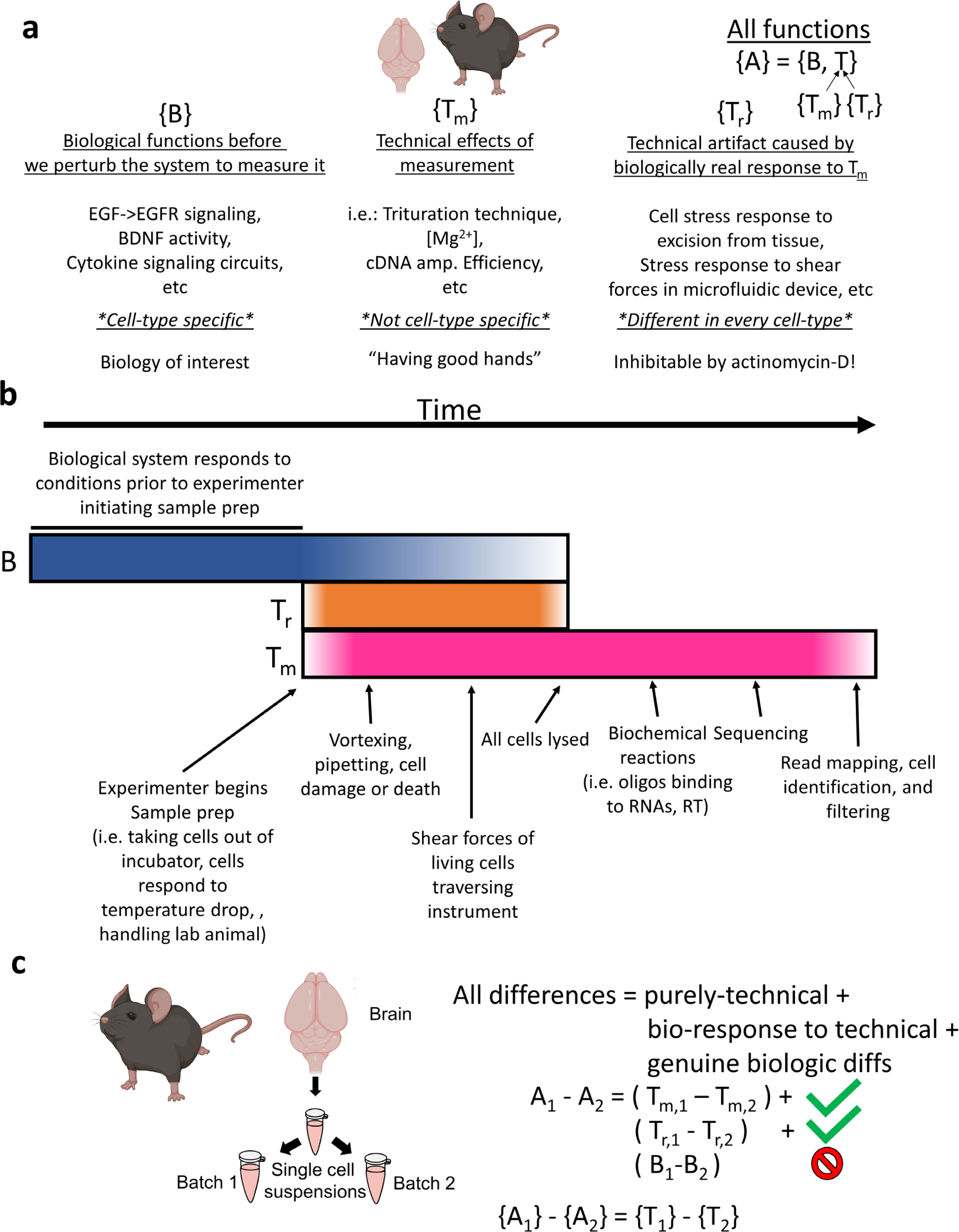
Conceptual basis for set-theoretic framework to classify biological and technical functions. **a**, Schematic description of the sets of effects at play in a typical scRNAseq experiment. **b**, A schematic timeline of when each set of effects will be acting in a manner that alters the observed count matrix. **c**, Using this functional framework, it becomes apparent that technical-replicates do not differ in the biological set, meaning that all difference in the observed matrix can be traced back to the technical set.

With this framework in hand, it became apparent that if we can obtain samples which are known to *only* differ by technical effects, and our original biology of interest is identical (i.e. true technical-replicates – not biological-replicates), then we can directly measure the downstream effects of the set of functions contained within T (**Supplementary Fig. 3c**).

We therefore used three sets of technical-replicates in which two pipette draws from the same exact single cell suspension were run in parallel, as these must only differ biologically by random Poisson sampling of the samples taken from the single cell suspension. We utilized 10x’s technical-replicates all within their version 3 chemistry: Mouse brain, mouse heart, and human PBMCs. Note that the results described here may be different for other chemistry versions and other technologies.

As discussed in the main text (**Fig. 1**), we found 6 categories of genes that appeared to be contained within T_m_ that were differentially expressed in a high percentage of clusters that were also highly conserved across tissue. These included genes encoding nuclear targeted proteins or lncRNA such as mRNA processing machinery such as capping and splicing, nucleosome binders such as High Mobility Group Box genes (HMGBs), which are both nucleoproteins and an extracellular stress related cytokines^61^, and the lncRNA MALAT1. We also observed genes relating to the other large structural components of the cell as members of T_m_ that were differentially detected across technical-replicates in a high fraction of cells including ER associated ribosomes and mitochondria (**Supplementary Fig. 4,5**). Interestingly, in the two solid tissue preps we examined, hemoglobin and related genes were also differentially detected, yet in PBMCs where red blood cells (RBCs) are physically separated from the cells of interest during sample preparation, these genes were not differentially detected (**Fig. 1g**, **Supplementary Fig. 6**). This suggests that RBC depletion may help technical consistency even in solid tissue.

**Supplemental Fig. 4:**
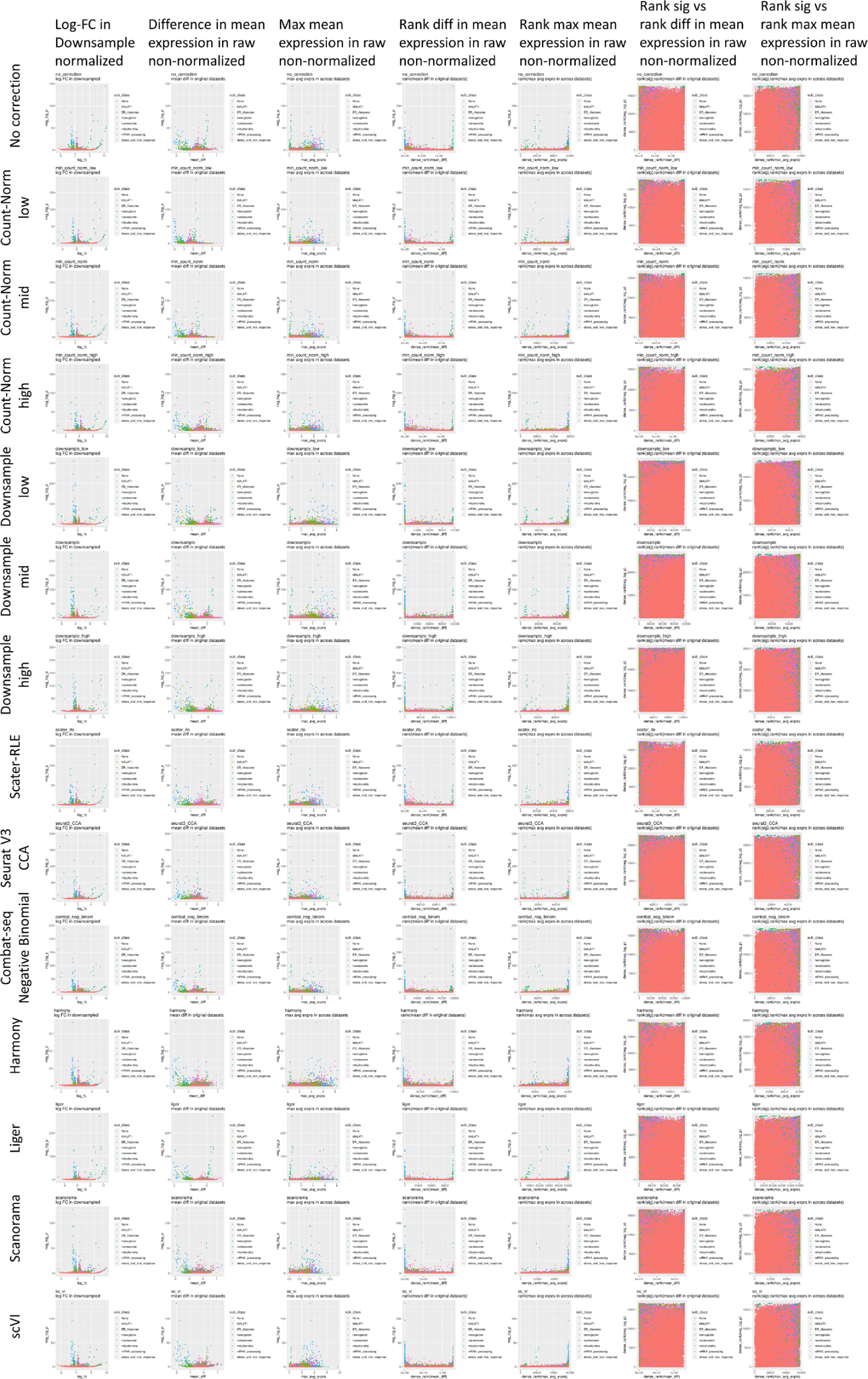
WCAB volcano plots and absolute expression differences in mouse brain technical-replicates. In the first column, volcano plots showing average log fold change (FC) (x-axis) and -log10(p-value) for differential expression within clusters across batch (WCAB) colorized by technical effect sub-class. This column’s average log-FC are those calculated on the downsampled datasets quantifying the difference in relative composition rather than absolute counts; these were used for DE analyses. We assessed whether this was a technical artefact of these genes being highly expressed by examining whether *all* highly expressed genes were detected as DEGs. As shown by the subsequent columns, while higher expression led to higher *sensitivity* and the *ability* to detect DEGs, high expression on its own did not determine significance as shown by the many highly expressed genes that traversed the spectrum of significant and non-significant. This points towards a change in measured composition within these pathways, rather than a general trend attributable to differences in total UMI reads.

**Supplemental Fig 5:**
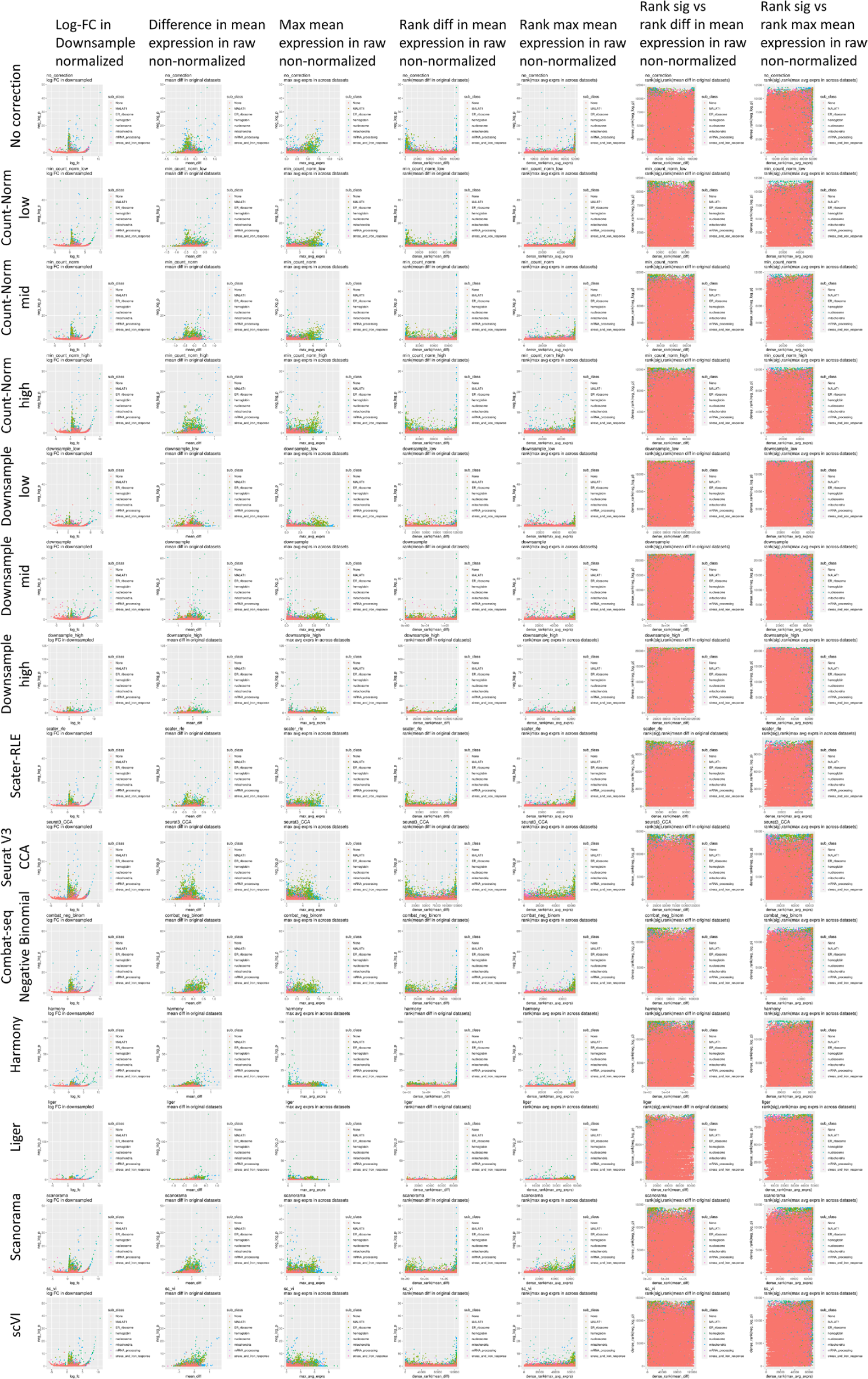
WCAB volcano plots and absolute expression differences in mouse heart technical-replicates. See legend for Supplemental Fig. 4.

**Supplemental Fig 6:**
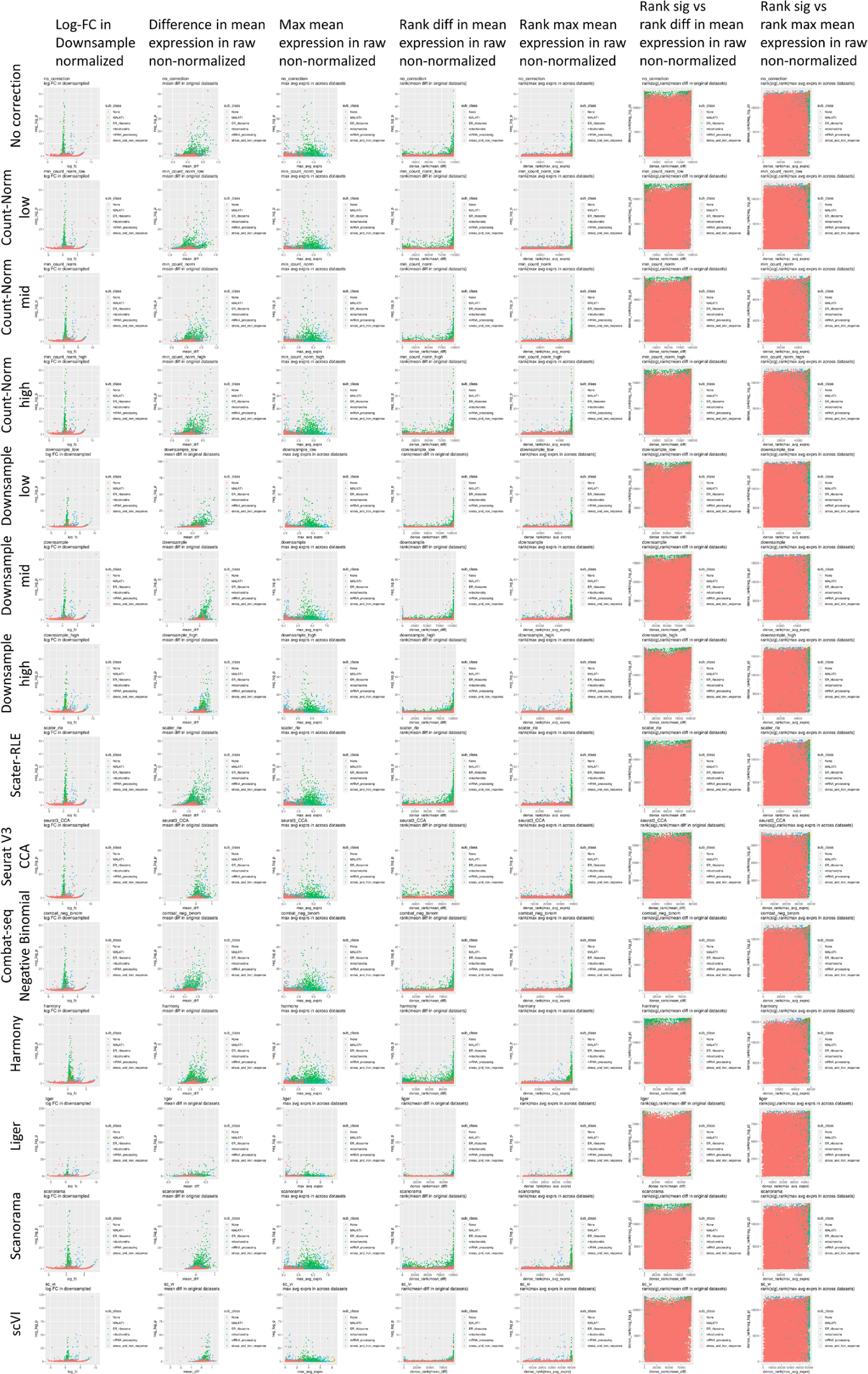
WCAB volcano plots and absolute expression differences in human PBMC technical-replicates. See legend for Supplemental Fig. 4.

The fact that we observed genes that were highly expressed in each technical replicate that were not differentially detected indicates that the simple property of high expression was not the *cause* of this differential detection (**Supplemental Figs. 4-6**). However, as with all statistics, when dropout from shallow sampling is a factor, greater sampling depth increases the power to detect differences. In other words, greater expression levels increase power and sensitivity (due to a greater probability of detection), but were not the single determinant of significant differences.

While the above findings should primarily be taken as measures of the technical effects for 10x’s Chromium V3 chemistry, this approach can be taken to further describe technical effects in other chemistries and platforms. Additionally, more work should be done using technical replicates across other tissues and species to build upon these results.

### Supplemental Note-3: Proof 1: What are the theoretical implications of the functional framework for Class-1, Class-2, and Class-3 correction algorithms?

To extend our experimentally determined descriptions of the underlying elements of T, we further sought to understand whether this these different sets may or may not be altered during the transformations applied to the observed data using different batch-correction approaches. We first begin with the definitions required to understand how different classes of algorithms operate.

#### Given the definitions

Dimensionality parameters:

1) *b*: the number of batches
2) *i* and *j*: Iterators that operate over *b*
3) *n*: the total number of cells observed
4) *n_i_*: the number of cells in a given batch *i* (with *i* ranging from 1 to *b*)
5) *m*: the number of genes

Types of expression matrices:

1) ***S***: The initial *m* by *n* dimensional data matrix that can be thought of as the “base-transcriptome” for all cells.
2) ***D***: An *m* by *n* dimensional matrix that represents the ground truth transcriptomic data matrix, whose values can be generated through the application of all biologically derived functions in the set *B* (defined below); this is the matrix we are interested in measuring perfectly.
3) ***D*^*i*^**: The subset of the ground truth transcriptomic matrix that belongs to each batch *i*.
4) ***O*** : The observed data matrix across all batches, representing the empirically determined version of ***D***, after application of the technical effects introduced to make the observation of ***D***.
5) ***O*^*i*^**: The set of observed data matrices, each of *m* by *n_i_* dimensions for each batch *i*, that can be described as the output of the functions contained in *T* and *B*: ***O*=*A*(*S*)**.
6) ***D*_*c*,*g*_**: A subset of cells (from the *n* columns in ***D***) and or genes (from the rows of ***D*)** that are assumed by the experimenter, or selected by an algorithm, to be invariant across a subset of batches.
7) ***C***: An *m* x *n* matrix that represents the batch corrected data, intended to accurately represent ***D***.

Sets of functions to transform expression matrices:

1) *B*^*i*^: The theoretical set of biologically encoded functions, for each batch *i*, that reflect the complex non-linear biological processes that transform the baseline expression levels represented by *S* to the states the cells are in when they are profiled represented by ***D***. That is, *B*^*i*^(***S***) = ***D***^*i*^. These are the functions we are trying to learn by performing the experiments giving rise to the cells being profiled. These functions encode the cell-lineage, -type, and -state prior to sample processing.
2) *T*^*i*^: The set of technical effects for each batch *i* that impact the observed measures of the true transcript.
3) 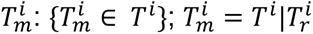 : The subset of *T*^*i*^ technical effects that are not encoded by the cells’ stress or other physiology derived responses to processing (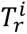 defined below), but are technical effects from the act of measurement. These include factors such as transcript biases in PCR amplification, internal priming while using an oligo-dT primer, the effects of minor variations in buffer composition due to pipette error, diffusion of small transcripts out of the cell after fixation for methods such as split-seq^62^, read mapping errors, and more.
4) 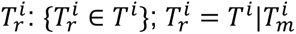 : The subset of *T*^*i*^ technical effects that are biologically real, caused by transcription, or other physiological changes, but are attributable to the cells’ response to sample collection and preparation. This may be cell stress, but can also include of forms of processing induced transcription, or transcript degradation, etc. This can also be interpreted as the set of functions that capture the interactions between 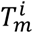 and *B* (linear and nonlinear alike). Because our central focus as biologists is to understand *B*, rather than how our processing affects the cell, these are typically viewed as uninteresting, and in some cases have been prevented through the use of actinomycin-D, preventing the application of these functions by inhibiting transcription during sample processing^22^. However, because these are processing artifacts, they will not be of biologic interest since they reflect the experimenter’s action on the basal state system *B* that we are trying to study. Notably, others have already reported that the cellular responses to differences to processing *T*_*r*_ differs per cell-type^22^.
5) *A*^*i*^ : The union of all biologically real functions in *B*^*i*^, whose outputs are passed to the set of functions contained in *T* for each batch i: {*T*^*i*^, *B*^*i*^}. Given *B* represents the biological functions giving rise to the transcriptome up until technical artifacts are introduced, these functions are composed as *A*^*i*^(*S*) = *T*^*i*^ (*B*^*i*^(*S*)).
6) *T*^−1^: The true inverse function of *T*. This can be interpreted as the “perfect” batch correction algorithm.
7) 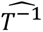: A batch correction function that takes as input ***O***, attempting to correct for the effects induced by functions in *T*, returning a corrected m x n transcriptome (***C***) that is intended to accurately represent ***D***.

With the above definitions that describe the series of processes that impact baseline states of cells in response to experimental conditions and the isolation and processing of the cells for sequencing, we can categorize the different algorithms for correcting for batch effects into 3 classes:

1. **Class 1** 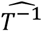 batch correction algorithms: Algorithms that attempt to directly model or mask the technical effects encoded by the functions in *T*. While these algorithms require an explicit understanding of *T*, they are not selection of a subset of cells and genes that are biologically equivalent across batches as these algorithms do not make any assumptions on biological equivalence across subsets of batches.

a. Model *T*^−1^ without assumptions on equality or inequality of *B* or subsets of *B*;
b. The parameterization of 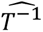 must be known or learned *a priori*, and then for a given dataset, parameter estimates are determined to come up with the specific 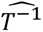 to apply on the dataset;
c. Some of the parameters may be convolved with the biological signal, so that some loss of the true biological signal may happen, but only to the extent that these signals covary.
2. **Class 2** 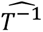 batch correction algorithms: Algorithms that attempt to calculate the inverse of T by learning the relationships between 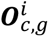 and 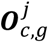 across all batch *i* and *j* comparisons. These algorithms do not explicitly model the functions in *T* and *B*, but rather they identify subsets of genes and cells between batches in an unsupervised way, and assume the 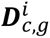 matrices are equivalent to the 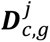 and thus by definition will remove differences between *A*^*i*^ and *A*^*j*^. As a consequence of this strong equivalence assumption, if the 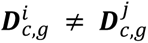, then these batch correction procedures will condition out true biological signal (see proof below). Identify regions (cell clusters) of matched topology, assume within those *B*^*i*^ and *B*^*j*^ are equal, and model *T*^−1^ as taking you back to a common *D* among these equal regions.

_a._ Because *T*^−1^ is determined by looking at biologically equivalent regions, the *T*^−1^ will be fit in a way that forces the biological signals to be the same, even if they are in fact different;
b. To see this, suppose 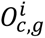 and 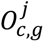 are determined to be comprised of *c* cells of the same type and state across the set of *g* genes, and then assumed to reflect the same biology, but suppose the true underlying state is such that we 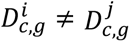
c. **<u>Proof 1</u>**: With 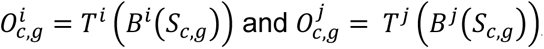, the application of the inverse would give us that:

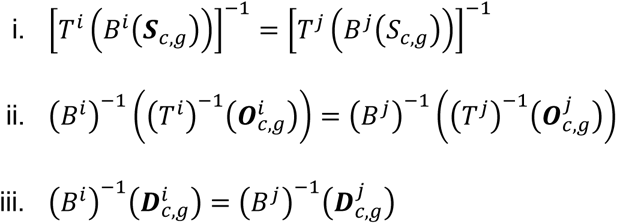

but in order for this last result (iii) to hold, given that contrary to the assumption of equivalence we have that 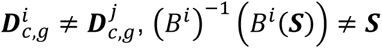, so that the inverse fit by the algorithm would be with respect to a different biological function than that which gave rise to 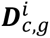 (and 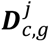);
d. Unlike class 3 approaches which are semi-supervised, in class 2 algorithms one is letting the algorithm pick the regions that are homologous and then acting to minimize the differences between them. This is frequently performed using a mutual nearest neighbors (MNN) algorithm.
3. **Class 3** 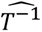 batch correction algorithms: Similar to class 2 algorithms, but perform a semi-supervised selection of c and g, cells and genes, which are assumed to share the same biological cell-lineage, -type, and -state. In practice, this takes the form of manual annotation of cell-identity labels in various datasets that are used to train an algorithm that performs an end-to-end learning of the transformation function to merge the shared labels for *c*.

For the class 2 and 3 algorithms, distance functions that measure the distance between the “matched” areas of topology are used to learn the inverse function by minimizing the distance between the areas of matched topology. The precise implementation of these distance functions vary by each algorithm. For example, Harmony performs a linear model correction on the batch factor, minimizing the least squares of the results of fuzzy clustering of the principle components identified in a PC analysis of the data restricted to the genes selected by a feature selection algorithm, with cell equivalent locations identified by gradient descent in the linear model’s correction^10^. LIGER uses non-negative matrix factorization as the dimensionality reduction technique and then again uses least squares to find the minimum distances, weighting for both full dataset reconstruction accuracy and minimized across dataset distance^11, 63^. Part of Seurat’s IntegrateData function includes simple subtraction of the difference between mutual nearest neighbor (MNN) “anchor” cells across datasets, 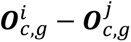, which they call the ‘integration matrix^64^.’ While the specific pipeline implementation of these local distance functions, and their implementation of 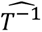 differs across methods, they are conceptually similar: estimate the inverse of *T* by minimizing the local distance between the regions of the dataset that the algorithm assumes are biologically equivalent, and apply a locally smoothed transformation that minimizes those distances.

It is worth noting the relation that the above has to traditional batch correction using linear models to regress batch using bulk data. In large part, these also assume biological equivalence across batch, like Class-2 algorithms in the context of scRNAseq; however they rely on the balancing of known biological factors across batch (age, sex, disease/treatment status, etc). This means that *B*^*i*^(*S*) ≈ *B*^*j*^(*S*) due to supervised allocation of *B* across *B*^*i*^ and *B*^*j*^symmetrically, while an additional offset for each batch is therefore likely to be caused by technical effects. However, as shown by the proof above, this would not be a safe assumption if each biological replicate were its own batch, because we would be unable to rely on the assumption of mean equality across batch due to balancing biological effects. With the advent of multiplexing single-cell omics however ^2, 62, 65, 66^, these algorithms may be modifiable in the future to rely on the same supervised balancing of *B* that traditional bulk batch correction algorithms use. However, as with traditional batch correction, this relies on high quality balancing of effects, and could also result in misleading results if important biological factors are not balanced across batch, whether those biological factors are known to the experimenter or not.

### Supplemental Note-4: PMD characterization and proof of invariance to number of clusters

#### PMD metric definition

##### Definitions

- O_actual_: An m-by-n matrix of observed cell counts, across the m identified clusters and n batches.
- E_actual_: An m-by-n matrix of expected cell counts under the assumption (or hypothesis in the context of a statistical test) that the distribution of clusters is the same across batches. This m-by-n matrix is tabulated in a manner similar to the “expectation matrix” in the context of a X^2^ test-of-independence.
- O_max_: An n-by-n matrix with the diagonal set to the number of cells in each of the *n* batches and all other elements of the matrix set to 0; that is, for each batch (i) with i = 0..n, O_max_[ i,i ] = batch_size[ i ] and O_max_[ i,j ] = 0 when *i* ≠ *j*. O_max_ represents the theoretic maximum level of asymmetry across batches, under the hypothetical that each batch was comprised of a single cluster that contained all of that batch’s cells.
- E_max_: An n-by-n matrix of expected max cell counts across the n batches under the assumption of each batch being comprised of a single cluster.

Using the observed and expected contingency matrices (similar to X^2^ test-of-independence), we found that the sum of the absolute value of the difference between expected and observed matrices were equivalent, regardless of the size of *m*, the number of clusters observed within each batch (Σ|**O**_actual_ – **E**_actual_|), so long as relative cluster composition by batch was conserved (**ED Fig. 2a,b**). A proof that the PMD metric has the property of invariance to the number of clusters is provided in a subsequent section (**Proof 2**).

This equivalence property allowed the creation of a hypothetical contingency table, in which the entirety of each dataset fell into a single cluster that was specific for this batch (**O**_max_). **O**_max_ can be thought of as, what a contingency matrix would look like, given no overlap in cluster composition by batch (given the input batch sizes), with only a single cluster per batch. Thus creating a hypothetical contingency matrix that maximizes the possible asymmetry by batch, yet mirrors batch sizes of the input datasets. Because there is no dependence on the number of clusters within batch on the outcome of Σ|**O** – **E**| (**ED Fig. 2a**; **Proof 2**), we can calculate this for the maximum asymmetric hypothetical contingency matrix (Σ|**O**_max_ – **E**_max_|) as well, and take the ratio of the actual difference relative to maximum possible (**ED Fig. 2b**):

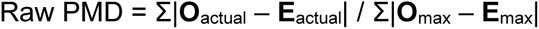

This yields the Raw PMD metric, that is bounded between 0 and 1, that quantifies how similar cluster composition is across batches relative to the version of the input that maximizes the possible asymmetry by batch. A result that equals 0 indicates that the input (**O**_actual_) showed batches where cluster abundances were *exactly* equivalent (**O**_actual_ ≈ **E**_actual_; **ED Fig. 2c**). Note that because of random Poisson sampling, it is likely that a real-world dataset would only *approach* zero, but that there would be fluctuation above zero even when there was no real difference – simply attributable to random sampling.

When no clusters appear in more than one batch, a dataset would maximize the Raw PMD value at 1 (**ED Fig. 2d**). Because of the previously mentioned equivalency of Σ|**O** – **E**| regardless of the number of clusters, Σ|**O**_actual_ – **E**_actual_| will be equivalent to Σ|**O**_max_ – **E**_max_|, thus making the ratio equal to 1. Unlike the lower bound however, Poisson sampling under this scenario does not yield a distribution because all clusters have a 0 percent chance of appearing in multiple batches.

#### Characterization of the Raw PMD metric and creation of the final PMD metric

To characterize the linearity and other properties of the Raw PMD function (**ED Fig. 2**), we simulated two batches containing 10 clusters each, with variable numbers of cells in each batch as noted in the legend of **ED Fig. 2e** (10 iterations each). In combination with varying the number of cells per batch, we also incrementally changed the amount of overlap in the two batches. This was done by decreasing the number clusters that were in batch 1 that also appeared in batch 2. In all simulations, each batch still maintained 10 clusters, but progressively increasing the number of clusters that were specific to a given dataset. The most dissimilar scenario therefore contained a total of 20 clusters (2 batches, 10 clusters each that were all batch specific). The most similar scenario contained a total of 10 clusters (2 batches, 10 clusters each that were all shared across both batches evenly). This characterization is implemented in the do_full_pmd_characterization() function in the PercentMaxDiff R package that we released (https://github.com/scottyler89/PercentMaxDiff).

In all simulated cases, the relationship between the number of non-shared clusters was linear with the Raw PMD value (**ED Fig. 2e**) as calculated with the equation in **ED Fig. 2b**.

Interestingly however, the slope and intercept of these lines appeared to depend on the number of cells in the datasets. We plotted histograms (**ED Fig. 2f**) to characterize the distributions of Raw PMD at the intercept of each line in **ED Fig. 2e**; this showed that when there was no actual difference in underlying cluster abundance (2 batches, 10 clusters total, all with equal relative abundance), the distributions of Raw PMD appeared to be Poisson distributed. This fits with the underlying discrete sampling errors that generated the cluster by batch contingency tables.

Importantly, the peak of these distributions (lambda parameter of the Poisson function) appeared to be dependent on the input dataset sizes, shown in different colors in **ED Fig. 2f**. This indicates that the Raw PMD function, while being bounded between 0 and 1 and equivalent under *idealized* circumstances regardless of the number of cells or clusters, it is affected by random Poisson sampling. This is particularly true when underpowered, as noted by the higher Raw PMD when comparing two batches with only 100 cells each, but 10 cell types. At this point, the background noise from Poisson sampling increases the observed Raw PMD, even when the underlying distributions in cluster abundance across batch are equivalent (**ED Fig. 2f**).

An ideal metric would allow all input datasets, regardless of number of clusters or number of cells, to scale linearly with the degree of overlap between the input datasets. We therefore sought to better characterize any dependence of the Raw PMD on 1) the number of clusters and 2) the number of cells. Specifically, for each dataset in these simulations, we performed Poisson sampling Monte Carlo simulations under the hypothetical that there are no differences in relative abundance of clusters by batch (as with the Y-intercept of the lines shown in **ED Fig. 2e**). This allowed an empirical observation of the input dataset’s Y-intercept under the “unpatterned by batch” scenario, while also capturing the effect of the input sample sizes and number of clusters given that these are maintained in the simulations.

This is similar to calculating the expected matrix, as with a X^2^ test-of-independence, but through simulation that matches the underlying cluster abundances and number of cells rather than just calculating the *idealized* expected matrix; this allowed the characterization of how Poisson sampling alters the Raw PMD when the underlying cluster distributions by batch are equivalent, but matching the input batch sizes. Using this background, we calculated the observed Raw PMD for each Monte Carlo simulation, under the hypothetical of no-pattern by batch, and fit the above described Poisson distributions, calculating the lambda parameter using the fitdistr function from the R library MASS.

Using 100 null datasets of each given input in the prior simulation (**ED Fig. 2e**), we observed that the lambda parameter distributions showed a clear dependence on both the total number of clusters and the size of the input datasets **ED Fig. 2g**. This can be intuitively understood as increasing the chance of random Poisson sampling away from *perfect* equivalence between batches, simply due to random sampling. Notably, this lambda parameter background is the intercept of the lines shown in **ED Fig. 2e**. Furthermore, due to the overall linearity of the shown lines, this intercept can be used to calculate the slope of the lines in **ED Fig. 2e** as well, thus allowing a modification of the Raw PMD, that incorporates and corrects for the background distributions caused by random Poisson sampling after fitting the distribution and calculating the best fit lambda parameter (**ED Fig. 2h**):

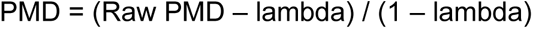

Indeed, applying this correction to Raw PMD, allowed the creation of the final PMD metric which shows no dependence on input size of the batches or total number of clusters (**ED Fig. 2i**). Thus, the final PMD metric has an upper bound equal to 1, whose lower distribution is centered around 0, and scales 1-to-1 linearly with increasing batch asymmetry.

#### Proof 2: Proof and example of cluster invariance with PMD equation

To prove whether the PMD is invariant to the number of clusters, let *C*_*i*,*j*_denote the clusters from *i* = 1. . *n* detected (*n* clusters in total) across the batches *B*_*j*_ for *j* = 1. . *b* (b batches in total). Let *n*(*C*_*i*,*j*_) denote the number of cells in cluster *C*_*i*,*j*_, and *n*(*B*_*j*_) = ∑_*i*_ *n*(*C*_*i*,*j*_) (the sum of the cell counts across all clusters identified in batch *j* – number of cells in the batch). Thus, the total number of cells over all batches and

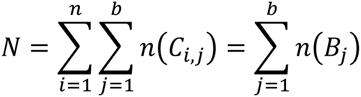

With this nomenclature, the PMD can be re-written as:

#### PMD Definition

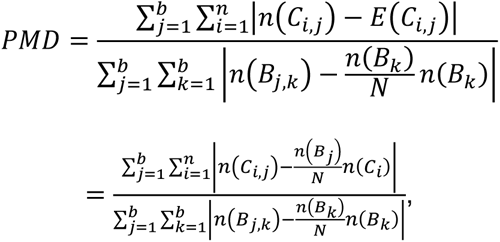

where *E*(*C*_*i*,*j*_) is the expected cell count for batch *j* in cluster *i* under the assumption of no association between batch and cell counts in the clusters, 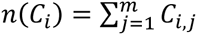 (number of cells in cluster *i* across all batches), and *n*(*B*_*j*,*k*_) = *I*_*j*,*k*_ · *n*(*B*_*j*_) with *I*_*j*,*k*_ = 1 when *j* = *k* and *0* otherwise.

For the denominator of the *PMD,* all of the terms are independent of the clusters and so by definition is invariant with respect to the number of clusters. We therefore need only to prove that the numerator is invariant to the number of clusters, given equal composition across batches.

To show that the numerator is also invariant, we need to show that the numerator delivers the same value for a cluster identified by one algorithm, as the value delivered for multiple clusters other algorithms may identified as representing that one cluster. The only constraint is that the proportion of cells across the different batches is preserved. Given this, without loss of generality, suppose that for cluster *C*_*i*_ we subdivide it into *l* subclusters:

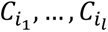

with the condition

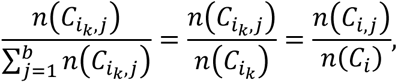

for *j* = 1. . *b* and *k* = 1. . *l*. We then have that

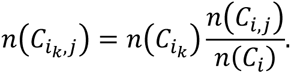

It then follows that

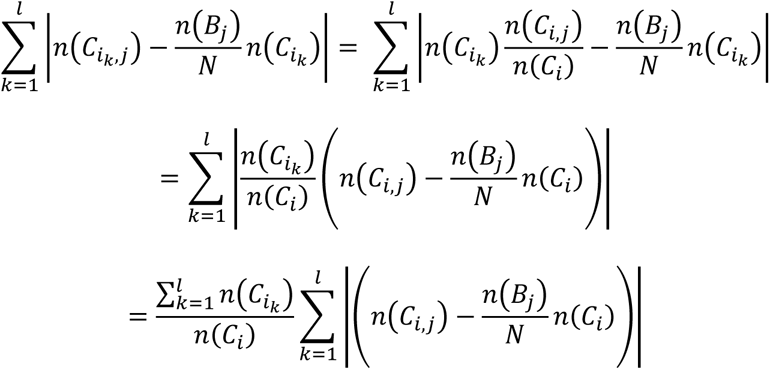

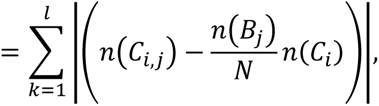

since 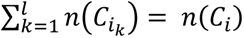. Thus we have that regardless of the number of clusters that subdivide any given cluster, so long as the proportion of cell counts across batches in the subdivided clusters is conserved, the *PMD* score will not change. Note also that subclusters in this context does not necessarily imply or require any hierarchical lineage relationships between clusters, but rather are simply representing different potential subdivisions of a dataset.

<u>Example</u>

Given two clustering results (R and R’), for two batches with 50% shared cluster composition, in which the shared cells are called either one cluster or two clusters, and third clustering result in which the relative composition was altered after cluster splitting (R’’), yielding slightly more asymmetry across batches:

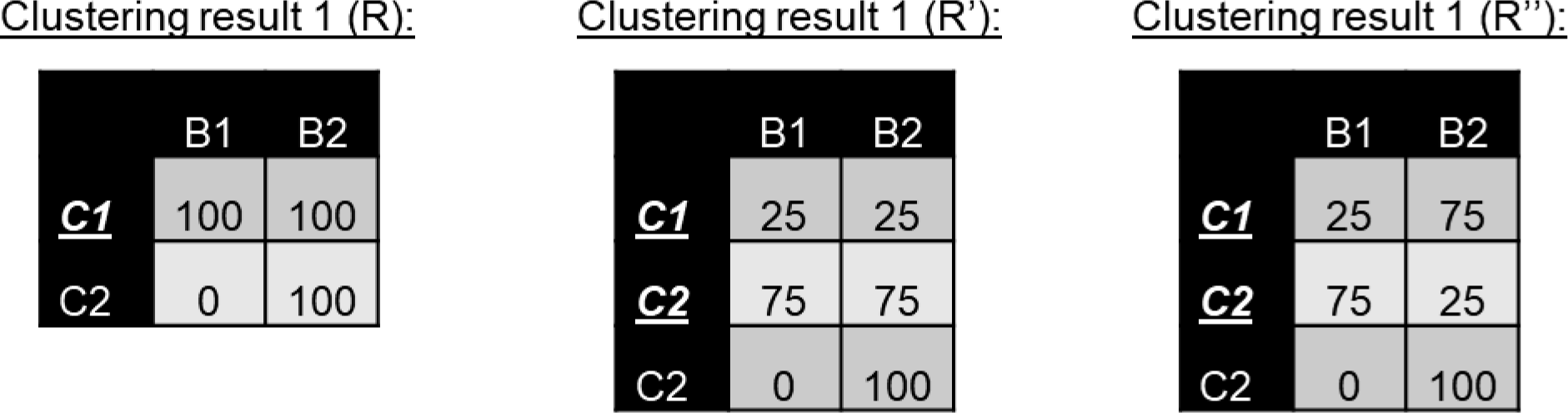

Yielding the expected matrices for the numerator (E_actual_):

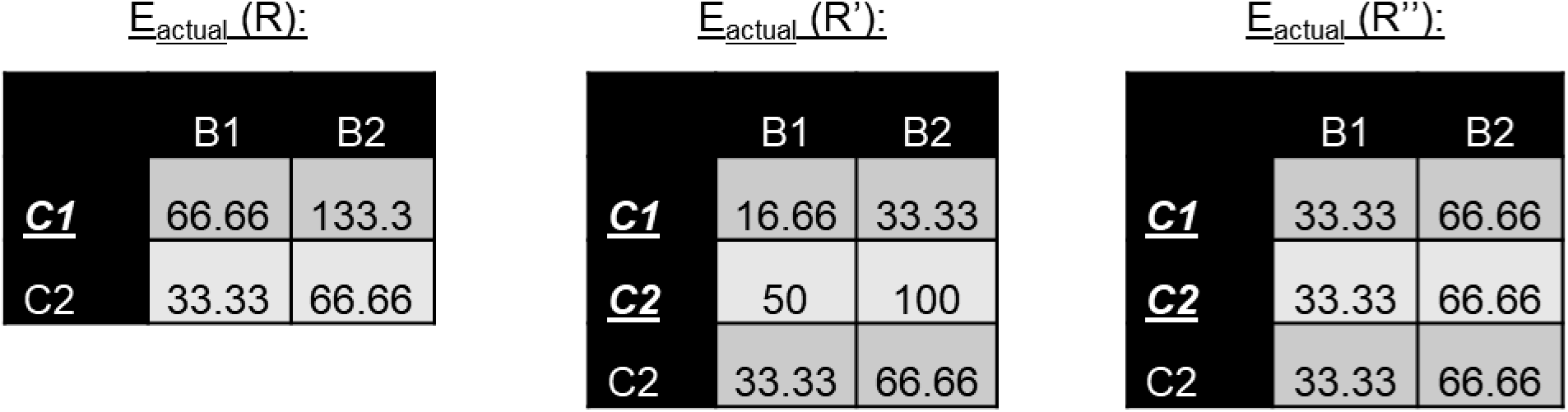

And the difference between O and E_actual_

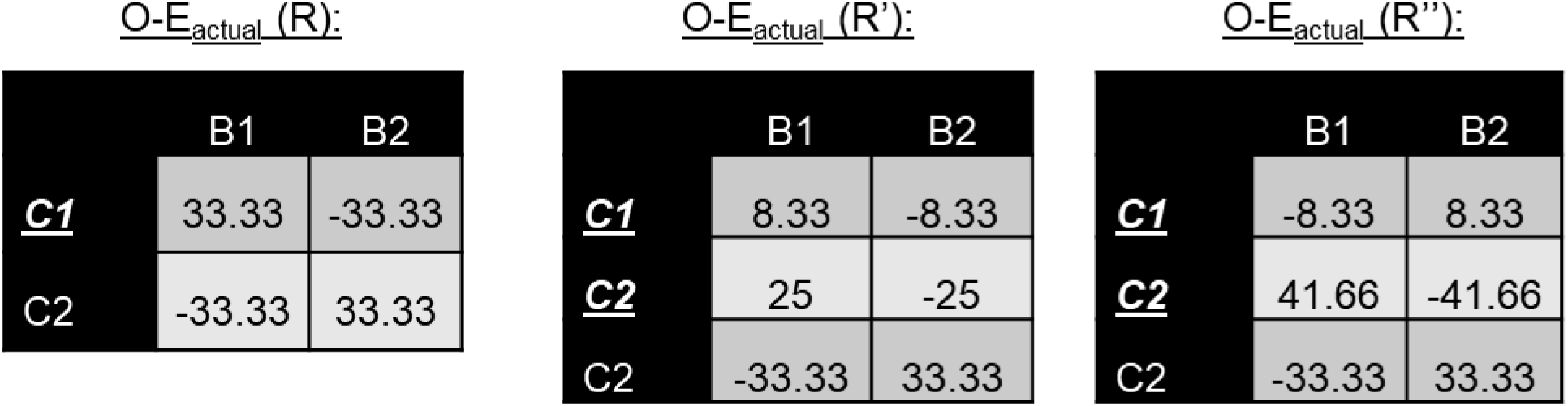

We can see that the numerator will not change between R and R’ because R:C1’s values are simply split between R’:C1 and R’:C2, and will therefore sum to the same value. However, this is not true for R’’ whose relative composition was changed with this subdivision.

When we complete the calculation of the PMD numerator, we observe:

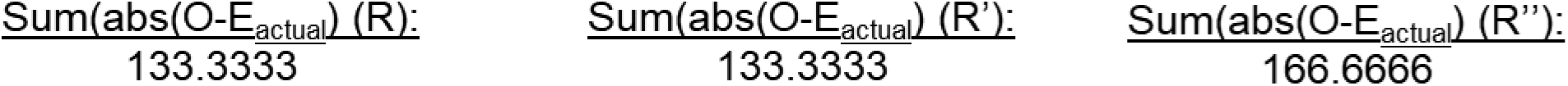

Showing that with R and R’, whose cluster composition was held constant, give the same PMD numerator value. Lastly, dividing by the universal denominator, as calculated independently from the clustering results from O_max_ and E_max_, yields the final PMD_raw_ metrics for these idealized examples.

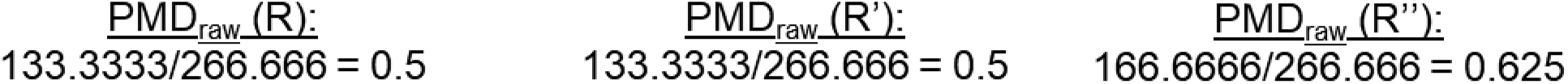

It is important again to note that actually observing ideal results from Poisson sampling is exceedingly rare, making it important to apply the lambda correction as described in **ED Fig. 2**, thus accounting for Poisson sampling noise.

#### Characterization of PMD, χ^2^ statistic, X^2^ statistic -log10(p-value), Cramer’s V, average inverse Simpsons index, and average count normalized Shannon’s entropy at the extremes of similarity and difference between batches

Analysis of contingency tables has been thoroughly investigated for many different purposes throughout history. Some of these methods include X^2^ test-of-independence, it’s - log10(p-value), Cramer’s V, average inverse Simpsons index, and average count normalized Shannon’s Entropy. However, in our specific use case, we sought a metric with very specific properties that could linearly quantify column similarity in a manner independent of differences in column totals, yielding the same result when relative row-wise composition is held constant between is the columns, even when row-wise dimensionality can be variable.

X^2^ and its corresponding p-value has been a standard metric for quantifying the independence of two sets of variables and their co-occurrence. However, in most cases the X^2^ or its p-value alone are the statistics of interest. Here however, with preliminary testing, we found some non-linearities, and non-robustness to differences in contingency table dimensionality and that differences in column totals impacted results.

Cramer’s V statistic is a previously published approach that builds on X^2^ to create a correlation-like metric based on counts of discrete classes in a manner similar to PMD^67^. A prior report indicated that this statistic demonstrated a high level of correctable bias, especially with contingency tables of dimension greater than 2×2, as will usually be the case in a cluster x batch contingency table.^68^ However, in initial testing, we found that the bias corrected Cramer’s V was still non-linear across the spectrum of similarity.

Diversity and information theoretic metrics can also intersect with this problem in the form of Inverse Simpsons index^69^ and Shannon’s entropy^70^. These metrics were originally designed to quantify diversity and information contained within strings respectively; however, since their introduction have become widely used across many fields of study.

To demonstrate specific scenarios in which the PMD metric provides an advantage over other metrics such as the X^2^ statistic, its -log10(p-value), bias corrected Cramer’s V (as implemented in rcompanion^71^), inverse Simpsons index (average across clusters), and count normalized Shannon’s Entropy (average across clusters), we designed a series of comparisons of all six metrics under different scenarios. An ideal metric would have all of the following characteristics:

1. **Robustness to differences in power**: An ideal metric would be unaffected by the total number of cells in the batches. For example, two datasets of equal size (1,000 cells each) that were sampled from the same distribution of cell clusters, should be identified as equivalent to each other. The returned metric should not be different if these datasets were 5,000 cells each. This allows direct interpretation on the degree of overlap between batches, regardless of their size.
2. **Robustness to differences in batch size**: An ideal metric for comparing how similar or different two batches are to each other in terms of their cluster composition, would be robust to differences in the size of two batches. For example, in the datasets used here, the dataset of 1,000 brain cells integrated with 10,000 brain cells should yield the same result as an integration of 1,000 brain cells with another dataset of 1,000 brain cells – in both cases, the desired metric would indicate that the batches come from the same cluster composition – unaffected by differences in batch sizes.
3. **Invariance to the number of clusters found within a given batch**: Batch-correction algorithms are expected to leave different levels of within-batch variation and across-batch variation, yet, with real-world datasets, we do not know the ground-truth number of clusters. However, when integrating datasets that came from the same single cell suspension, we know that cluster composition should be the same. Additionally, we can make reasonable assumptions that intestinal epithelial cells should not be found in the same clusters as neuronal cells. However, we cannot claim to know the ground-truth number of clusters within these datasets. To be able to benchmark batch-correction algorithms, which leave different levels of variation in a dataset, without making assumptions on the number of clusters, a metric is required that would report the degree to which cluster composition was similar across batches, while still allowing the number of clusters to vary, as this is determined by unbiased algorithms. For example, if two batch-correction algorithms were used, on intestine and brain datasets, and both found no batch overlap, but algorithm one found three brain derived clusters, while the other found 4 brain derived cell clusters, an ideal metric would indicate that in both cases, the methods were equivalent in yielding no batch overlap. As formally proven above, PMD is invariant to the number of clusters found, however, we will also show this with Monte Carlo simulations (**Supplemental Fig. 7,8**).
4. **Linearity across the spectrum of batch similarity:** For two batch-correction algorithms to be fairly compared to each other, there must be a consistent mathematical relationship between the degree to which batches are similar/dissimilar, and the reported metric. In the particular case where the relationship between the percentage of overlap in cluster composition and the reported metric is linear, the difference between having 50% and 60% of cells derived from the same cluster composition would be the same difference in the reported metric as 80% to 90% of cells coming from shared cluster composition. This gives immediately interpretable results.

We therefore sought to benchmark PMD against the more traditional metrics such as X^2^ statistic, its -log10(p-value), the less frequently used X^2^ statistic derived bias corrected Cramer’s V, inverse Simpson’s index, and Shannon’s entropy, under different idealized or adverse situations to assess for all of these properties. In each case, we calculated these metrics for a simulation of integrating two different batches, with differing numbers of clusters, and differing degrees of overlap between batches.

Importantly, what it means to have a given percentage overlap in cluster composition can come in two different modes. The first mode, and our first simulation paradigm (**Supplemental Fig. 7a**), is by sharing different numbers of clusters. For example, with two batches of four equally abundant clusters each could share no clusters (eight clusters total, four in each batch). They could share one out of the four clusters that appear in each batch (25% similar composition, 75% non-overlapping composition), etc. The second mode by which two batches can have varying degrees of overlap in cluster composition, is by sharing all, or some clusters, but having different relative abundance of the more-shared or less-shared clusters (**Supplemental Fig. 8a**). In our implementation of this second paradigm, we allocated a single cluster to batch 1, and vary the relative abundance of this cluster in batch 2, while batch 2’s remaining clusters are unique to it. In the case of 0% overlap, 0% of the second batch’s cells were allocated to cluster #1 (the cluster that harbors all of the cells from batch 1), and 100% of batch 2’s cells are allocated to clusters number 2 to n with equal probability. In the case of 25% overlap, 25% of batch 2’s cells are allocated to the first cluster, while the remaining 75% are distributed evenly across the remaining batch 2 specific clusters. Similarly, in the case of 50% overlap, 50% of batch 2 is allocated to cluster #1, and the remaining 50% of cells are allocated evenly to clusters unique to batch 2, etc. Note that in this second simulation paradigm, when there is 100% overlap between batches 1 and 2, each cell from batch 2 has a 100% probability of being allocated to cluster #1, thus being equivalent to having only a single cluster across both batches.

We utilized both of these simulation paradigms (**Supplemental Fig. 7a,8a**) to benchmark PMD against the previously mentioned status quo metrics, to evaluate each of these metric’s adherence to the above listed four desirable traits of an ideal metric for comparing cluster composition after batch-correction, integration and unsupervised clustering.

For our first Monte Carlo simulation in this comparison, we created two fully equivalent batches, with equal abundance (1000 cells each), and equal probabilities of sampling each cluster. We further simulated equivalent batches that harbored differing number of clusters (still all equivalent across batch, such that there was no pattern of clusters across batch). As expected from our theoretical work, PMD was robust to changes in dimensionality of the input matrix; however, with increasing number of clusters, the X^2^ statistic rose linearly the number of clusters in the batches (**Supplemental Fig. 7b**), thus demonstrating that the X^2^ statistic gives linear background signal when batches are equivalent, increasing with the number of clusters in each batch. Similarly, Cramer’s V showed a slight trend upwards. This is likely a phenomenon similar to the Raw PMD metric with a loss of power due to random sampling. Note that this pattern goes away when there are only a single cluster simulated (**Supplemental Fig. 8b**). Additionally, in the special case of one cluster, Cramer’s V cannot compute as it requires ≥2×2 contingency table. Interestingly, under the first simulation paradigm inverse Simpson’s index and Shannon’s entropy show negative trends with increasing number of clusters, but no trend under the second simulation paradigm (**Supplemental Fig. 7b,8b**).

Next, we performed a Monte Carlo simulation in which batches were completely orthogonal in their cluster composition, with no overlap; still simulating a variable number of clusters within each batch, but with no overlap (100% different), again equivalent in batch sizes. As expected, because of the invariance property, PMD was robust to different numbers of clusters, when there was no overlap in cluster composition between batches (**Supplemental Fig. 7c**). The X^2^ statistic however, lacked the invariance property, particularly when moving from 1 cluster per batch to 4 clusters per batch, but then plateaued. The -log10(p-value of the X^2^ test-of-independence however bottomed out with the lowest possible p-value due to floating capacity errors at -log10(p-value) = ∼308. Interestingly, Cramer’s V was negatively correlated with the number of clusters despite batches being completely different in each circumstance. The same pattern of results was observed under the second simulated paradigm (**Supplemental Fig. 8c**). Interestingly, similar to PMD, in this scenario inverse Simpson’s index and Shannon’s entropy, showed no relationship with the number of clusters, consistently yielding 1 or 0 respectively under both simulation paradigms (**Supplemental Fig. 7c,8c**).

We next assessed the effect of differences in batch sizes, where one batch was larger than the other. When cluster composition was the same, PMD was centered around zero, regardless of the number of clusters or differences in batch size (**Supplemental Fig. 7d**), whereas the X^2^ statistic was highly significant with one cluster, but dropped to non-significant with more than one cluster. The degree to which the one cluster results were significant by the X^2^ statistic was dependent on the dataset sizes as shown by the difference between dotted and solid lines (**Supplemental Fig. 7d**). Similarly to **Supplemental Fig. 7b** Cramer’s V showed some level of background above zero likely due to sampling error, as with Raw PMD. Inverse Simpson’s index and Shannon’s entropy showed different slopes dependent on the differences in dataset size, and Shannon’s entropy showed a global downward trend with increasing number of clusters.

Under the second simulation paradigm, where the number of clusters was only expanded in the second batch, this simulation was equivalent to having only a single cluster in total that held all cells from both batches. Consistent with the prior simulation, PMD was 0 (indicating no difference), while the X^2^ statistic and its corresponding p-values indicated highly significant differences by batch in a manner dependent on batch sizes (**Supplemental Fig. 8d**). Again, in the special case of a single cluster present, Cramer’s V fails to compute. Inverse Simpson’s index differed by size of the datasets, but under this simulation scenario, Shannon’s entropy was invariant with differing number of clusters or batch size (**Supplemental Fig. 8d**).

We next assessed a scenario similar to the above (**Supplemental Fig. 7e,8e**), but now with no overlap in cluster composition. PMD, inverse Simpson’s index, and Shannon’s entropy gave consistent results with PMD=1 (completely different), inverse Simpson’s index=1, and Shannon’s entropy=0, regardless of input batch sizes (**Supplemental Fig. 7e**). The X^2^ statistic however was fully dependent on the size of the input datasets. This indicates that differences in cell-filtering through variable QC pipelines, or other sources will directly impact the ability to use the X^2^ statistic or its -log10(p-value) for comparisons. Cramer’s V was dependent on both the sizes of the datasets and the number of clusters, despite the composition of clusters across batches being completely different in all cases. Furthermore, the invariance property of PMD and robustness to input dataset sizes demonstrate that it can be used, even to compare completely different dataset integrations, such as brain-heart concordance, relative to intestine-PMBC concordance. The same results were seen under the second simulation paradigm (**Supplemental Fig. 8e**).

#### Quantifying the linearity of PMD, χ^2^ statistic, and X^2^ statistic -log10(p-value), and Cramer’s V across the spectrum of similarity

We sought to quantify the linearity of these metrics, at intermediate levels of batch similarity, rather than only at the extreme poles (completely different and identical cluster composition). Linearity is important because it will enable the satisfaction of many parametric frequentist statistics assumptions without the need for additional transformations. For example, that the difference between 0 and 0.5 be equivalent to the difference between 0.5 and 1.0.

To test the property of linearity across the spectrum of similarity, we first created simulations in which batches were of equal size, with differing levels of overlap as determined by the presence or absence of shared clusters (**Supplemental Fig. 7a**). PMD gave consistent results, in which the PMD readout matched the simulated percentage of clusters that appeared in a batch specific manner (**Supplemental Fig. 7f**). For example, when 25% of clusters appeared in a batch specific manner, PMD was centered around 0.25, etc (**Supplemental Fig. 7f**, upper left panel). This concordance manifested in full linearity across the spectrum of overlap under this simulation paradigm for PMD (**Supplemental Fig. 7f**, lower left panel). Interestingly, this pattern held true for the X^2^ statistic as well (**Supplemental Fig. 7f**, middle-left panels). However, the X^2^ -log10(p-value) was non-linear, and was inversely proportional to the number of clusters (**Supplemental Fig. 7f**, middle panels). Similarly, Cramer’s V was non-linear across the spectrum of similarity (**Supplemental Fig. 7f**, middle-right panels). Inverse Simpson’s index and Shannon’s entropy were both non-linear and varied with number of clusters (**Supplemental Fig. 7f**; right panels). Importantly, however, many of these patterns changed under our second simulation paradigm.

Using the second simulation paradigm in which the percent overlap was determined by the relative abundance of a single potentially shared cluster, we again found that PMD was robust to the number of clusters and linear across the spectrum of batch similarity (**Supplemental Fig. 8f**, left panels). The X^2^ statistic however showed an exponential pattern across the spectrum of similarity that was consistent regardless of the number of clusters (**Supplemental Fig. 8f**, middle-left panels) and the -log10(p-values) were slightly non-linear as well, in a manner dependent on the number of clusters (**Supplemental Fig. 8f**, middle-left lower panel). Under these circumstances, Cramer’s V was largely linear (unlike the first simulation paradigm) (**Supplemental Fig. 8f**, middle-right lower panel), however, as previously mentioned, the special case of one cluster fails to compute. Inverse Simpson’s index and Shannon’s entropy both collapse towards one and zero respectively as the number of clusters increased, independently of the actual batch similarity (**Supplemental Fig. 8f**, right top panels). This resulted in very unstable estimates of similarity when compared to the actual similarity (**Supplemental Fig. 8f**, right bottom panels).

Next, we simulated a range of batch similarity, with identical batch sizes, but testing the effect of power under two scenarios – integrating two batches with 5,000 cells each (solid lines), compared to integrating two batches with 1,000 cells each (dashed lines). PMD was linear across the spectrum of similarity and robust to different levels of power under both simulation paradigms (**Supplemental Fig. 7g**, left panels). Interestingly, the X^2^ statistic was again linear within the two simulated batch sizes (two batches, 5,000 cells each and two batches 1,000 cells each), however, the slope of these lines was different, in which the simulated integrations with a larger number of cells had a greater slope compared to the smaller simulations (**Supplemental Fig. 7g**, middle-left panel). With the X^2^ test-of-independence -log10(p-value), we observed floating point errors, as the p-value approaches zero, that resulted in non-linearity with dataset similarity in the larger dataset (**Supplemental Fig. 7g**, middle-left panel). Cramer’s V was robust to dataset sizes, but was non-linear across the spectrum of similarity (**Supplemental Fig. 7g**, middle-right panel). Inverse Simpson’s index and Shannon’s entropy were again non-linear (**Supplemental Fig. 7g**, middle-right panels).

Interestingly, under the second simulation paradigm, the X^2^ statistic was exponential in relation to batch similarity, instead of being linear when batches as with the first simulation paradigm (**Supplemental Fig. 8g**, lower middle panel). Cramer’s V was robust to dataset size and linear, because batch sizes were identical, but failed under the special case of one cluster (**Supplemental Fig. 8g**, middle-right panels). Similarly to the first simulation paradigm, PMD, was again robust to dataset size, number of clusters, and linear across the spectrum of similarity (**Supplemental Fig. 8g**, left panels), while inverse Simpson’s index and Shannon’s again entropy dropped to one and zero respectively, when ground truth difference was intermediate (**Supplemental Fig. 8g**, right top panels), resulting in highly variable and non-linear estimates (**Supplemental Fig. 8g**, right bottom panels).

Lastly, we performed the same simulations as in (**ED Fig. 7g,8g**, but with datasets that were of different sizes within each integration (solid lines: batch1:1,000 cells, batch2:5,000 cells; dashed lines: batch1:500 cells, batch2:1,000 cells). PMD was again robust to differences in batch size, differences in power (as indicated by overlapping solid and dashed lines), was unaffected by the number of clusters, and operated linearly with batch similarity under both simulation paradigms, (**Supplemental Fig. 7h,8h**, left panels). The X^2^ statistic and its -log10(p-value) operated linearly only under the first simulation paradigm, but in a manner that was dependent on the number of clusters and power, while again under the second simulation paradigm, the X^2^ statistic and its -log10(p-value) were non-linear with batch similarity, dependent on the number of cells, the difference in batch size, and in the case of -log10(p-value), decreased with increasing number of clusters (**Supplemental Fig. 7h,8h**, middle-left panels). Under the first simulation paradigm, Cramer’s V was non-linear with percent of clusters that were unique to a given batch, but was robust to different dataset sizes (**Supplemental Fig. 7h**, middle-right panels). Under the second simulation paradigm, Cramer’s V was both non-linear and dependent on dataset sizes under this simulation paradigm (**Supplemental Fig. 8h**, middle-right panels). Surprisingly, the pattern of non-linearity in Cramer’s V was different under the two simulation paradigms (**Supplemental Fig. 7h,8h**, middle-right-lower panels). Inverse Simpson’s index and Shannon’s entropy followed the same patterns of collapse towards one and zero for intermediate similarity dependent on the number of clusters resulting in highly variable estimates (**Supplemental Fig. 7h,8h** right panels).

Overall, these results demonstrate that PMD is 1) invariant to the number of clusters, 2) robust to the size of input datasets, 3) robust to differences between batch sizes, and 4) operates linearly across the spectrum of similarity in batches when simulated as presence/absence of shared clusters across batches (first simulation paradigm) or relative abundance of a shared cluster (second simulation paradigm). The X^2^ statistic, its -log10(p-value), bias corrected Cramer’s V, inverse Simpson’s index, and Shannon’s entropy did not consistently share these properties robustly to all simulated circumstances.

**Supplemental Fig. 7:**
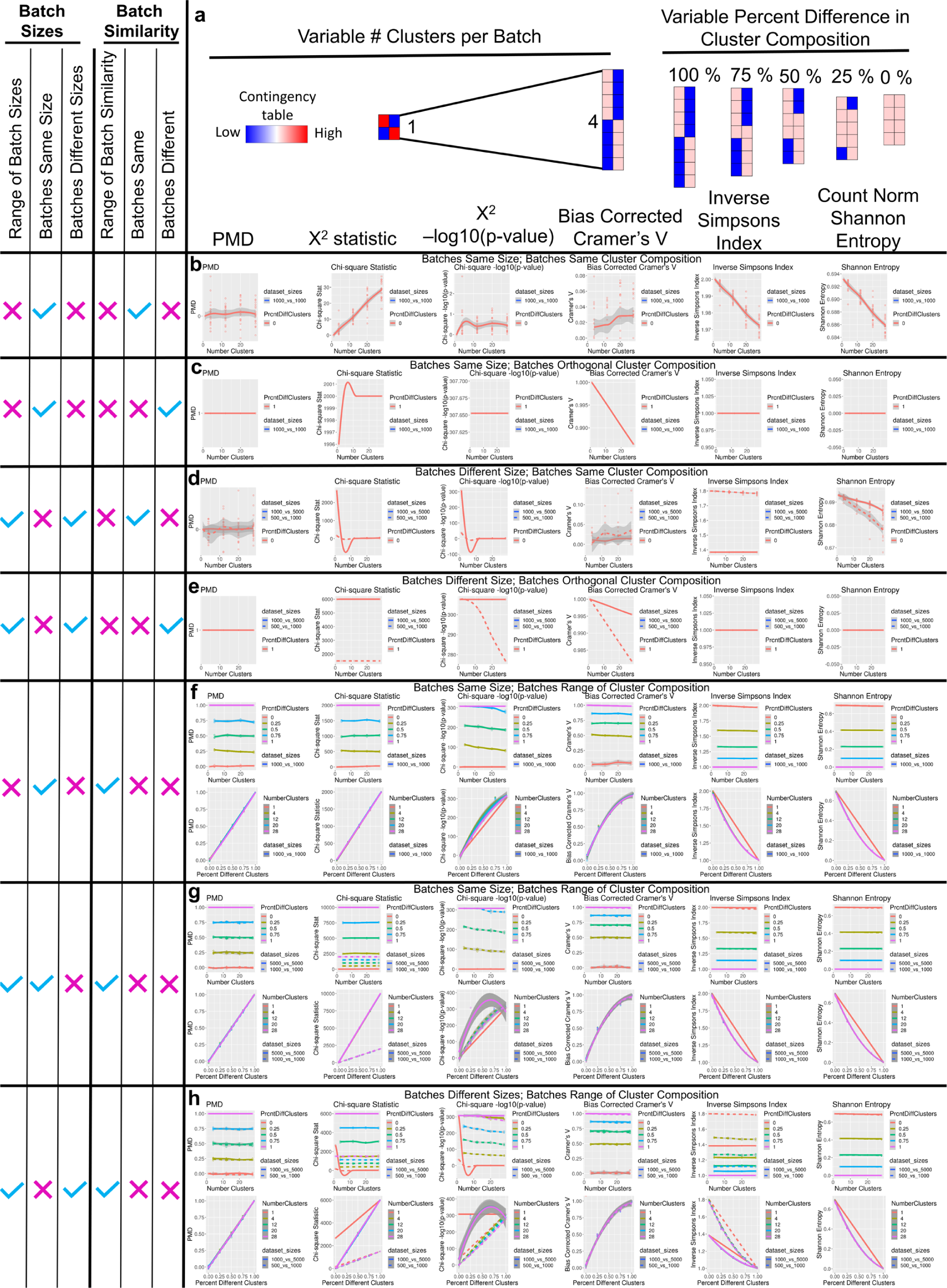
Extended Data Figure 9: Characteristics of PMD, X^2^ test-of-independence statistic, X^2^ -log10(p-value), Cramer’s V, inverse Simpsons index, and count normalized Shannon’s entropy. **a**, Schematic of the first simulation paradigm to vary batch similarity. Batches were comprised of the same number of clusters, but varied by the number of clusters that were batch specific clusters. **b,** PMD, X^2^, X^2^’s -log10(p-value), bias corrected Cramer’s V, inverse Simpson’s index, and Shannon’s entropy (y-axes) for the scenario in which batches have varying number of clusters (x-axis), batches contain the same number of cells, and batches share the same cluster composition. **c**, Batches are fully orthogonal, sharing no overlap in cluster composition, but batches contain the same number of cells with varying number of clusters (x-axis). **d**, Batches are identical, but with different batch sizes (indicated by solid & dashed lines). Note that Cramer’s V does not compute due to small contingency table size. **e**, Batches are different sizes (solid and dashed lines), and orthogonal cluster composition. **f-h**, (top row) Panels depicting the relationship between the simulated number of clusters (x-axis), and the corresponding metric (y-axis). Color indicates the ground-truth batch overlap. (Bottom row) x-axis is the ground-truth percent differences between batches, while colors indicate the number of clusters. In all panels, solid or dotted lines correspond to the indicated dataset sizes. **f**, Batches are the same size, over a range of cluster compositions. **g**, Similar to **f**, Batches are the same size, over a range of similarities, but now simulated with two different batch sizes (B1 & B2:1000 or 5000 cells each; solid or dotted lines). **h**, Batch sizes are each different within integration tasks, with variable scales of difference shown by solid or dotted lines.

**Supplemental Fig 8:**
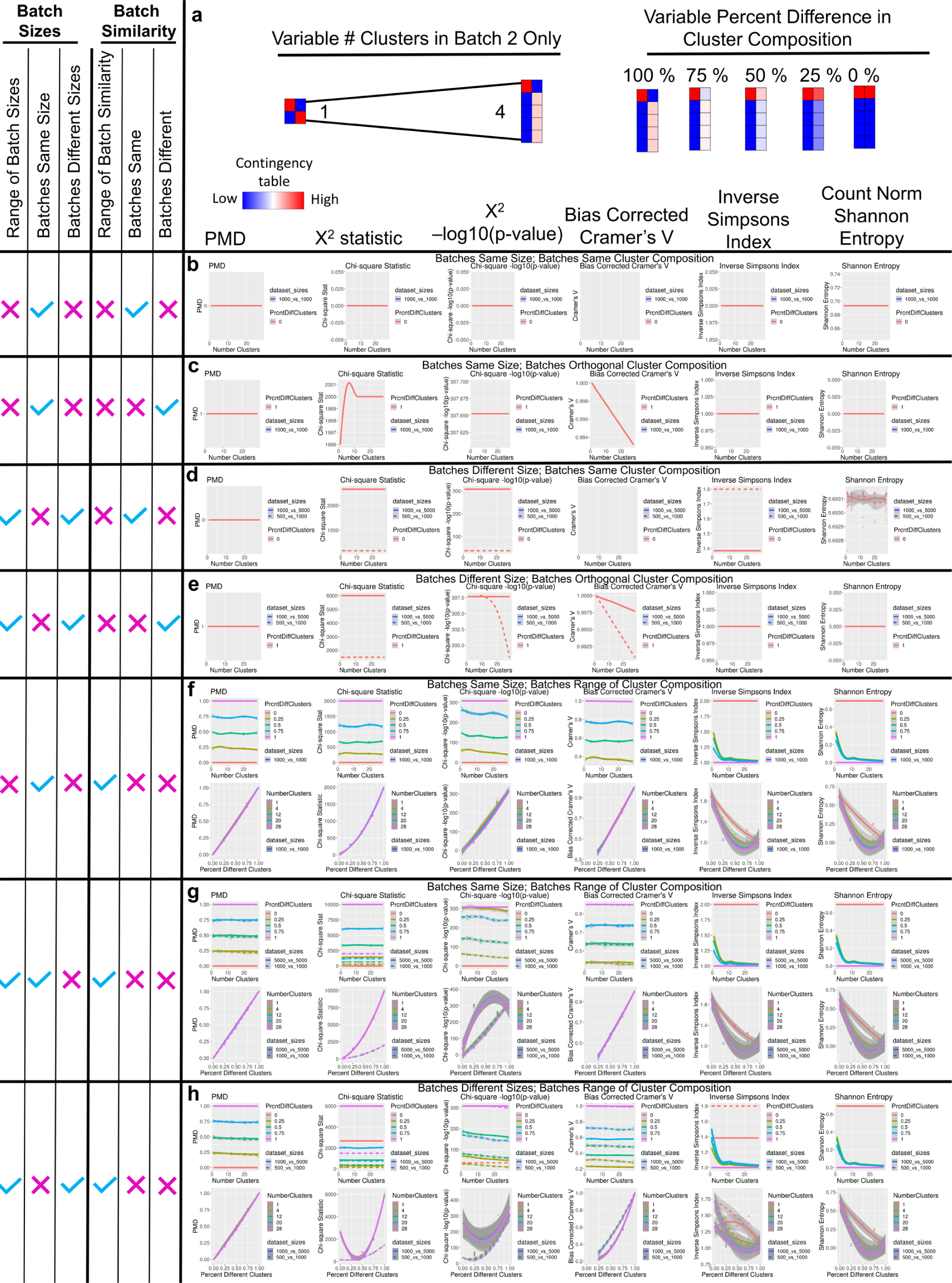
Benchmarking PMD against status quo metrics with altered cluster composition by relative abundance. **a**, The second simulation paradigm schematic for benchmarking performance with changing cluster relative abundance, rather than presence/absence of shared clusters as in **Supplemental Fig. 7. b-h**, See legend for **Supplemental Fig. 7**.

### Supplemental Note-5: Simulations of mixed expression programs

We simulated a mixture-model of negative binomial distributed gene expression profiles,^50, 72^ to test the effect of batch confounded gene-expression program mixtures, where batches differ in sequencing depth and are variably confounded with biology (See **Methods** for details).

We simulated six cell-types, with a cell-state optionally added, some of which appear in a batch confounded manner (**Fig. 6a**; **Supplemental Fig. 9**). Overall, we simulated 8 different scenarios in which cell-states were added in different arrangements across batches, in combinations with varying levels of clusters appearing in both batches, or in a batch specific manner; this totaled 32 simulation scenarios, each simulated in triplicate (96 batch integration tasks total) (**Supplemental Fig. 9**). To analyze these simulations, we employed 9 different quantitative metrics including those that measure global cluster accuracy after correction, the ability of algorithms to successfully remove the depth aspect of the batch effect, and the efficacy of algorithms in maintaining multiple sources of biologic variation, including when confounded by batch (**Box 1f**; **ED Table 4**).

**Supplemental Figure 9:**
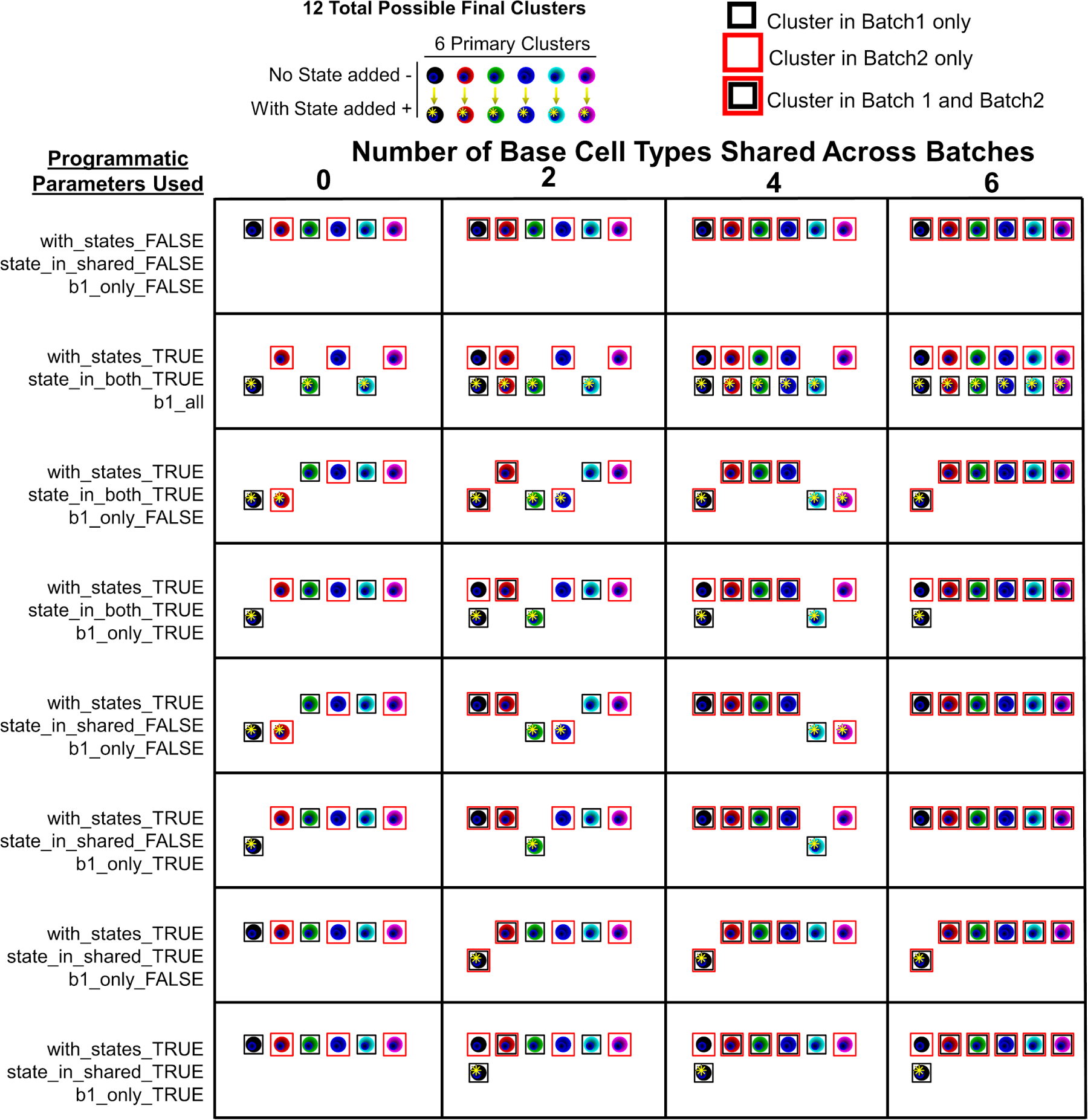
All simulated scenarios. We simulated 8 different parameter combinations (rows), with differing degrees of overlap in base cell-type presence in both or a single batch (columns). In all combinations, this generated 32 simulation combinations that were repeated in triplicate for 96 different scRNAseq batch-correction and integration tasks; note that some of these scenarios converged to similar schematics. The noted simulation parameters are also reported in **ED Table 4**.

In general, Class-1 algorithms tended to provide more accurate results using measures of PMD_observed_ distance to PMD_ground-truth_ (**Supplemental Fig. 10a-f**) and using cluster purity, and relative mutual information. Liger however was a Class-2 algorithm outlier. It appeared to have a, threshold of ground-truth PMD (PMD≈.25-.30) at which point its behavior changed; when batches were more different than this threshold, they would not be aligned (resultant “corrected” PMD≈1), explaining its periodically different behavior compared to other Class-2 algorithms (**Supplemental Fig. 10h**).

We next sought to directly answer the question: do integration methods inappropriately erase batch confounded cell-state information? To this end, we included simulations that had a base cell-type (with no cell-state added) in batch-1, while in batch-2, we simulated the same base cell-type, but with an added cell-state. With clusters identified as the mixture of all expression programs (cell-lineage, cell-type, cell-state), these two clusters should be identified as separate. We therefore quantified the purity of these populations, to measure if clusters are inappropriately merged relative to the ground-truth. Indeed, Class-1 normalization approaches showed high purity in this context, while Class-2+LDM approaches did not, indicating that normalization approaches preserve cell-state differences across batch (F=205.9, *P*=2.49e-78 main effects; *P*<1e-7 in all post-hoc comparisons; 1-way ANOVA/TukeyHSD; **Fig. 6b**). Similarly, if the largest source of variation (arbitrarily called cell-type variation) was different, yet shared the smaller source of variation (cell-state), these populations were often also merged by class-2 algorithms (F=73.95, *P*=8.31e-38 main effects; *P*<1e-7 in all post-hoc comparisons; 1-way ANOVA/TukeyHSD; **Fig. 6c**).

**Supplemental Figure 10:**
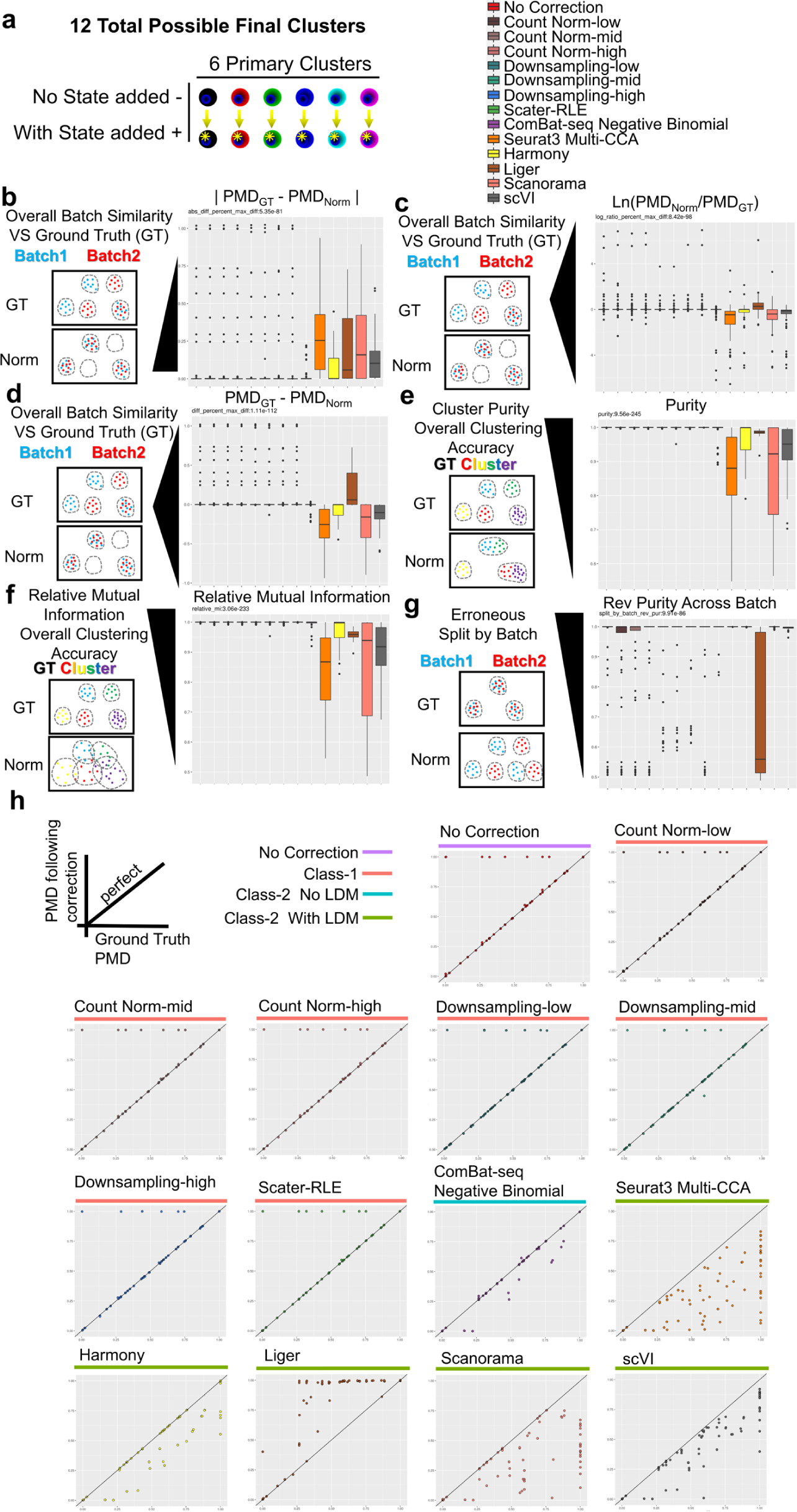
General accuracy of methods and efficacy in batch effect in simulations. **a**, Twelve possible clusters were used in simulations; they were generated from six primary clusters, with a cell-state optionally added (**Supplemental Fig. 6**). **b-g,** Boxplots and schematics illustrating the possible failure-mode are shown for six metrics. Boxplots are the typical display of median, with box around second-third quartiles, whiskers to the first and fourth quartiles, with outliers defined as 1.5x the inner quartile range. **b**, The absolute difference in ground-truth (GT) observed PMD indicates how different the dataset similarity is after batch-correction compared to the known ground-truth. **c**, Similar to **b**, but the log-ratio PMD ln(PMD_observed_ / PMD_ground-truth_). Higher indicates that an algorithm tends to leave residual batch effects (batches appear more different than they are in ground-truth). Lower indicates that an algorithm tends to make batches more similar than they are in the ground-truth. **d**, Difference between the ground-truth and observed PMD (PMD_ground-truth_ - PMD_observed_); interpretation is the same as **c**. **e**, Purity measures the quality of clustering results, penalizing when separate ground-truth clusters are erroneously merged together. **f**, Mutual information quantifies overall clustering accuracy using ground-truth known cluster identities as reference. **g**, Reverse purity quantifies cluster splitting; when subset for clusters that only appear in both batches, this metric penalizes cluster splitting specifically attributable to the batch effect. **h**, Scatterplots showing ground truth PMD (x-axis) against the observed PMD following application of the noted correction method.

### Supplemental Note-6: Analysis to determine if droplets positive for CD19 and mural cell markers were actually multiplets or multi-lineage contaminated droplets

Given that human brain samples are obtained at variable time intervals post-mortem, cell-death related signal, and cell-fragments/-debris may be present. In mice however, samples are perfused and rapidly harvested, reducing cell death, fragments and the potential for ambient RNA. We sought to determine whether the candidate CD19+/mural-like droplets show characteristics of being doublets or harboring across lineage cell fragments. Indeed, in all datasets, CD19-mural-cell-marker co-positive droplets exhibited both a higher number of observed genes and total counts (**Supplemental Fig. 11a,b**), metrics previously shown as indicative of doublets^54^. Moreover, the transcriptomes of these candidate cells were more correlated with a 50:50 transcriptome mixture model of B-cells and “pure” mural cells, compared to the mural-droplets that were negative for *Cd19* expression; this 50-50 mixture was also supported by the mean squared error (MSE) as the best fitting model. Note that these correlations and MSE calculations were performed holding out the markers that were used to identify those populations, thus A) preventing data leakage and self-fulfilling prophecy, and B) confirming that the B-like properties of the CD19+ mural-like droplets were *globally* more similar to B-cell transcriptomes rather than only being positive for CD19.

To directly measure the relative contribution of the “pure” mural and B-cell-like droplet transcriptomes to explain the candidate CD19/mural population, we fit a linear mixed model of the average log(count+1) transcriptomes of the reference “pure B-cell” (negative for mural markers) and “pure mural cell” transcriptomes. In the mouse dataset, this model clearly shows that, the candidate CD19/mural-cells fit best with a ∼70% contribution from B-cells and 30% contribution from mural cell transcriptomes. Our methodologic positive control, fitting “pure” mural cells (as a held-out population not used for reference construction), worked as expected with ∼0% contribution from B-cells and ∼100% contribution from mural cells in the mouse dataset (**Supplemental Fig. 11d**).

Notably however, when we performed this same analysis on the Zhong 2018 human dataset originally used to identify the putative CD19+/mural-cells, we observed similar coefficients between the candidate populations as well as when fitting a held out selection of the “pure” population of mural cells. A possible reason for this could be that that even the “pure” populations of mural cells (CD19 negative, B-cell marker negative, mural marker positive) were themselves also mixed in origin (**Supplemental Fig. 11d**). Fittingly, when we examined marker genes for other cell-types, we found that essentially all observed droplets were contaminated with marker genes across many lineages, indicating the presence of highly contaminated droplets in the original dataset (**Supplemental Fig. 11e**).

We further investigated the pathways of genes that were significantly correlated with CD19. Interestingly, using the full dataset, CD19 was correlated with genes enriched for both lymphocytes/immune system related pathways (-log10(p)=12.72; **ED Table 5a**) and vasculature/tube morphogenesis pathways (-log10(p)=12.97,9.07; **ED Table 5a**); however, when “pure” B-cells were removed from the dataset (Zong 2018^73^), CD19 *remained* correlated with the “immune system” (-log10(p)=8.05; **ED Table 5a**), indicating that from a global perspective, CD19 transcripts *continued* to track with the co-occurrence of other immune-related transcripts, fitting with a model of cell-fragment contaminations. When the candidate CD19+/mural-droplets were removed, the CD19 correlation with vascular and tube-morphogenesis was no longer significant, switching the CD19 correlations primarily to those correlated with immune related pathways, similar to those found with CD79A/B correlations (**ED Table 5b**), further supporting the hypothesis that this small population was indeed a mixture population.

Lastly, examination of a recently released human brain vasculature atlas (Garcia 2022) showed only a single droplet that was CD19+/mural-marker co-positive droplet (**Supplemental Fig. 11a-e**)^43^.

**Supplemental Figure 11:**
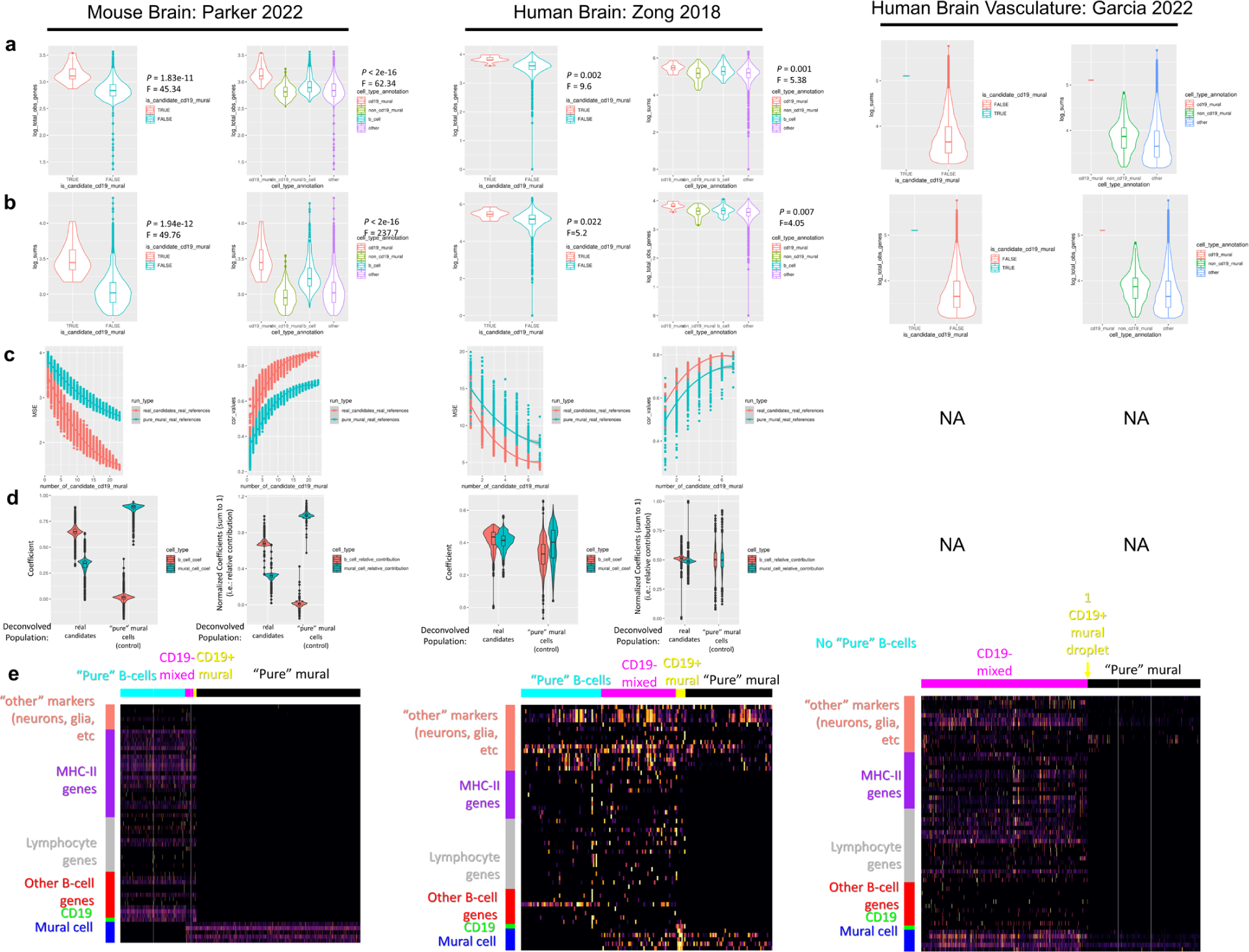
Droplets positive for CD19 and mural cell markers in the Zong 2018 dataset are consistent with cell-type mixtures rather than a novel population of mural cells. **a**, Violin plots showing the total number of observed genes in the “candidate” CD19+/mural-cell-marker+ double positive droplets and either all other cells, or the relevant possible mixture sources (B-cells, mural cells, and others). **b**, Similar to (**a**), violin plots showing the total number of counts rather than number of observed genes. **c**, Assessment of mean squared error (MSE) and Spearman correlation values determining how well a 50:50 mixture model of the average “pure” B-cell and “pure” mural cell transcriptomes fit with either the real candidates or a subset of “pure” mural cells (which were not included in the reference mixture model) shows that a 50:50 mixture model of the full transcriptome fits the data best (with the exclusion of CD19 and other markers used to identify these populations, to prevent data leakage), supporting that the candidate CD19+/mural cells are a global transcriptome mixture of mural and B-cells as opposed to a population of “pure” mural cells. Note where there were no “pure” B-cell-like droplets identified making a mixed model impossible, indicated by “NA.” **d**, Violin plots showing raw and normalized coefficients fitting a linear mixed model to “pure” B- and mural-cell references. Candidate CD19+ mural cells and methodologic positive controls (“pure” mural droplets) were held out from references to prevent data leakage. In the Zong 2018 dataset, *all* populations were consistent with a transcriptome mixture, suggesting broad contaminations rather than strict doublets, and that even the reference “pure” populations were not strictly lineage-pure. **e**, Heatmaps of lineage marker genes in the Zong 2018 human dataset confirm that nearly all droplets are lineage contaminated, while the mouse dataset appeared comparatively non-mixed. This fits with the mixture model results (**d**), in which the mouse mixture model had clearly distinct “pure” references, where the reference “pure” mural cells fit 100% with other mural cells, and 0% with B-cells, whereas in the Zong 2018 dataset, all cells fit well with a mixture model due to the transcriptomes of each lineage already being mixed. The human brain vasculature atlas did not contain a convincing CD19+/mural-droplet population (1 droplet).

